# Distinct regulatory networks control toxin gene expression in elapid and viperid snakes

**DOI:** 10.1101/2023.02.13.528252

**Authors:** Cassandra M. Modahl, Summer Xia Han, Jory van Thiel, Candida Vaz, Nathan L. Dunstan, Seth Frietze, Timothy N. W. Jackson, Stephen P. Mackessy, R. Manjunatha Kini

## Abstract

Venom systems are ideal models to study genetic regulatory mechanisms that underpin evolutionary novelty. Snake venom glands are thought to share a common origin, but there are major distinctions between venom toxins from the medically significant snake families Elapidae and Viperidae, and toxin gene regulation in elapids is largely unexplored. Here, we used high-throughput RNA-sequencing to profile gene expression and microRNAs between active (milked) and resting (unmilked) venom glands in an elapid (Eastern Brown Snake, *Pseudonaja textilis*), in addition to comparative genomics, to identify *cis*- and *trans*- acting regulation of venom production in an elapid in comparison to viperids (*Crotalus viridis* and *C. tigris*). Although there is conservation in high-level mechanistic pathways regulating venom production, there are histone methylation, transcription factor, and microRNA regulatory differences between these two snake families. Histone methyltransferases (KMT2A, KMT2C and KMT2D) and transcription factor (TF) specificity protein 1 (Sp1) were highly upregulated in the milked elapid venom gland, whereas nuclear factor I (NFI) TFs were upregulated after viperid venom milking. Sp1 and NFI *cis*-regulatory elements were common to toxin gene promoter regions, but many unique elements were also present between elapid and viperid toxins. microRNA profiles were distinctive between milked and unmilked venom glands for both snake families, and microRNAs were predicted to target different toxin transcripts. Our comparative transcriptomic and genomic analyses between toxin genes and isoforms in elapid and viperid snakes suggests independent toxin evolution between these two snake families, demonstrating multiple toxin genes and regulatory mechanisms converged to underpin a highly venomous phenotype.

## Introduction

Snake venom glands are highly specialized vertebrate secretory tissues, exhibiting elevated, tissue-specific expression of 50-100 or more toxins. Based on structural similarity to cognate proteins involved in physiological processes, snake toxins are hypothesized to have primarily evolved by gene duplication of nontoxic genes (Vogel et al. 1984; Kochva et al. 1993; Hite et al. 1994; Joseph et al. 1999; Miwa et al. 1999; Rao and Kini 2002; Tamiya and Fujimi 2006) and subsequent neofunctionalization through accelerated evolution of their exons (Deshimaru et al. 1996; Nobuhisa et al. 1996; Chang et al. 1997; Ohno et al. 1998). Therefore, snake venom glands provide a unique opportunity to study epigenetic and genetic mechanisms that regulate tissue-specific expression and gene neofunctionalization in multi-locus gene families. Furthermore, as venom glands are a rich source of diverse bioactive proteins applicable to bioprospecting and antivenom development, understanding how toxins are produced would also benefit large-scale production *in vitro* (Post et al. 2020).

Snakebite annually leads to the death or disability of over 500,000 victims (Gutiérrez et al. 2017) and medically significant species are predominantly from the families Elapidae and Viperidae. Venoms from these two families cause debilitating effects and appear to share a common evolutionary origin, which is supported by anatomical and developmental evidence (Vonk et al. 2008), and phylogenetic analysis of toxins suggests venom evolved once, at the base of the advanced snake radiation (Fry and Wüster 2004). Although elapids and viperids both have tubular fangs positioned anterior on the maxilla, elapid and viperid delivery systems and venom components are distinct in several ways. Elapids have a less mobile maxillary bone, and relatively smaller fangs and venom gland lumen, while viperids have a highly kinetic maxillary, long fangs and a wide lumen (Kardong 1982; Kochva 1987). Different external adductor muscles are also used to compress the venom gland in elapids and viperids (Jackson 2003), as well as likely convergent evolution of accessory glands (Kerkkamp et al. 2015). After venom is expelled, venom gland cells do not exhibit any marked size differences in elapids (Kochva et al. 1982; Lachumanan et al. 1999); while in viperids, secretory epithelial cells dramatically increase in size from cuboidal to columnar with proliferation of the rough endoplasmic reticulum (ER) (Oron and Bdolah 1973; Mackessy 1991). Further, elapid venoms are largely dominated by non-enzymatic three-finger toxins (3FTxs), and enzymatic group I phospholipases (PLA_2_s) of pancreatic origin (Mackessy 2010; Tasoulis and Isbister 2017), while viperid venoms have primarily enzymatic toxins, including snake venom metalloproteinases (SVMPs), serine proteinases (SVSPs), and group II PLA_2_s with structural features similar to non-pancreatic inflammatory PLA_2_s (Heinrikson et al. 1977; Seilhamer et al. 1989). Chromosomal locations of dominant toxins also vary between these two snake families, with most located on macrochromosomes in elapids (Suryamohan et al. 2020; Zhang et al. 2022) and microchromosomes in viperid snakes (Shibata et al. 2018; Schield et al. 2019; Margres et al. 2021).

In both elapids and viperids, toxin gene transcription and translation are upregulated when the venom gland is emptied by venom extraction or “milking” (Rotenberg et al. 1971; Paine et al. 1992; Lachumanan et al. 1999; Currier et al. 2012). Studies investigating mechanisms regulating the expression of elapid toxins have focused on a small number of genes. These studies have identified *cis*-regulatory elements (CRE) that regulate the expression of genes encoding 3FTx and group I PLA_2_ expression in the Javan Spitting Cobra (*Naja sputatrix*) (Jeyaseelan et al. 2000; Ma et al. 2001; Ma et al. 2002), the pseutarin C catalytic subunit in the Eastern Brown Snake (*Pseudonaja textilis*) (Kwong et al. 2009), and silencer AG-rich motifs in the first intron of the pseutarin C catalytic subunit gene (Han et al. 2016). Investigations of viperids have found α1− and β−adrenoceptor signaling as a mechanism activating transcription factors (TFs) NF-kB and AP-1 to initiate toxin synthesis in the Jararaca (*Bothrops jararaca*) (Yamanouye et al. 1997; Luna et al. 2009), tissue-specific TF ESE-3 upregulated group II PLA_2_s genes in the Habu (*Protobothrops flavoviridis*) (Nakamura et al. 2014), and CREs for binding GRHL-and NFI-family TFs were enriched within SVMP, SVSP, and group II PLA_2_ gene promoters in the Prairie and Tiger rattlesnakes (*Crotalus viridis* and *C. tigris*, respectively) (Schield et al. 2019; Margres et al. 2021; Perry et al. 2022), in addition to co-opted TFs from the Unfolded Protein Response (UPR) pathway (Perry et al. 2022). Current evidence suggests that for the viperids *C. viridis* and *C. tigris,* toxin gene expression is regulated by chromatin structure, TFs, and gene methylation levels (Schield et al. 2019; Margres et al. 2021). Here, we compare active (milked) and resting (unmilked) venom glands of *P. textilis* to provide the first evidence of multiple toxin gene regulatory networks in an elapid snake.

*Pseudonaja textilis* is the second most lethal terrestrial venomous snake in the world, based upon murine models of toxicity (Broad et al. 1979), and is responsible for the majority of snakebite deaths in eastern Australia (Sutherland 1992; White 2009). Venom gland transcriptomics and venom proteomics have been used to profile *P. textilis* venom composition (Birrell et al. 2006; Viala et al. 2015; Reeks et al. 2016), and studies have documented geographic, seasonal, ontogenetic, and individual venom variation (Williams and White 1992; Flight et al. 2006; Skejić and Hodgson 2013; Skejić et al. 2015; Jackson et al. 2016; McCleary et al. 2016; van Thiel et al. 2023). Using high-throughput transcriptomics, we evaluated *P. textilis* mRNAs and microRNAs (miRNAs) in a Milked Venom Gland (MVG) and Unmilked Venom Gland (UVG) from an individual snake, eliminating contributing factors involved in venom variation, to identify mechanisms regulating toxin expression. We identified higher-level venom synthesis activation pathways common to both *P. textilis* and viperid venom glands, but differences in *cis-* and *trans-*acting regulation of toxin expression. Further, posttranscriptional miRNA regulation was not conserved between venom glands from the two snake families. Therefore, distinct gene regulatory networks produce elapid and viperid venom phenotypes, and thus, our results suggest an independent origin of toxin genes and their associated regulatory mechanisms in elapid and viperid snakes.

## Results

### Toxin genes exhibit high and variable expression in the *Pseudonaja textilis* venom gland

From a *P. textilis* individual, the left venom gland was milked to stimulate venom production while the right venom gland remained unmilked as an ‘unstimulated’ control. Four days later (96 hours post venom milking, hpvm), both glands were collected, RNA and miRNA isolated and sequenced, and the genome (EBS10Xv2-PRI) used as a reference to profile gene expression between glands. Evaluations of *P. textilis* toxin transcript diversity and expression were also completed with transcripts from *de novo* assembled transcriptomes for each gland (Supplemental Table 1). Fold-changes in transcript expression between the MVG versus UVG were determined with GFOLD (Feng et al. 2012) (Supplemental Table 2). A total of 7,062 transcripts were upregulated and 8,038 transcripts downregulated in the MVG.

For both the MVG and UVG, toxins were among the most highly expressed transcripts and the same toxin superfamilies made up similar proportions of overall toxin expression (Supplemental Figure 1A-B). From a combined *de novo* assembly of each gland (Supplemental Table 1), 28,569 transcripts were expressed (>1 Transcript Per Million, TPM), and 52 full-length toxins from 18 toxin superfamilies were identified. Toxin superfamilies that were expressed, in order of total abundance in the MVG, include fourteen 3FTxs (78% of all toxin transcripts), three group I PLA_2_s (13%), four Kunitz-type serine proteinase inhibitors (KUNs, 3%), 13 snake venom C-type lectins (Snaclecs, 2%) and the prothrombin activator pseutarin C (venom coagulation factors V and X, 2%) (Supplemental Figure 1A). Low abundance toxins, making up 1% or less of reads, were two natriuretic peptides (NPs,), one cysteine-rich secretory protein (CRISP), three cystatins (CYSs), two SVMPs, one hyaluronidase (HYAL), one nerve growth factor (NGF), one waprin (WAP), one 5’-nucleotidase (5’NUC), one cobra venom factor (CVF), one L-amino acid oxidase (LAAO), one acetylcholinesterase (AChE) and one vespryn (VES). All transcript isoforms in a superfamily were not expressed equally; for the two superfamilies with the highest expression levels, the fourteen 3FTx isoforms ranged from 236,773 TPM to 10 TPM, and the three PLA_2_ isoforms ranged from 100,141 TPM to 1,846 TPM.

This *de novo* assembled venom gland transcriptome is currently the most comprehensive to date for *P. textilis*, having detected all previous reported toxin transcripts along with full-length transcripts for low abundance CYSs, HYAL, NGF, WAP, 5’NUC, CVF, LAAO, AChE, and VES toxins (Supplemental Table 3). Three 3FTxs (3FTx_6, 7, and 8), one CRISP (CRISP_1), one KUN (KUN_1), and one PLA_2_ (PLA2_1) were identical to previously identified sequences (Supplemental Figure 2A-D). PLA_2_s and short-chain 3FTxs exhibited the greatest sequence variation (as low as 43% and 48% amino acid sequence identity, respectively) between isoforms. There was an absence of transcripts sequences for textilotoxin, a pentameric PLA_2_ complex unique to *P. textilis* venom (Tyler et al. 1987a). Pseutarin C, a toxin composed of two subunits homologous to the mammalian coagulation factor Xa-Va complex (Rao and Kini 2002), were highly conserved, 99% and 95% identical to the published sequences for *P. textilis* venom coagulation factors V and X, respectively (Rao et al. 2003; Rao et al. 2004) (Supplemental Figure 2E-F), and >84% identical to venom coagulation factor homologs in *Oxyuranus microlepidotus* and *O. scutellatus* venom.

### Upregulation of toxin gene expression after venom milking is lower in an elapid compared to viperids

To compare changes in toxin gene expression in the MVG of an elapid to that of vipers, we retrieved available RNA-seq libraries from the National Center for Biotechnology Information (NCBI) for MVGs and UVGs of *C. viridis* at 0 and 72 hpvm (Schield et al. 2019), and *C. tigris* at 0, 72 and 96 hpvm (Margres et al. 2021). To have a comparable time course of 0 to 96 hpvm for all species, we milked venom from a *C. viridis* individual of the same northeastern Colorado locality as snakes used by Schield *et al*. (2019), collected both venom glands at 96 hpvm, and performed RNA-seq on these tissues. Annotated transcriptomes from genome assemblies (UTA_CroVir_3.0 and ASM1654583v1 for *C. viridis* and *C. tigris*, respectively) and myotoxin transcripts from a *de novo* assembled venom gland transcriptome for *C. viridis* (96 hpvm individual) (Supplemental Table 4) were used as references to determine toxin transcript abundances. Toxin transcripts were identified from transcriptomes from key word searches for each venom protein family, in addition to using previously annotated toxin genes (Supplemental Table 5 and 6 for *C. viridis* and *C. tigris*, respectively) (Schield et al. 2019; Margres et al. 2021). We reanalyzed fold-changes in expression of transcripts between MVGs and UVGs at each time point with GFOLD (Feng et al. 2012) (Supplemental Table 7), allowing for comparisons to the *P. textilis* dataset.

Toxin expression in the venom gland of *C. viridis* from northeastern Colorado was predominately myotoxins (75% of total toxin transcripts), SVSPs (11%), PLA_2_s (7%), SVMPs (6%), and minor toxins (< 1%) (Supplemental Table 5). For *C. tigris,* PLA_2_s were the highest expressed (56% of all toxin transcripts), followed by SVSPs (21%), bradykinin-potentiating peptides (BPPs; 12%), vascular endothelial growth factor (VEGF; 5%), SVMPs (4%), and all other minor toxins (1% or less) (Supplemental Table 6). A myotoxin from *C. viridis* had the highest expression level (520,544 TPM at 96 hpvm) of any toxin from either the two rattlesnake venom glands or from the elapid *P. textilis* (Figure 1). For all species, the highest expressed toxins (>50% of toxin transcripts) were the major components in each venom proteome (Figure 1), and are also the primary lethal or tissue damaging toxins in these venoms (Cameron and Tu 1978; Ho and Lee 1981; Su et al. 1983; Tyler et al. 1987b; Calvete et al. 2012; McCleary et al. 2016).

**Figure 1.**
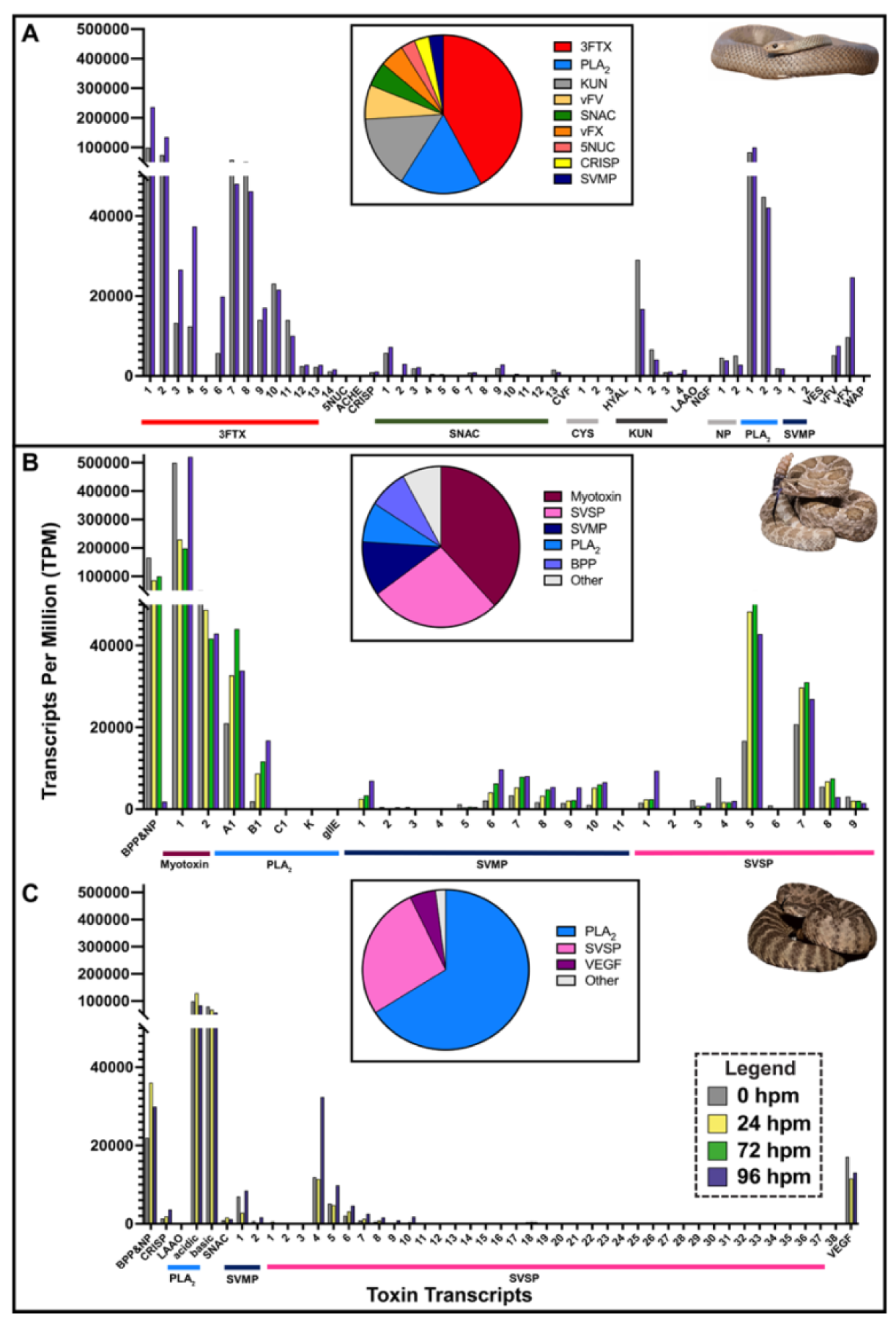
Toxin gene expression in an elapid (*Pseudonaja textilis*) and two viperids (*Crotalus viridis* and *C. tigris*) over a time course of 96 hours post venom gland milking (hpvm) in comparison to venom proteomes. Time points are shown for (A) *P. textilis* at 0 and 96 hpvm, (B) *C. viridis* at 0, 24, 72, and 96 hpvm, and (C) *C. tigris* at 0, 24, and 96 hpvm. Bar colors represent the time after venom milking. Pie charts (insets in respective panels) correspond to the venom proteomes of *P. textilis* (McCleary et al. 2016), *C. viridis* (Saviola et al. 2015) and *C. tigris* (Calvete et al. 2012). Toxin identifications are as follows: 3FTx = three-finger toxin, 5NUC = 5’-nucleotidase; AChE = acetylcholinesterase; BPP = Bradykinin-potentiating peptide; CRISP = cysteine-rich secretory protein; CVF = cobra venom factor; CYS = cystatin; HYAL = hyaluronidase; KUN = Kunitz serine proteinase inhibitor; LAAO = L-amino acid oxidase; NGF = nerve growth factor; NP = natriuretic peptide; PLA_2_ = phospholipase A_2_; SNAC = snake C-type lectin; SVMP = snake venom metalloproteinase; SVSP = snake venom serine protease; VEGF = vascular endothelial growth factor; VES = vespryn; vFV = venom factor V (pseutarin C non-catalytic subunit); vFX = venom factor X (pseutarin C catalytic subunit); WAP = waprin. Photo credits: *P. textilis,* Ákos Lumnitzer; *C. viridis*, Wolfgang Wüster; *C. tigris,* Ben Lowe.

For toxin expression in the *P. textilis* MVG, highly abundant toxins (3FTx_1, 2, 3, 4 and PLA_2__1) exhibited TPM increases (Figure 1A) with overall fold-changes of two-fold or less in comparison to the UVG. There were small decreases in expression (less than 30%) for 5’NUC, two 3FTxs (3FTx_8 and 10), two PLA_2_ (PLA2_2 and 3) and HYAL, and larger decreases (36-66%) for two 3FTxs (3FTx_7 and 11), AChE, two CYSs (CYS_1 and 2), two KUN (KUN_1 and 2), two NPs, four snaclecs (SNAC_5, 8, 11, and 13) and WAP. Snaclec transcript isoforms exhibited the greatest variability between the MVG and UVG; SNAC_2 had the overall highest increase (11.4-fold) and SNAC_5 had the greatest decrease (0.31-fold).

For toxin expression in the viperid MVG, SVMP_1 in *C. viridis* was the top upregulated transcript by 43-, 58-, and 127-fold at 24, 72, and 96 hpvm, respectively. However, there were not parallel increases in expression for all rattlesnake toxins over the time course. For *C. viridis,* the highest expressed PLA_2_ (PLA2_A1) peaked 72 hpvm and another PLA_2_ (PLA2_B1), the majority of SVMPs (7 out of 11) and myotoxin_1 peaked at 96 hpvm (Figure 1B). SVSP transcripts in *C. tigris* venom glands showed the greatest fold changes over the time course: SVSP_1 (XM_039325703.1) was upregulated 699-fold at 24 hpvm and SVSP_2 (XM_039360544.1) was the top upregulated toxin transcript (45-fold) at 96 hpvm.

### UPR, Notch signaling, and cholesterol homeostasis are enriched in MVGs, and different biological process regulated between an elapid and viperids

A Gene Set Enrichment Analysis (GSEA) (Mootha et al. 2003; Subramanian et al. 2005) was conducted using human gene orthologs for all non-toxin genes expressed in MVGs and UVGs. No gene sets were found to be significantly enriched in MVGs for any of the three species, but the lowest familywise error rate (p-value=0.50) was for UPR in *P. textilis*, Notch signaling in *C. viridis* (familywise error rate p-value=0.49) and cholesterol homeostasis in *C. tigris* (familywise error rate p-value=0.49), although these three biological processes were listed in enriched datasets of all three species with the exception of cholesterol homeostasis in *C. viridis* (Supplemental Table 8).

Focusing on genes with the greatest fold-change between the MVG and UVG from *P. textilis*, we selected all genes upregulated at least 10-fold (373) and downregulated to less than 0.10-fold (415) in the MVG. A gene ontology and network analysis of the upregulated gene set found the following overrepresented biological processes: chromatin and histone remodeling (GO:006325, GO:0016569, and GO:006338), regulation of transcription (GO:0006355, GO:0006357, and GO:0045944) and phospholipid biosynthesis (including phosphatidylinositol-3-phosphate signaling GO:0036092 and inositol phosphate metabolic process GO:0043647) (Figure 2A,C). Chromatin organization was found to be the only significantly upregulated biological process (p=0.0005; for details, see below). Downregulated genes were significantly enriched for proteins involved in striated muscle contraction and sarcomere assembly (p < 0.002; GO:0006936, GO:0014733, GO:0006937, GO:0090257, GO:0055002, and GO:0030239) (Figure 2B,C).

**Figure 2.**
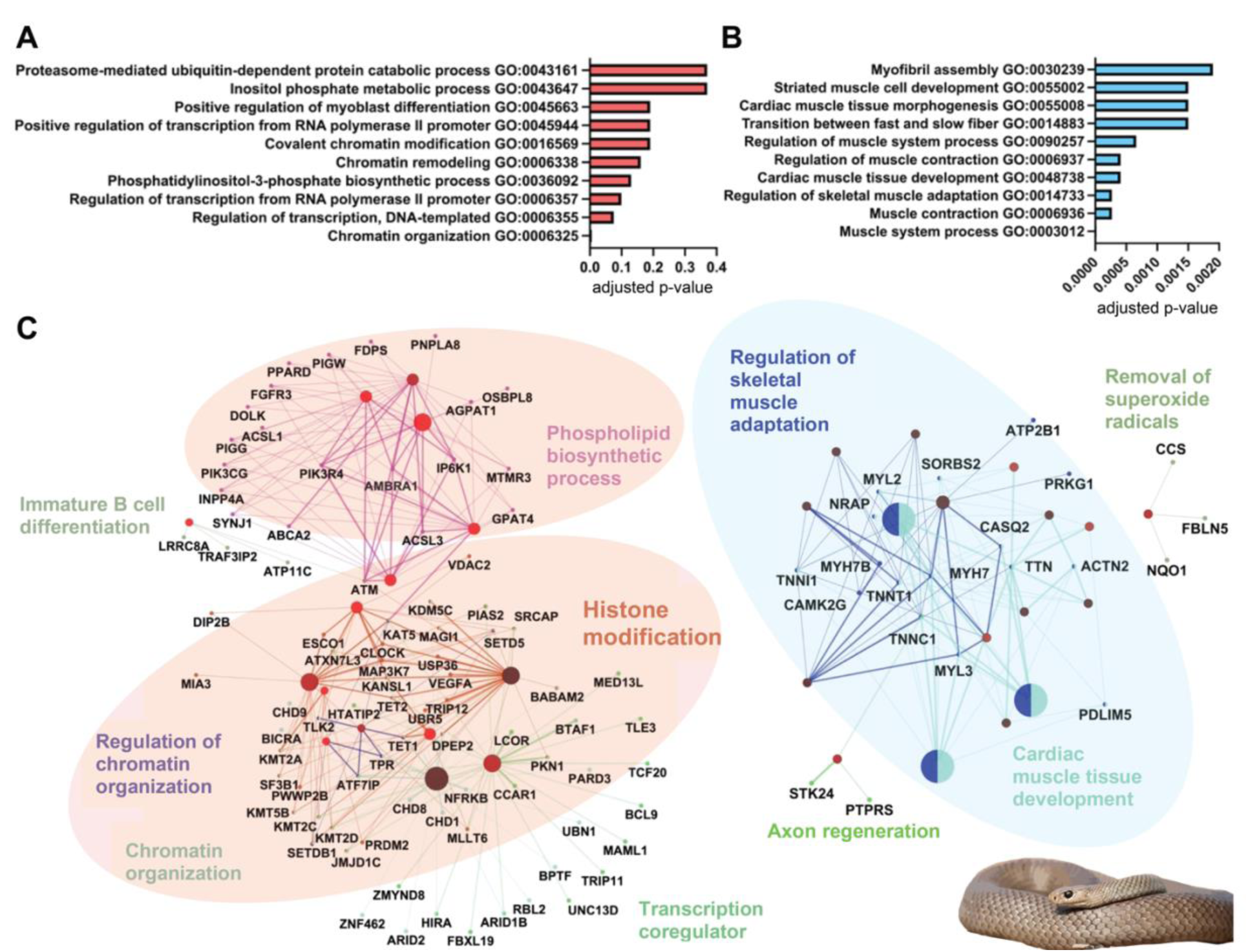
Enriched biological processes and associated networks for genes up-and downregulated in *Pseudonaja textilis* milked venom gland. The top ten biological processes are shown with their level of significance for genes (A) upregulated at least 10-fold and (B) downregulated to less than 0.10-fold 96 hpvm. Gene ontology analysis was completed using DAVID Bioinformatics Resources (Huang da et al. 2009b; Sherman et al. 2022) and Benjamini-Hochberg adjusted p-values were used for identifying levels of significance. For each up-and downregulated gene set, (C) gene networks and associated biological processes were generated using the ClueGo app plug-in (Bindea et al. 2009) in Cytoscape (Shannon et al. 2003) with *Homo sapiens* orthologs. Light red colored ovals highlight upregulated gene networks with the greatest number of related nodes, and the light-colored blue oval for those downregulated. Photo credit: *P. textilis*, Ákos Lumnitzer.

For the viperids *C. viridis* and *C. tigris*, 16 and 197 genes were upregulated, and 85 and 515 genes downregulated 96 hpvm, respectively, using the same thresholds as used for *P. textilis*. A gene ontology and network analysis of the upregulated gene set found the following overrepresented biological processes: transcription (GO:0045944, GO:0000122, and GO:006354), protein translation and transport (GO:0017148, GO0001822, and GO:0015031), and the UPR (GO:0030968) (Figure 3A, C). Positive regulation of transcription and negative regulation of translation were significant upregulated biological processes (p < 0.05); Figure 3A). Downregulated biological processes were complement activation (GO:0006958 and GO:0006956), immune response (GO:0006954, GO:0045087, and GO:0006955), cellular component organization (GO:0016043, GO:0071840, and GO:0051128) and metabolic processes (GO:0019219, GO:0051173, GO:0009893, and GO:0031325) (Figure 3B); all were significant (p < 0.008). A gene regulatory network analysis of the downregulated gene sets found the largest gene network related to negative mechanisms of regulating nucleobase-containing macromolecules, which included transcriptional repressors (Figure 3C).

**Figure 3.**
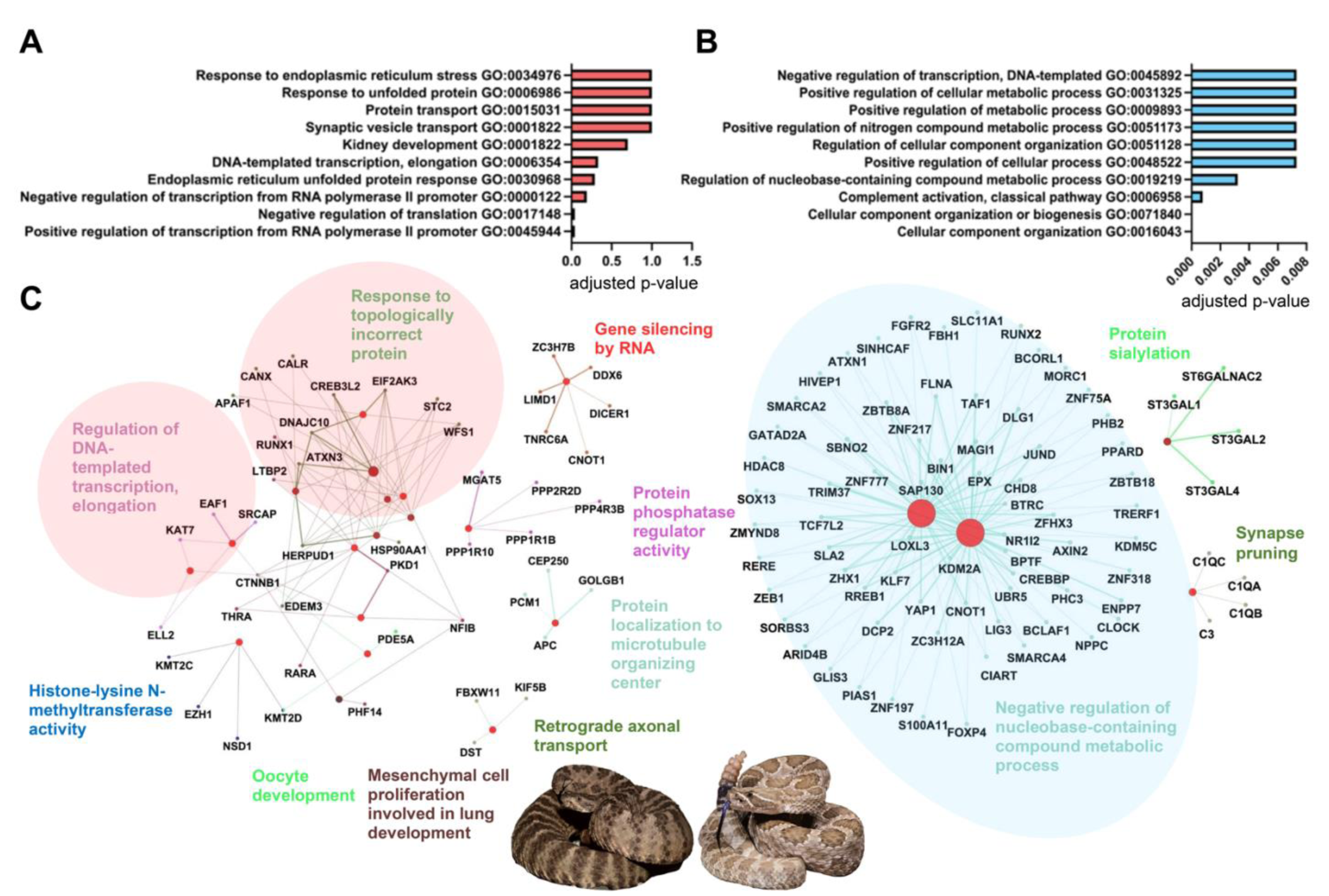
Enriched biological processes and associated networks for genes up-and downregulated after venom milking viperids *Crotalus viridis* and *C. tigris.* The top ten biological processes are shown with their level of significance for genes (A) upregulated at least 10-fold and (B) downregulated to less than 0.10-fold at 96 hpvm. Gene ontology analysis was completed using DAVID Bioinformatics Resources (Huang da et al. 2009b; Sherman et al. 2022) and Benjamini-Hochberg adjusted p-values were used for identifying levels of significance. For each up-and downregulated gene set, (C) gene networks and associated biological processes were generated using the ClueGo app plug-in (Bindea et al. 2009) in Cytoscape (Shannon et al. 2003) with *Homo sapiens* orthologs. Light red colored ovals highlight upregulated gene networks with the greatest number of related nodes, and the light-colored blue oval for those downregulated. Photo credits: *C. tigris,* Ben Lowe*; C. viridis,* Wolfgang Wüster.

### Chromatin remodelers and transcription factors are differentially regulated after venom miking between an elapid and viperids

Given the upregulation of genes involved in chromatin organization and histone modification in the *P. textilis* MVG (Figure 2A, C), we identified all chromatin modifiers upregulated at least 40-fold: Snf2 related CREB activator protein (*SRCAP*, 86-fold), jumonji domain containing 1C (*JMJD1C*, 43-fold), lysine methyltransferases 2A (*KMT2A,* 80-fold), KMT2C (*KMT2C,* 45-fold), KMT2D (*KMT2D*, 64-fold), and chromodomain-helicase-DNA-binding protein 8-like (*CHD8,* 52-fold) (Table 1 and Supplemental Table 9). *CHD8* was upregulated only 2-fold in the *C. viridis* MVG at 72 hpvm and *SRCAP, KMT2C* and *KMT2D* were upregulated 12-, 24- and 18-fold, respectively, in the *C. tigris* MVG at 96 hpvm. Chromatin modifiers were not seen as highly upregulated in viperid MVGs (Supplemental Table 10).

**Table 1.**
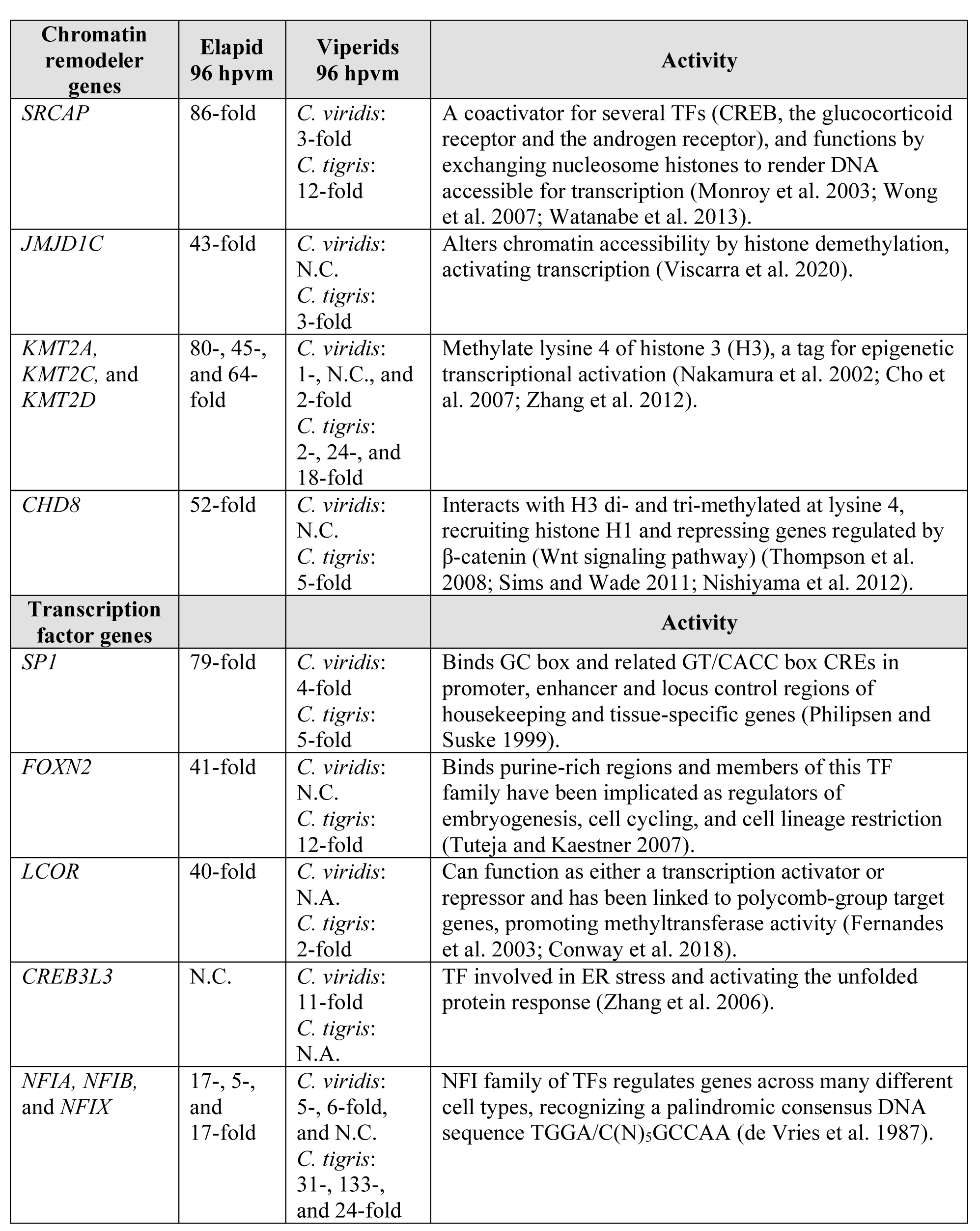
Gene expression and activities of upregulated chromatin remodeler genes and transcription factors in elapid and viperid milked venom glands. N.C. = No Change; N.A. = Not Annotated.

Although chromatin structure regulates the accessibility of gene regulatory elements, TFs play vital roles in the regulation of transcription. The TF with the greatest fold upregulation in the *P. textilis* MVG, and with a transcript variant X2 uniquely expressed in the MVG, was specificity protein 1 (*SP1*, 79-fold) (Table 1 and Supplemental Table 9). In the MVG of the rattlesnakes *C. viridis* and *C. tigris*, *SP1* was only slightly upregulated comparatively, 1.5-, 2- and 4-fold at 24, 72 and 96 hpvm in *C. viridis* and 14- and 5- fold at 24 and 96 hpvm, respectively, in *C. tigris.* In addition, Forkhead box N2 (*FOXN2*) and ligand-dependent corepressor (*LCOR*) were upregulated 41- and 40-fold, respectively, in the *P. textilis* MVG (Supplemental Table 9), but not to this extent in either viperid (Table 1). The TF with the greatest fold-change in the *C. viridis* MVG was cAMP-responsive element binding protein 3-like (*CREB3L3*), which was upregulated 14-, 12-, and 11-fold at 24, 72, and 96 hpvm, respectively. Nuclear factor I isoforms (*NFIA, NFIB,* and *NFIX*) were found to be the TFs with the greatest upregulation in *C. tigris*. *NFIA* was upregulated 31-fold at 96 hpvm, *NFIB* was upregulated 104-fold and 133-fold, and *NFIX* was upregulated 8- and 24- fold at 24 hpvm and 96 hpvm, respectively, in *C. tigris* (Table 1 and Supplemental Table 10). *NFIA* and *NFIB* were also found upregulated 5-fold and 6-fold, respectively, at 96 hpvm in *C. viridis* (Table 1).

### *Cis*-regulatory elements (CREs) and *trans-*factor upregulation vary between elapids and viperids

Using toxin gene promoter regions from elapids and viperids, we predicted CREs and evaluated the expression of corresponding *trans*-regulatory factors in MVGs. Promoter activities determined either by reporter gene chloramphenicol acetyl transferase or luciferase assays have identified the importance of CREs within the first 500 base pairs upstream of toxin genes (Jeyaseelan et al. 2000; Kwong et al. 2009). Due to variability in toxin gene expression, we evaluated regions upstream each toxin gene isoform independently to determine CREs potentially contributing to differential expression. For *P. textilis,* this included 341 bp for the 3FTx pseudonajatoxin b (AY027493) (Figure 4A), 684 bp for short-chain 3FTx (AF204969) (Figure 4B), and 714 bp for a non-conventional 3FTx (Figure 4C). Only one gene is present for pseudonajatoxin b (Gong et al. 2001), and five different short-chain 3FTx genes share the same 684 bp promoter sequence (Gong et al. 2000). Interestingly, among 3FTxs, although many Old World elapids have non-conventional 3FTxs in their venoms with a fifth disulfide bond in the first loop (Nirthanan et al. 2003), *P. textilis* did not express non-conventional 3FTxs in the venom gland, despite the presence of a non-conventional 3FTx gene in its genome (XP_026561523). We included this toxin gene in the analysis to determine if unique CREs were present for a non-expressed 3FTx.

**Figure 4.**
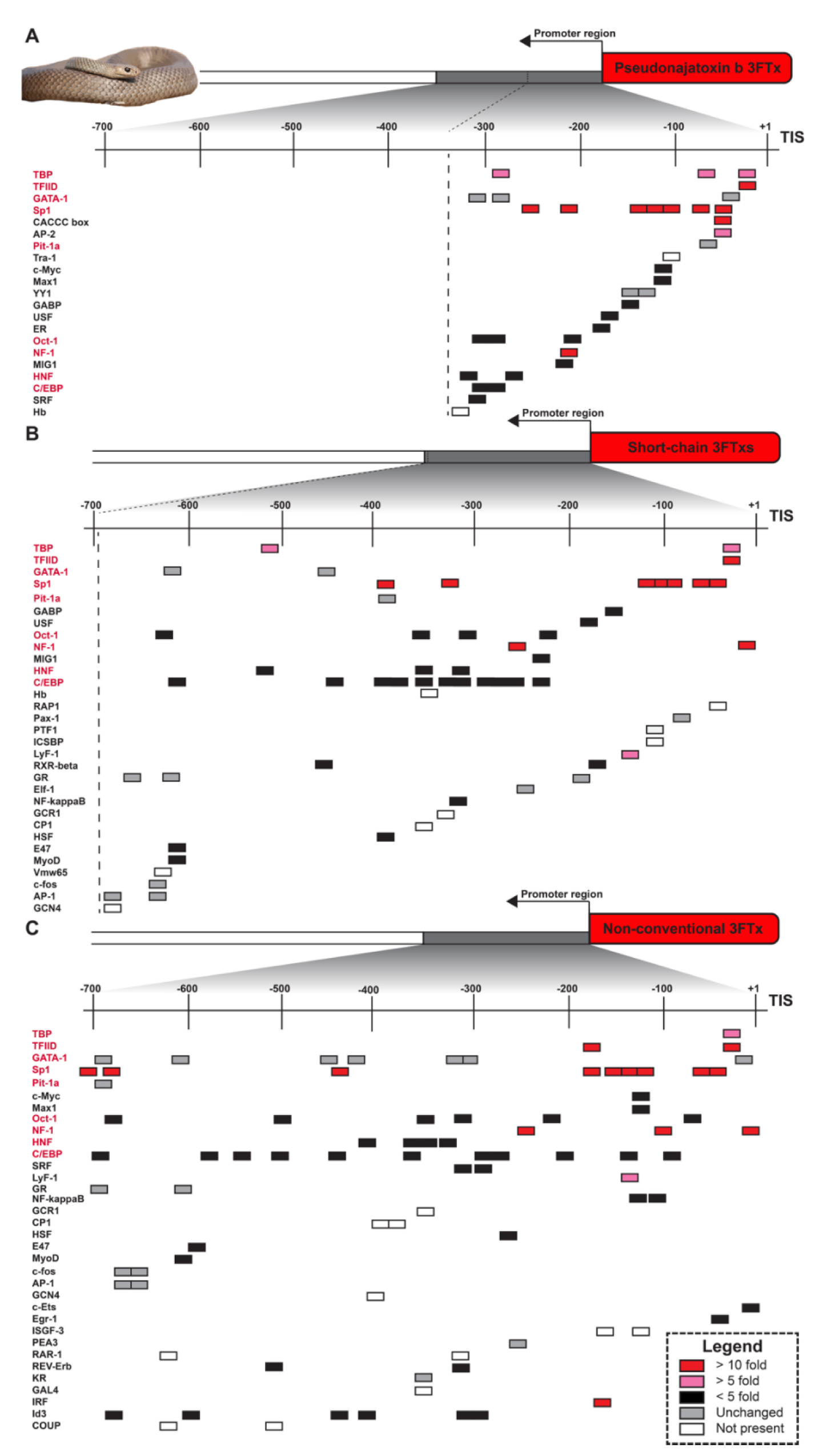
Predicted *cis-*regulatory elements in the promoter regions of *Pseudonaja textilis* three-finger toxins. Only the first 341 bp upstream from the Transcription Initiation Site (TIS) could be evaluated for (A) three-finger toxin (3FTx) pseudonajatoxin b (AY027493) and 684 bp upstream from the TIS for (B) short-chain 3FTxs (AF204969) due to limited sequence availability and fragmented genome assembly. *Pseudonaja textilis* did not express transcripts encoding non-conventional 3FTxs in the venom gland, despite the presence of a non-conventional 3FTx gene in its genome (XP_026561523). From the genome reference, 713 bp upstream from what would be the TIS for this (C) non-conventional 3FTx was evaluated. Fold-changes in expression levels are shown between the *P. textilis* milked venom gland and unmilked venom gland for *trans*-factors known to interact with predicted *cis-*regulatory elements (CREs). CRE predictions were completed with the online server AliBaba2.1 using the TRANSFAC 4.0 database. CREs shared across panels are bolded red. Photo credit: *P. textilis,* Ákos Lumnitzer.

Predicted CREs in promoter regions of the three 3FTx classes varied, likely due to nucleotide sequence diversity, as there was 86-88% conserved sequence in the first 341 bp; however, we did identify shared sites for TATA-box binding proteins (TBP) and TATA-box binding protein associated factors of the RNA polymerase II preinitiation complex (TFIID). Additionally, there were GATA-1, Sp1, Pit-1a, Oct-1, NFI, HNF, and c/EBP binding sites across all promoters, but these varied in number and location. All 3FTx gene promoter sequences did have multiple (six to nine) Sp1 binding sites (Figure 4). Of the predicted TFs binding to 3FTx gene promoter regions, Sp1/CACCC-box, NFI, and interferon regulatory factor (IRF) were found upregulated at least 10-fold in the *P. textilis* MVG. The IRF binding site was only present in the non-conventional 3FTx gene promoter sequence. We did not find any TFs that were downregulated to less than 0.10-fold, which was our threshold.

CREs upstream of PLA_2_ genes that are highly expressed in elapids (group I PLA_2_) and viperids (group II PLA_2_) were also evaluated (Figure 5). Elapid group I PLA_2_s are subdivided into group IA and IB, group IB is likely the ancestral PLA_2_ gene with the presence of the complete pancreatic loop (Fujimi et al. 2004). For *P. textilis,* group IB PLA_2_s are the most abundant and two group IB PLA_2_ genes have been found in the *P. textilis* genome with identical 385 bp sequence upstream from the transcription initiation site (TIS) (Figure 5A) (Armugam et al. 2004). For *Laticauda semifasciata* and *Naja sputatrix*, group IA PLA_2_s are the most abundant, and 706 bp and 367 bp upstream from TISs were evaluated, respectively (Figure 5B,C) (Jeyaseelan et al. 2000; Fujimi et al. 2004). Group II PLA_2_s with the highest expression levels in *C. viridis* and *C. tigris* venom glands were PLA2_A1 and PLA2_acidic (XM_039367474), respectively, and just over 700 bp of promoter regions were evaluated for each (Supplemental Table 5 and 6, Figure 5D,E). Regardless of group IA, IB or II, CREs for binding TBP, Sp1, c/EBP, USF, and NFI were present for all PLA_2_ genes. Multiple Sp1 binding sites (eight to 19) were observed clustered together within all PLA_2_ upstream gene regions. For the *P. textilis* group IB PLA_2_, TFs with CREs present and upregulated at least 10-fold in the MVG were Sp1/CACCC-box and NFI, the same two TFs as seen for 3FTxs. For group II PLA_2_s in the viperids, TFs with CREs and upregulated over 10-fold were NFI, retinoic acid receptor (RAR), upstream stimulatory factor 1 (USF1), and thyroid hormone (3,5,3’-triiodothyronine) receptor (T3R) (Figure 5D,E). Interestingly, although Sp1 was slightly upregulated (5-fold) in *C. tigris* 96 hpvm, this was not observed for *C. viridis*, but NFI was upregulated at least 5-fold in both viperids. TFs that were downregulated less than 0.10-fold did not have any predicted CREs.

**Figure 5.**
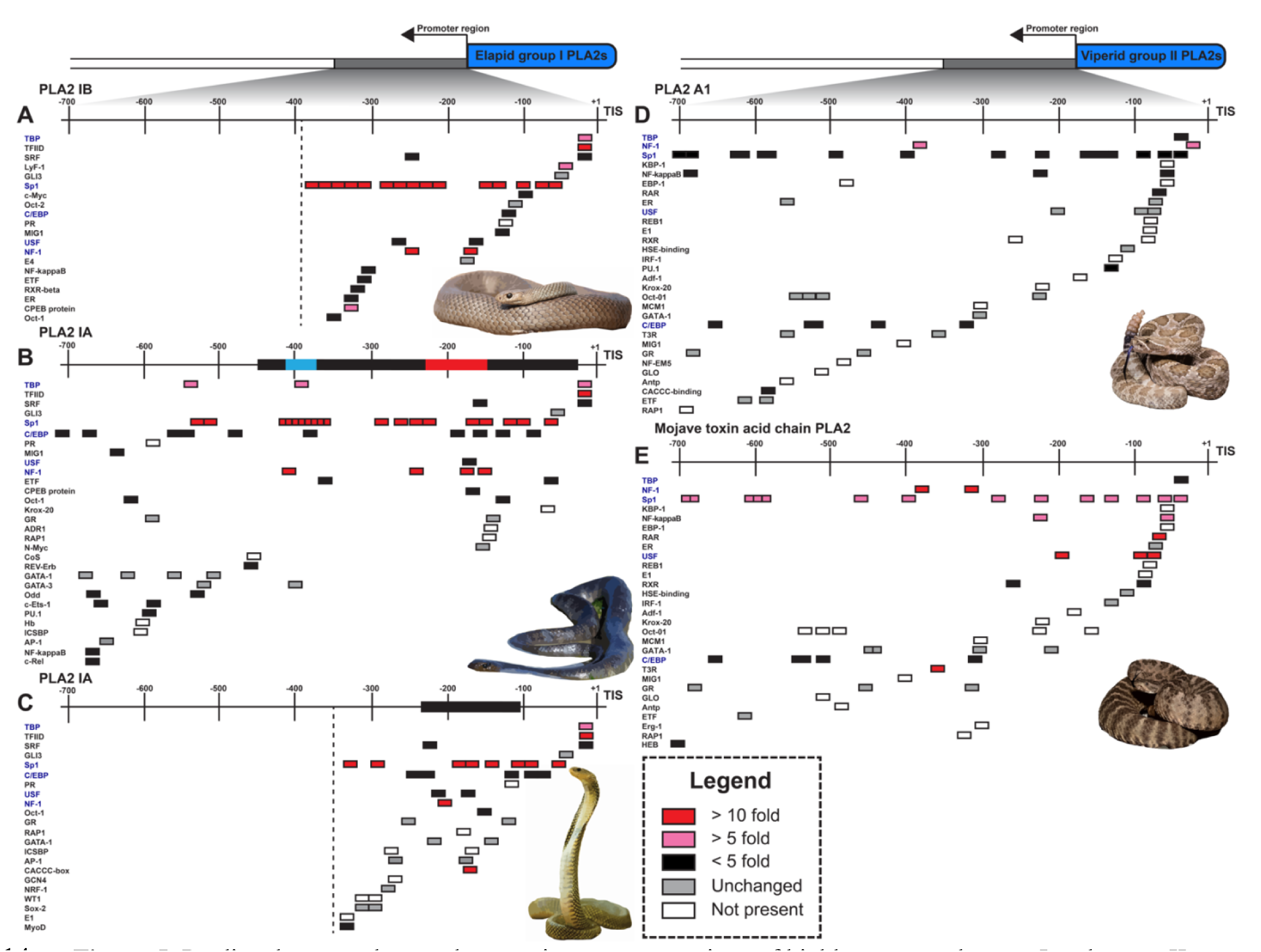
Predicted *cis-*regulatory elements in promoter regions of highly expressed group I and group II phospholipase A_2_s. Promoters regions upstream from Transcription Initiation Sites (TISs) are shown: 385 bp for (A) group IB PLA_2_ (AY027495) from *Pseudonaja textilis,* 706 bp for (B) group IA PLA_2_ (AB111958) from *Laticauda semifasciata*, 367 bp for (C) group IA PLA_2_ (AF101235) from *Naja sputatrix,* 702 bp for (D) group II PLA2_A1 from *Crotalus viridis,* and 705 bp for (E) group II PLA2_acidic (XM_039367474) from *C. tigris.* Fold-changes in expression levels are shown between milked and unmilked glands for *trans*-factors known to interact with the predicted *cis-*regulatory elements (CREs). Identified regulatory regions are bolded in promoter regions for AB111958 (Fujimi et al. 2004) and AF101235 (Jeyaseelan et al. 2000). This includes a region from -232 to -162 in AB111958 that was found to be responsible for an increase in promoter activity (bolded red) and a suppressor region from -410 to - 382 (bolded blue) (Fujimi et al. 2004). CRE predictions were completed with the online server AliBaba2.1 using the TRANSFAC 4.0 database. CREs shared across panels are bolded blue. Photo credits: *P. textilis*, Ákos Lumnitzer; *L. semifasciata*, Patrick Davis; *N. sputatrix,* Matej Dolinay; *C. viridis*, Wolfgang Wüster; *C. tigris*, Ben Lowe.

Additionally, we evaluated upstream regions that have been experimentally shown to regulate toxin gene promoter activity. Fujimi et al. (2004) found a 411 bp insertion sequence (-444 to -34) present in the highly expressed group IA PLA_2_s that was absent in the lowly expressed group IB for *L. semifasciata*. Luciferase activity assays from construct variations of this insertion identified a region from -232 to -162 that triggered elevated expression and a suppressor region from -410 to -382 (Fujimi et al. 2004). Jeyaseelan et al.(2000) used chloramphenicol acetyl transferase reporter gene assays and DNase 1 footprinting approaches with promoter constructs from a *N. sputatrix* group I PLA_2_ gene to identify a region from -116 to -233 that contained crucial CREs (Jeyaseelan et al. 2000). We found CREs with binding sites for Sp1 and NFI in all of these identified regulatory regions (Figure 5B, C). Previously, we identified a 271 bp insertion (-308 to -37) upstream of the gene for pseutarin C catalytic subunit (venom coagulation factor X) that differed from the endogenous coagulation factor X gene. We termed this segment *VERSE* (*Ve*nom *R*ecruitment/*S*witch *E*lement) (Reza et al. 2007). Within the *VERSE* core promoter there are two regions that upregulate the pseutarin C catalytic subunit (Up1 and Up2) and one that suppresses expression (Sup1) (Kwong et al. 2009). Here, we re-analyzed the TFs binding to these regulatory regions and found CREs for Sp1 in Up2 and c/EBPdelta in Sup1 (Supplemental Figure 3). c/EBPdelta was downregulated 0.82-fold in the *P. textilis* MVG.

### Venom gland miRNA profiles and targets are distinct between elapid and viperid snakes, and after venom milking

MiRNAs are known to post-transcriptionally regulate over 60% of mammalian genes (Friedman et al. 2009). We sequenced small RNA-seq libraries from the *P. textilis* MVG and UVG, and MVGs from *C. viridis* (96 hpvm), to examine miRNA expression and regulation in snake venom glands. miRNAs were identified by lengths of 18-23 bp, alignment to genomes, and the presence of a transcribed hairpin structure, predicted by miRDeep2 (Friedländer et al. 2012). A total of 366 miRNAs (308 non-redundant mature sequences) in the MVG and 375 miRNAs (299 non-redundant mature sequences) in the UVG were identified in *P. textilis* (Supplemental Table 11), and 501 miRNAs (420 non-redundant mature sequences) were identified in the MVGs from *C. viridis* (Supplemental Table 12). Approximately 50% of miRNAs from the *P. textilis* MVG and UVG were common (Supplemental Figure 4A), but only 18% of miRNAs found in *C. viridis* were also present in *P. textilis* (Supplemental Figure 4B). The most abundant miRNAs in *P. textilis* venom glands were found to be *miR-148a-3p* with 359,436 Counts Per Million reads (CPM) in the MVG and *miR-10c* with 210,834 CPM in the UVG (Supplemental Figure 5A, B). *miR-375* had been found to be the most abundant miRNA in the venom gland of the king cobra (*Ophiophagus hannah*) (Vonk et al. 2013) (Supplemental Figure 5C), but this was not the case for *P. textilis* where it was ranked 13^th^ in the MVG and 7^th^ in the UVG. Interestingly, *miR-375* was not present in the venom gland of *C. viridis.* For *C. viridis, miR-21-5p* was the most abundant (157,080 CPM), followed by *miR-148a-3p* (140,288 CPM) (Supplemental Figure 5D), both of which were present but varied in expression between the elapid and viperid venom glands (Table 2).

**Table 2.** miRNA differences between elapid and viperid milked venom glands.

| miRNA | Elapid<br>96 hpvm | Viperid<br>96 hpvm | Activity |
| --- | --- | --- | --- |
| *miR-375* | Present | Absent | Multiple transcript targets predicted that relate to ubiquitin regulation, stress response, collagen biosynthesis, translation, endocytosis and trafficking (Supplemental Table 14) (The UniProt 2021). |
| *miR-148a-3p* | 359,436 CPM | 140,288 CPM | Predicted to target C-type lectin-like toxin transcript in *P. textilis* (SNAC_5), no predicted targets in viperids (Supplemental Table 13 and Supplemental Table 15). |
| *miR-21-5p* | 22,809 CPM | 157,080 CPM | No predicted mRNA targets in the elapid but predicted to target the N-alpha-acetyltransferase 25 auxiliary subunit in viperid (Supplemental Table 15). |
| *miR-215-5p* | Absent | Present | Predicted to target a SVMP transcript (SVMP_4) in *C. viridis* MVGs (Supplemental Table 13). |
| Top ten miRNAs | Toxin transcripts targeted:<br>Snaclecs, group I PLA<sub>2</sub>s, 3FTxs and pseutarin C. | Toxin transcripts targeted:<br>SVMPs | Different toxin transcript targets were predicted from elapid and viperid venom glands (Supplemental Table 13). |
| miRNAs over 100 CPM | mRNA transcripts targeted:<br>Intracellular transport, catabolic processes, metabolic processes, organelle organization, and ER to Golgi mediated transport. | mRNA transcripts targeted:<br>Regulation of mRNA processing, catabolic processes, negative regulation of cytoskeleton organization, steroid hormone receptor signal, stress granule assembly, transport, and ER stress. | Transcripts for proteins involved in different biological processes were targeted in elapid and viperid MVGs (Figure 6). |
| *Pte-miR-1* | Present in the *P. textilis* MVG | Absent | Predicted to target 510 mRNAs expressed in the *P. textilis* MVG. These included transcripts for proteins involved in ER to Golgi vesicle transport, ubiquitin-dependent ERAD pathway and proteasome-mediated ubiquitin-dependent protein catabolic process (Supplemental Figure 6). |

miRNAs post-transcriptionally repress mRNAs by base pair complementary binding (Bartel 2004), usually within the 3’ untranslated region (3’ UTR) of target mRNAs, but interactions can also be within the 5’ UTR or coding sequences (O’Brien et al. 2018). Predicted miRNA:mRNA interactions are largely controlled by seed complementarity and duplex free energy (Doench and Sharp 2004). To explore potential miRNA regulation of toxin transcripts (Durban et al. 2013), toxin transcript targets were predicted for the top ten most abundant miRNAs. This was done with the miRanda position-weighted local alignment algorithm using a criteria of -19 kcal/mol or less free energy pairing between miRNA:mRNA (Enright et al. 2003; John et al. 2004), as used previously to identify toxin transcripts targeted by venom gland miRNAs (Durban et al. 2013; Durban et al. 2018). In the *P. textilis* MVG, miRNAs targeted transcripts for seven C-type lectin-like toxins (SNAC_2, 3, 5, 7, 8, 9, and 12), all three PLA_2_s, and the pseutarin C catalytic subunit. In the *P. textilis* UVG, the same toxin transcripts were targeted with an additional six short-chain 3FTxs (3FTx_7, 8, 10, 11, 12 and 13) and the pseutarin C non-catalytic subunit (Table 2). In the *C. viridis* venom gland, two snake venom metalloprotease transcripts (SVMP_4 and 9) were targeted (Supplemental Table 13), although not all toxin transcripts were evaluated (only those listed in Supplemental Table 5). *miR-215-5p,* which targeted the SVMP_4 transcript, was found uniquely expressed in the *C. viridis* venom gland (Table 2 and Supplemental Figure 5D).

Next, we predicted mRNAs targeted by abundant miRNAs (over 100 CPM) using all transcript annotations from the *P. textilis* and *C. viridis* genomes to identify the most likely regulated mRNAs and pathways. To reduce false positives, we used a strict criteria of -30 kcal/mol or less free energy pairing between miRNA:mRNA and only evaluated transcripts that were co-expressed (at least over 10 TPM) in the venom glands. A total of 750, 264, and 244 transcripts in the *P. textilis* MVG, *P. textilis* UVG and *C. viridis* MVGs, respectively, met these criteria with predicted miRNA binding sites (Supplemental Table 14 and 15). Gene regulatory network analyses of miRNA targets identified a greater number of biological processes targeted for downregulation in MVGs, especially for *P. textilis* (Figure 6). Biological processes targeted in *P. textilis* MVG included intracellular transport (GO:0006886 and GO:0006888) and several metabolic and catabolic related processes (GO:0043170, GO:1901565, GO:1901564, among others) (Figure 6A). With the exception of ER to Golgi vesicle transport, different biological processes were targeted for downregulation in the *P. textilis* UVG (Figure 6B). Targeted processes in *C. viridis* MVGs shared interestingly little overlap to those of *P. textilis* and included regulation of mRNA processing (GO:0006397), response to endoplasmic reticulum stress (GO:0034976), and ribonucleoside diphosphate metabolic process (GO:0009185), among others (Figure 6C and Table 2).

**Figure 6.**
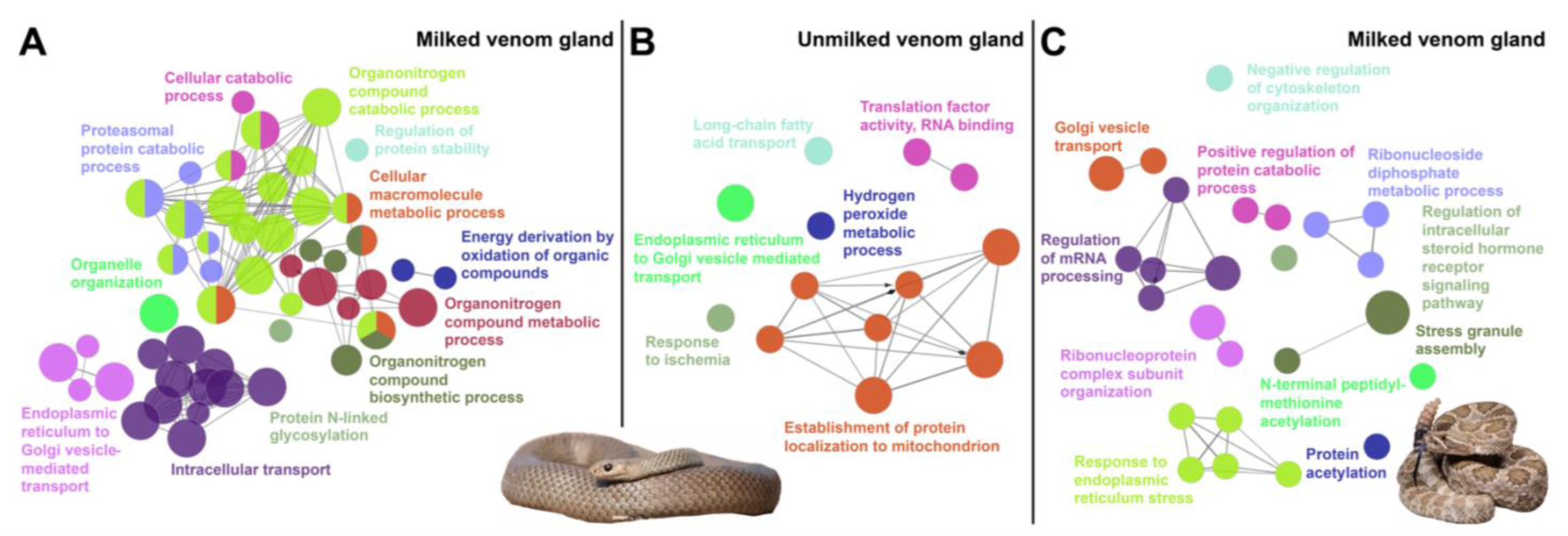
Gene networks and enriched biological processes for transcripts that are targeted by miRNAs expressed in snake venom glands. Transcripts from *Pseudonaja textilis* and *Crotalus viridis* genome annotations were used to predict mRNA targets for abundant miRNAs (over 100 Counts Per Million), in the (A) milked venom gland from *P. textilis,* (B) unmilked venom gland from *P. textilis,* and (C) milked venom gland from *C. viridis.* Gene networks were generated using *Homo sapiens* ortholog accessions of all targeted transcripts and the ClueGo app plug-in (Bindea et al. 2009) in Cytoscape (Shannon et al. 2003) to visual significant associated biological processes. Photo credit: *P. textilis,* Ákos Lumnitzer; *C. viridis,* Wolfgang Wüster.

We observed transcripts with multiple miRNA binding sites, as well as miRNAs that targeted multiple transcripts. One novel miRNA (3’-cgCCGCCGCCGTCGCCGCc-5’), uniquely expressed in the *P. textilis* MVG, had 510 targets. This miRNA did not share identity to any currently in the miRBase database (Griffiths-Jones et al. 2006), and due to the potential importance of this miRNA, we have named it *Pte-miR-1*. It is located on the *P. textilis* genome scaffold NW_020769308.1, 14,222,810 bp to 14,222,853 bp in the sense direction. Transcripts targeted by *Pte-miR-1* were for proteins involved in the following biological processes: ER to Golgi vesicle transport (GO:0006888), ubiquitin-dependent ERAD pathway (GO:0030433) and proteasome-mediated ubiquitin-dependent protein catabolic process (GO:0043161) (p-value=0.008) (Supplemental Figure 6).

## Discussion

Snakes of the families Elapidae and Viperidae are responsible for medically relevant snakebite envenoming worldwide, and venom variation between and within these families impacts the management of snakebite victims (Gutiérrez et al. 2017; Casewell et al. 2020). Investigations into how toxin genes, which most commonly arise from gene duplications, are variably expressed have been few due to the paucity of non-model genomes, including those for venomous snake species, and this question has been almost entirely unexplored for elapids. Using high throughput RNA-sequencing of MVGs and UVGs from the elapid *P. textilis* and viperids *C. viridis* and *C. tigris*, as well as comparative genomic approaches evaluating CREs in promoter regions of highly expressed toxin genes, we identified distinct toxin gene regulatory networks between these two venomous snake families.

3FTxs, group I PLA_2_s, KUNs, Snaclecs, and pseutarin C were the most abundant toxin families expressed in *P. textilis* venom glands (Figure 1A). Geographic variation in venom composition is found between South Australian (SA) and Queensland (QLD) *P. textilis* populations; SA snakes have an abundance of postsynaptic neurotoxins (3FTxs) and QLD snakes have greater amounts of textilotoxin and pseutarin C in their venoms (Skejić et al. 2015). We previously evaluated the venom proteomes of 12 *P. textilis* individuals from SA and found none contained all textilotoxin subunits, three individuals even entirely lacked textilotoxin subunits in their venoms (McCleary et al., 2016), but it was uncertain if this was due to method sensitivity limitations. Our high-throughput transcriptomic results from a *P. textilis* SA individual demonstrates a complete lack of gene expression of all four textilotoxin PLA_2_ subunits, and pseutarin C expression was only 2% of total toxin transcripts, whereas 3FTxs were highly expressed, exhibiting greater than 70% total toxin transcripts. These data demonstrate that *P. textilis* venom variation is partly due to differences in toxin gene transcription between populations. Evaluation of *P. textilis* genomes from different geographic regions would provide insight into resolving whether PLA_2_ genes for textilotoxin subunits are present but lacking transcription, or if these genes have been lost altogether in certain populations. This could contribute to the noted ‘brown snake paradox’, where although textilotoxin is a potent neurotoxin in *P. textilis* venom, *P. textilis* envenoming more frequency causes coagulopathy disturbances and rarely neurotoxicity (Barber et al. 2012).

Toxin expression in the viperid venom glands consisted primarily of myotoxins, SVSPs, group II PLA_2_s, and SVMPs for *C. viridis,* and group II PLA_2_s, SVSPs, BPPs, and VEGF for *C. tigris* (Figure 1B, C). From 0 to 96 hpvm, there was asynchrony in toxin synthesis, corroborating observations for the Palestine Viper (*Daboia palaestinae*) (Oron et al. 1978). However, different individual snakes were used at each time point for the viperid venom glands and intraspecific variation is likely. For this reason, we used a single *P. textilis* individual to investigate elapid toxin gene expression dynamics after venom milking.

Using MVGs and UVGs, we compared toxin gene expression between an elapid and viperids. Highly expressed *P. textilis* toxins exhibited fold-changes that were two-fold or less after venom milking. There were similar increases of 2.5 – 3.5-fold for 3FTxs and group I PLA_2_ genes 96 hpvm the venom gland of another elapid, *N. sputatrix* (Lachumanan et al. 1999). Increases over 40-fold for toxin gene expression were observed for the viperids *C. viridis* and *C. tigris*, with a SVSP gene peaking as high as 699-fold in *C. tigris* at 24 hpvm. Similar fold changes in SVSPs and SVMPs (over 50-fold and almost 20-fold, respectively) have been documented for the Puff Adder (*Bitis arietans*), although these levels of expression were determined from mRNA in venom, not venom glands (Currier et al. 2012). The lower fold-changes in toxin gene expression in the elapid MVG is due to the high levels of toxin expression in the UVG, suggesting elapid toxin genes might be more constitutively expressed. Greater fold changes in viperid toxin expression after venom milking is also likely because of the extensive physiological changes (cell elongation and expansion of ER) that take place in secretory cells of viperid MVGs (Oron and Bdolah 1973; Mackessy 1991), resulting in increased toxin synthesis capabilities. These physiological cell changes in viperid MVGs are largely absent in elapid MVGs (Kochva et al. 1982; Lachumanan et al. 1999). This also explains the absence of downregulated cellular component biogenesis and metabolic processes in the *P. textilis* MVG that was present in the viperid MVGs (Figure 2B and Figure 3B). Genes involved in striated muscle contraction were downregulated in the *P. textilis* MVG, aligning with the observation of the downregulation of troponin that was seen after venom milking of another elapid, the Many-banded Krait (*Bungarus multicinctus*) (Yin et al. 2020).

Genes related to the UPR, Notch signaling, and cholesterol homeostasis were enriched in the MVGs of both *P. textilis* and the two viperids. Increases in toxin synthesis, secretion, and posttranslational folding likely trigger upregulation of UPR and similar ER pathways to mediate ER stress and ensure protein quality control. Perry et al. (2020) identified the UPR pathway as a feedback regulatory mechanism increasing venom production in *C. viridis* (Perry et al. 2020), and UPR pathway components were present in the conserved metavenom network of venom glands from the viper *Protobothrops mucrosquamatus* (Barua and Mikheyev 2021), as well as conserved across venom glands in Metazoa (Zancolli et al. 2022). Cell-surface receptor Notch signaling is important in cell division and development (Bray 2006), and Notch signaling has been found to be critical for salivary gland cell growth and differentiation (Dang et al. 2009). In human salivary glands, β−adrenergic receptor activation upregulates Notch-mediated cell proliferation and differentiation of acinar cells (Wang et al. 2021), and this could be mediated by β-adrenergic receptor activation in the initiation of toxin synthesis in venom glands (Luna et al. 2009). This additionally supports conserved higher-level regulatory networks between venom glands and salivary glands (Barua and Mikheyev 2021). Cholesterol homeostasis enrichment is likely due to increases in vesicle-mediated transport and exocytosis in MVGs. Lipids that include cholesterol, phosphatidylinositol 4,5-bisphosphate and sphingolipids cluster as plasma membrane microdomains, concentrating and regulating SNARE proteins to create active exocytotic sites (Salaün et al. 2005). These data suggest that there is conservation in high-level cellular pathways regulating venom production for both elapids and viperids.

In contrast, we observed differences in chromosomal, TF, and CRE regulation between elapid-and viperid-specific venom production. Genes involved in chromatin organization/regulation, histone modification and transcription were upregulated in the *P. textilis* MVG (Figure 2A, C). This included chromatin-remodeler *SRCAP* and histone lysine methyltransferases *KMT2A, KMT2C* and *KMT2D*. KMT2C and KMT2D are known to function together as super-enhancers (Lai et al. 2017), and could be a potential mechanism to increase transcription related to venom production. Although *SRCAP, KMT2C* and *KMT2D* were also upregulated in *C. tigris* after venom milking, there were less genes overall associated with chromatin organization/regulation upregulated for viperid MVGs (Figure 3C). This could be due to the differences in chromosome locations of major toxins between elapids and viperids, macrochromosomes in elapids (Suryamohan et al. 2020; Zhang et al. 2022) and microchromosomes in viperid snakes (Shibata et al. 2018; Schield et al. 2019; Margres et al. 2021).

The transcription factor Sp1 was upregulated 79-fold in the *P. textilis* MVG, and Sp1 binding sites have been identified in promoter regions of *P. textilis* toxin genes for 3FTxs (Gong et al. 2001), group I PLA_2_s (Armugam et al. 2004) and the pseutarin C catalytic subunit (Reza et al. 2007), as well as toxin genes from other elapids (Supplemental Table 16). We also identified clusters of Sp1 CREs in toxin gene promoters belonging to different venom protein families and isoforms (Figure 4, 5), and although Sp1 binding sites were also present in viperid toxins, Sp1 was not upregulated to the same extent as observed for the elapid. Multiple Sp1 binding sites are known to enhance transcription (Pascal and Tjian 1991) and could be a regulatory mechanism to elevate the expression of toxins. Sp1 binding sites are highly enriched in promoter-proximal regions and instead of initiating the opening of chromatin structure, Sp1 activates transcription through interacting with general transcription machinery and p300, a core member of the enhancer complex (Ibañez-Tallon et al. 2002; Grossman et al. 2018). Thus, Sp1 has been associated with constitutive transcription (Ibañez-Tallon et al. 2002).

The shared Sp1 CREs of elapid 3FTx and group I PLA_2_ genes could allow for coordinated co-expression of these two venom protein families, which are both abundant toxins in the venom of many elapid snakes (Tasoulis and Isbister 2017), and for a potential pre-adaptive regulatory mechanism that could have contributed to the evolution of defensive venom in spitting cobras (Kazandjian et al. 2021). Notably, Sp1 sites have been identified in regulatory promoter regions for 3FTx and group I PLA_2_ genes in the spitting cobra *N. sputatrix* (Jeyaseelan et al. 2000; Ma et al. 2001). In *P. textilis,* shared Sp1 regulation of 3FTx, group I PLA_2_, and pseutarin C catalytic subunit genes could provide collective polygenic upregulation after venom milking. TATA boxes and associated factors were also identified shared across 3FTx, group I PLA_2_, and pseutarin C catalytic subunit genes. The TATA box regulates approximately 24% of genes in humans, and these genes are more tightly regulated by biotic or stress stimuli in comparison to TATA-less genes (Yang et al. 2007; Bae et al. 2015). However, given that the two group I PLA_2_ genes in *P. textilis* share identical 385 bp sequence upstream the TIS, but we found three group I PLA_2_ transcripts expressed at variable abundances (100,141 TPM to 1,846 TPM), and therefore additional regulatory mechanisms (e.g. chromatin structure, CREs farther upstream, number of CREs, enhancer sequences present at other sites, or RNA editing) likely contribute to the differential relative abundances of these toxins.

We identified potential *cis-*regulatory suppressors of *P. textilis* toxin genes: the non-conventional 3FTx found in the *P. textilis* genome had an IRF binding site not found to be present in expressed 3FTx homologs, and a c/EBP binding site was identified within the suppressor region of the pseutarin C catalytic subunit gene. Further, c/EBP was downregulated 0.82-fold in the *P. textilis* MVG. Gamma interferon response elements (γ−IRE) contributed to cell-specific silencing of group I PLA_2_ genes in *N. sputatrix* (Jeyaseelan et al. 2000), and c/EBP has been previously identified as a toxin suppressor CRE, as deletion of this site resulted in a 2-fold increase in promoter activity for a cytotoxic 3FTx in *N. sputatrix* (Ma et al. 2001).

Although NFI CREs are present in both elapid and viperid toxin gene promoters, NFI-family TFs appear to likely be of greater importance in the expression of viperid toxin genes in comparison to the elapid *P. textilis* as NFI genes were highly upregulated in both viperids. NFIA, NFIB, and NFIX, are RNA polymerase II core promoter binding TFs with binding sites present and accessible in promoter regions of multiple viperid toxin genes (SVMPs, SVSPs, and group II PLA_2_s) (Schield et al. 2019; Margres et al. 2021), potentially also coordinating the expression of these multiple gene families. NFI-family genes are ubiquitously expressed in different tissues but are known to regulate tissue-specific expression, including mammalian glands (Murtagh et al. 2003), through interactions with other TFs, members of the transcription initiation complex, and epigenetic regulators (Gronostajski 2000; O’Connor et al. 2016). NFI family binding sites are highly correlated with the center of nucleosome depletion regions, suggesting that their binding directly shapes local chromatin structures and can function as pioneer factors (Grossman et al. 2018; Adam et al. 2020). Pioneer factors are the first factors to engage target sites in chromatin and recruit histone modifying proteins, similar to many other identified viperid toxin gene TFs such as AP-1, CREB3, and FOX family TFs (Zaret and Carroll 2011; Perry et al. 2022). In addition, we found multiple CREs and associated *trans-*factors that were upregulated in viperids unique to group II PLA_2_s (RAR, USF1, and T3R), as well as varying upregulation of these factors between the two rattlesnake species, highlighting potential regulatory differences that could contribution to venom variation between and within the two snake families.

Post-transcriptional regulation of venom toxins by miRNAs has also been proposed (Durban et al. 2013). We observed that highly expressed miRNAs in the *P. textilis* MVG and UVG shared toxin transcript targets, with additional 3FTxs and pseutarin C non-catalytic subunit transcripts targeted in the UVG. Given that these targeted toxin transcripts are all major components of *P. textilis* venom, and we evaluated only top 10 most abundant miRNAs, there is a high likelihood that miRNAs could be regulating the translation of these toxin transcripts, including the downregulation of toxin translation in UVGs. Post-transcriptional miRNA regulation of venom toxins has been hypothesized to be responsible for ontogenetic venom variation in snakes (Durban et al. 2013; Durban et al. 2017; Durban et al. 2018), and our finding that there are miRNAs targeting both pseutarin C subunit transcripts could contribute to the ontogenetic shift in abundance of pseutarin C, as neonate *P. textilis* venoms lack this toxin complex and have venoms that fail to induce clot formation in plasma and whole blood (Jackson et al. 2016; Cipriani et al. 2017). Venom gland miRNAs from other *P. textilis* age classes will need to be evaluated to test this hypothesis. For *C. viridis,* we only identified two SVMP transcripts targeted, therefore less miRNA regulation of toxin transcripts was seen in the viperid venom gland. Viperid venom gland miRNAs that target SVMP transcripts have also been observed for the Mexican rattlesnakes *C. simus, C. tzabcan* and *C. culminatus* (Durban et al. 2013; Durban et al. 2017), suggesting this could be a common post-transcriptionally regulated toxin gene family in viperids.

Unique miRNA expression signatures were present in the *P. textilis* and *C. viridis* MVGs. This included the presence of *Pte-miR-1* in the *P. textilis* MVG with no known miRNA homology, the absence of *miR-375,* a highly abundant elapid miRNA, from the venom gland of *C. viridis,* and the unique presence of *miR-215-5p* in the *C. viridis* venom gland that was predicted to target SVMP toxin transcripts. Although there was overall a greater number of identified miRNAs in the viperid MVGs, miRNAs in the *P. textilis* MVG were more abundant and target predictions for miRNAs over 100 CPM demonstrated a greater extent of miRNA regulation in comparison to the viperid (Figure 6). For both elapid and viperid MVGs, miRNAs targeted transcripts for proteins involved in intracellular transport, but a greater extent of metabolic processes were targeted in the elapid MVG, and a greater number of processes related to mRNA processing and response to ER stress were targeted in the viperid MVGs.

As we can now produce snake venom gland organoids (Post et al. 2020; Puschhof et al. 2021), it is of even greater relevance to understand the epigenetic and genetic processes that regulate the expression of toxin genes. These insights would be useful to optimize *in vitro* toxin expression, with applications across fields in biotechnology and therapeutics (e.g., drug development from toxins and antivenom production), and would reduce the need for live venomous snakes to be used in research. The toxin gene regulatory mechanisms we have identified that potentially contribute to venom variation between elapid and viperid snakes will require additional evidence from larger snake venom gland and genome datasets. Our dataset is limited by only having one MVG and UVG for *P. textilis*, but this was done to avoid intraspecific variation in toxin expression, and the logistical and ethical considerations of sacrificing multiple animals. Additional sequencing approaches would be insightful, Assaying for Transposase-Accessible Chromatin followed by sequencing (ATAC-seq) would help to determine which toxin genes are in chromatin accessible regions, and Chromatin Immunoprecipitation followed by sequencing (ChIP-seq) for targeted histone markers (lysine 4 methylation of histone 3 associated with methyltransferases KMT2A, KMT2C, and KMT2D) and TFs (Sp1 and NFI-family TFs) would better determine their relationships to highly expressed toxin genes. Snake-specific TF antibodies are unfortunately not commercially available at this time, but venom gland organoids may offer an alternative approach where CRISPR/Cas9 technology could be used to Epitope Tag endogenous TFs for ChIP-seq (CETCh-seq) (Savic et al. 2015). Additionally, promoter activities and miRNA targets require experimental validation, and future venom gland organoid experiments could also facilitate such investigations by providing tissue-specific cell cultures for this work.

## Conclusions

Snake venom glands are tractable models to investigate gene regulation, as toxin gene expression and synthesis are upregulated in MVGs, and we can use manual venom milking to experimentally initiate these processes. Elapid and viperid venom glands are thought to share a common evolutionary origin, supported by anatomical and developmental evidence (Vonk et al. 2008), however, these two families have differing venom delivery systems, notably fang and venom gland morphology (Kerkkamp et al. 2015), and major differences in venom composition (Supplemental Table 17). Although there are shared toxin gene families and phylogenetic analysis of toxins suggests venom evolved once at the base of the advanced snake radiation (Fry and Wüster 2004), there are distinct toxin genes present in these two snake families and these toxin genes have differing chromosomal locations. Here, we used high-throughput RNA-seq to profile gene expression and miRNAs between MVGs and UVGs in these two snake families, in addition to performing comparative genomic analyses to identify *cis-* and *trans-*acting regulation of venom production in an elapid in comparison to previous viperid datasets (*C. viridis* and *C. tigris*). We identified CREs that are common across multiple toxin genes between these two snake families, but differences in potential key chromatin modifiers, TFs, and miRNAs regulating elapid and viperid toxin expression and synthesis (Supplemental Table 17). Therefore, elapid and viperid venom delivery systems, and their toxin genes and associated regulatory mechanisms, likely evolved independently.

## Materials and Methods

### Venom gland collection and RNA-sequencing

An adult *P. textilis* was collected from the Barossa Valley region, South Australia and maintained at Venom Supplies Pty Ltd. (Adelaide, SA, Australia). Venom was milked from the left gland, and the right gland was left untouched. Four days following venom milking, the snake was humanely euthanized, both glands dissected, placed in RNAlater (Thermo Fisher), and then shipped to the National University of Singapore. RNA was then extracted from the venom glands using Qiagen’s RNeasy Mini kit, following the manufacturer’s protocol. For mRNA libraries, 1 µg of total RNA was used as input into Illumina’s TruSeq RNA Sample Preparation v2 protocol. The small RNA library preps from the *P. textilis* MVG and UVG were completed using Illumina’s TruSeq Small RNA Sample Preparation protocol with 1 µg of total RNA as input. The finished small RNA library was loaded onto a 6% PAGE gel (Invitrogen) and a band of ∼170-320 bp was excised from the gel. The size-selected library was then extracted from the PAGE gel and recovered by ethanol precipitation. Quantitation of libraries was performed using Invitrogen’s Picogreen assay and the average library size determined by running the libraries on a Bioanalyzer DNA 1000 chip (Agilent). Library concentration was normalized to 2 nM and concentrations validated by qPCR on a ViiA-7 real-time thermocycler (Applied Biosystems), using qPCR primers recommended in Illumina’s qPCR protocol and Illumina’s PhiX control library used as a standard. Libraries were then pooled at equal volumes and the two library types sequenced on separate lanes of an Illumina HiSeq2500 rapid run at a final concentration of 11 pM, a read-length of 101 bp paired-end for the mRNA library and 51 bp single-end for the small RNA library.

An adult *C. viridis* rattlesnake of the same northeastern Colorado locality as used by Schield et al. (2019) for prior *C. viridis* genome sequencing was collected and venom milked. Four days following venom milking, the snake was humanely euthanized and both glands dissected. RNA isolation was completed for both glands combined, following the TRIzol reagent (Thermo Fisher Scientific) manufacturer’s protocol, with an additional overnight −20 °C incubation in 300 μL 100% ethanol with 40 μL 3 M sodium acetate, and resuspension in nuclease-free H2O following incubation. Two different Illumina sequencing libraries were prepared, one selecting mRNA and a second selecting small RNA. The same mRNA library preparation protocol as described above for *P. textilis* was also followed for *C. viridis and* paired-end sequencing completed to a read length of 150 bp. The small RNA library preparation was done with 1 µg of total RNA used as input into the NEBNext Small RNA Library Prep Set for Illumina (New England BioLabs) and an AMPure XP Bead (Beckman Coulter) selection step for a 110-160 bp size range. These small RNA libraries were sequenced to 75 bp, single-end.

### Venom gland *de novo* transcriptome assembly, toxin annotation and expression

Sequenced reads were assessed with the Java program FastQC (Babraham Institute Bioinformatics, UK) to confirm that all adapters and low quality reads (<Q20) were removed before assembly. To obtain a comprehensive *de novo* venom gland transcriptome assembly for *P. textilis*, separate assemblies for each gland were completed, and three assemblers used: 1) ABySS (release v1.5.0) (Birol et al. 2009; Simpson et al. 2009) with paired-end default parameters and k-mer sizes 30 to 66, increased in increments of 4, and merged with TransABySS (v1.5.1) (Robertson et al. 2010), 2) Trinity (release v2014-07-17) (Grabherr et al. 2011) with genome-guided assembly default parameters using Bowtie2 (v2.2.6) (Langmead and Salzberg 2012) aligned reads to the *P. textilis* genome (assembly EBS10Xv2-PRI), and 3) Extender (Rokyta et al. 2012) with 10,000 starting seeds, where seeds were reads first merged with PEAR (Paired-End read mergeR; v0.9.6 using default parameters) (Zhang et al. 2014) and seed extensions required 100 nucleotide overlaps and quality scores of at least 30. For *C. viridis,* a *de novo* venom gland transcriptome assembly was completed using the same approached detailed above for *P. textilis,* except excluding ABySS and with an additional Trinity assembly using *de novo* parameters. For the *de novo* assembled transcriptomes for both species, contigs less than 150 nucleotides and redundancies between assemblies were removed with CD-HIT (v4.6.6) (Li and Godzik 2006; Fu et al. 2012). Coding contigs were then identified with EvidentialGene (downloaded May 2018) (Gilbert 2013). Abundances of the coding contig set were determined with RSEM (RNA-seq by Expectation Maximization, v1.3.0) (Li and Dewey 2011), using the aligner Bowtie2 (v2.2.6) (Langmead and Salzberg 2012). Contigs less than 1 TPM (Transcript Per Million) were filtered out, and the remaining contigs annotated with Diamond (Buchfink et al. 2014) and BLASTx (E-value 10^-05^ cut-off) searches against the *P. textilis* genome-predicted protein set and the National Center for Biotechnology Information (NCBI) non-redundant protein database. OrfPredictor (v3) (Min et al. 2005) was used for final coding and protein sequence prediction with BLASTx input to aid in proper transcript translation. Venomix (Macrander et al. 2018) was used to help to identify all toxin transcripts in addition to toxins being manually evaluated to determine if venom proteins were full-length, shared sequence identity to currently known toxins, and contained a conserved signal peptide sequence within each venom protein family.

### Venom gland gene expression and *cis-*regulatory element predictions

From *P. textilis* genome annotations (assembly EBS10Xv2-PRI), the global transcriptome (34,614 predicted transcripts) was used as a reference for aligning reads originating from the *P. textilis* MVG and UVG. For rattlesnake MVGs and UVGs, the following NCBI data sets were used: SRR11524062 (*C. tigris* UVG), SRR11524063 (*C. tigris* UVG), SRR11524059 (*C. tigris* 24 hpvm), SRR11524060 (*C. tigris* 24 hpvm), SRR11524050 (*C. tigris* 96 hpvm), SRR11524051 (*C. tigris* 96 hpvm), SRR7401989 (*C. viridis* UVG), SRR7402004 (*C. viridis* 24 hpvm), and SRR7402005 (*C. viridis* 72 hpvm). Reads from these data sets were aligned to annotated transcriptomes from genome assemblies (UTA_CroVir_3.0 and ASM1654583v1 for *C. viridis* and *C. tigris*, respectively), in addition to myotoxin transcripts from the *de novo* assembled venom gland transcriptome for *C. viridis*. Transcript abundances were determined with Bowtie2 (v2.2.6) (Langmead and Salzberg 2012) read alignments and RSEM (Li and Dewey 2011). RSEM output of expected counts, transcript length and TPM were used as input into GFOLD (Feng et al. 2012) to identify transcript fold-changes between the conditions. A Gene Set Enrichment Analysis (GSEA) (Mootha et al. 2003; Subramanian et al. 2005) was performed using RSEM estimated transcript abundances as input. Transcripts were searched against the UniProt *Homo sapiens* protein database (The UniProt 2021) with Diamond BLASTx (v0.8.34) (Buchfink et al. 2014) to identify orthologs. UniProt accessions were then entered into DAVID Bioinformatics Resources 6.8 (Huang da et al. 2009a) to identify functional annotations and pathways. In addition, gene networks were constructed using the ClueGo app plug-in (Bindea et al. 2009) in Cytoscape (Shannon et al. 2003). *Cis*-regulatory element predictions were completed upstream toxin gene transcription start sites, up to 700 bp if available, using the online server AliBaba2.1 (http://gene-regulation.com/pub/programs/alibaba2/) with the TRANSFAC 4.0 database embedded within the webserver. GFOLD determined fold-changes in MVGs for *trans*-factors associated with CREs were then evaluated.

### Venom gland microRNA expression and target prediction

TrueSeq small RNA library adapters were trimmed with the fastx_clipper tool, provided in the FASTX-Toolkit (Hannon lab Cold Spring Harbor Laboratory). Trimmomatic (Bolger et al. 2014) was used to filter out low quality (< Q20) reads, evaluated with a sliding window of 4 nucleotides. To filter out rRNA, reads were then aligned with Bowtie2 (v2.2.6) to known snake rRNA sequences. Non-rRNA reads were used as input into miRDeep2 (v2.0.1.2) (Friedländer et al. 2012) to identify species-specific miRNAs from the *P. textilis* genome (GCF900518735.1, assembly EBS10Xv2-PRI) and *C. viridis* (UTA_CroVir_3.0). Expression levels of miRNAs in venom glands were estimated by normalization to Counts Per Million (CPMs; CPM = mature miRNA reads / total mapped miRNA reads * 10^6^). Target prediction was performed with the position-weighted local alignment miRanda (v3.3) algorithm (Enright et al. 2003; John et al. 2004). A free energy value of at least -19 kcal/mol was used as thresholds for toxin transcript target identification, using only contigs coding for full length venom proteins from the *de novo* assembled transcriptome from *P. textilis* and toxin transcripts from the annotated in *C. viridis* genome, in addition to the *de novo* assembled myotoxin transcript. A stricter free energy value of at least -30 kcal/mol was used for target identification from genome annotated transcriptome datasets. Non-toxin transcripts targeted were searched against the UniProt *Homo sapiens* protein database (The UniProt 2021) with Diamond BLASTx (v0.8.34) (Buchfink et al. 2014) to identify orthologs, and accessions were then entered into DAVID Bioinformatics Resources 6.8 (Huang da et al. 2009a) to identify functional annotations and pathways. In addition, gene networks were constructed using the ClueGo app plug-in (Bindea et al. 2009) in Cytoscape (Shannon et al. 2003).

## Data access

All next-generation sequencing data was submitted to NCBI under BioProject ID PRJNA931953 (BioSample SAMN33139130 for *P. textilis* and SAMN33139131 for *C. viridis*).

## Acknowledgements

This study was supported by an academic research grant from the National University of Singapore to RMK.

## References

Adam RC, Yang H, Ge Y, Infarinato NR, Gur-Cohen S, Miao Y, Wang P, Zhao Y, Lu CP, Kim JE et al. 2020. NFI transcription factors provide chromatin access to maintain stem cell identity while preventing unintended lineage fate choices. Nat Cell Biol 22: 640–650.

Armugam A, Gong N, Li X, Siew PY, Chai SC, Nair R, Jeyaseelan K. 2004. Group IB phospholipase A2 from *Pseudonaja textilis*. Arch Biochem Biophys 421: 10–20.

Bae S-H, Han HW, Moon J. 2015. Functional analysis of the molecular interactions of TATA box-containing genes and essential genes. PLOS One 10: e0120848.

Barber CM, Isbister GK, Hodgson WC. 2012. Solving the ‘Brown snake paradox’: *in vitro* characterisation of Australasian snake presynaptic neurotoxin activity. Toxicol lett 210: 318–323.

Bartel DP. 2004. MicroRNAs: Genomics, biogenesis, mechanism, and function. Cell 116: 281–297.

Barua A, Mikheyev AS. 2021. An ancient, conserved gene regulatory network led to the rise of oral venom systems. Proc Natl Acad Sci USA 118: e2021311118.

Bindea G, Mlecnik B, Hackl H, Charoentong P, Tosolini M, Kirilovsky A, Fridman WH, Pagès F, Trajanoski Z, Galon J. 2009. ClueGO: a Cytoscape plug-in to decipher functionally grouped gene ontology and pathway annotation networks. Bioinformatics 25: 1091–1093.

Birol I, Jackman SD, Nielsen CB, Qian JQ, Varhol R, Stazyk G, Morin RD, Zhao Y, Hirst M, Schein JE et al. 2009. De novo transcriptome assembly with ABySS. Bioinformatics 25: 2872–2877.

Birrell GW, Earl S, Masci PP, de Jersey J, Wallis TP, Gorman JJ, Lavin MF. 2006. Molecular diversity in venom from the Australian Brown snake, *Pseudonaja textilis*. Mol Cell Proteomics 5: 379–389.

Bolger AM, Lohse M, Usadel B. 2014. Trimmomatic: a flexible trimmer for Illumina sequence data. Bioinformatics 30: 2114–2120.

Bray SJ. 2006. Notch signalling: a simple pathway becomes complex. Nat Rev Mol Cell Biol 7: 678–689.

Broad AJ, Sutherland SK, Coulter AR. 1979. The lethality in mice of dangerous Australian and other snake venom. Toxicon 17: 661–664.

Buchfink B, Xie C, Huson DH. 2014. Fast and sensitive protein alignment using DIAMOND. Nat methods 12: 59.

Calvete JJ, Pérez A, Lomonte B, Sánchez EE, Sanz L. 2012. Snake venomics of *Crotalus tigris*: the minimalist toxin arsenal of the deadliest Nearctic rattlesnake venom. Evolutionary Clues for generating a pan-specific antivenom against crotalid type II venoms. J Proteome Res 11: 1382–1390.

Cameron DL, Tu AT. 1978. Chemical and functional homology of myotoxin a from prairie rattlesnake venom and crotamine from South American rattlesnake venom. Biochim et Biophys Acta 532: 147–154.

Casewell NR, Jackson TNW, Laustsen AH, Sunagar K. 2020. Causes and consequences of snake venom variation. Trends Pharmacol Sci 41: 570–581.

Chang LS, Chou YC, Lin SR, Wu BN, Lin J, Hong E, Sun YJ, Hsiao CD. 1997. A novel neurotoxin, cobrotoxin b, from *Naja naja atra* (Taiwan cobra) venom: purification, characterization, and gene organization. J Biochem 122: 1252–1259.

Cho YW, Hong T, Hong S, Guo H, Yu H, Kim D, Guszczynski T, Dressler GR, Copeland TD, Kalkum M et al. 2007. PTIP associates with MLL3-and MLL4-containing histone H3 lysine 4 methyltransferase complex. J Biol Chem 282: 20395–20406.

Cipriani V, Debono J, Goldenberg J, Jackson TNW, Arbuckle K, Dobson J, Koludarov I, Li B, Hay C, Dunstan N et al. 2017. Correlation between ontogenetic dietary shifts and venom variation in Australian brown snakes (*Pseudonaja*). Comp Biochem Physiol C Toxicol Pharmacol 197: 53–60.

Conway E, Jerman E, Healy E, Ito S, Holoch D, Oliviero G, Deevy O, Glancy E, Fitzpatrick DJ, Mucha M et al. 2018. A Family of vertebrate-specific polycombs encoded by the LCOR/LCORL genes balance PRC2 subtype activities. Mol Cell 70: 408–421.e408.

Currier R, Juan C, Libia S, Robert H, Paul R, Simon W. 2012. Unusual stability of messenger RNA in snake venom reveals gene expression dynamics of venom replenishment. PLOS One 7: 1–10.

Dang H, Lin AL, Zhang B, Zhang H-M, Katz MS, Yeh C-K. 2009. Role for Notch signaling in salivary acinar cell growth and differentiation. Dev Dyn 238: 724–731.

de Vries E, van Driel W, van den Heuvel SJ, van der Vliet PC. 1987. Contactpoint analysis of the HeLa nuclear factor I recognition site reveals symmetrical binding at one side of the DNA helix. EMBO J 6: 161–168.

Deshimaru M, Ogawa T, Nakashima K, Nobuhisa I, Chijiwa T, Shimohigashi Y, Fukumaki Y, Niwa M, Yamashina I, Hattori S et al. 1996. Accelerated evolution of crotalinae snake venom gland serine proteases. FEBS Lett 397: 83–88.

Doench JG, Sharp PA. 2004. Specificity of microRNA target selection in translational repression. Genes Dev 18: 504–511.

Durban J, Pérez A, Sanz L, Gómez A, Bonilla F, Rodríguez S, Chacón D, Sasa M, Angulo Y, Gutiérrez JM et al. 2013. Integrated “omics” profiling indicates that miRNAs are modulators of the ontogenetic venom composition shift in the Central American rattlesnake, Crotalus simus simus. BMC Genom 14: 234.

Durban J, Sanz L, Trevisan-Silva D, Neri-Castro E, Alagón A, Calvete JJ. 2017. Integrated venomics and venom gland transcriptome analysis of juvenile and adult Mexican rattlesnakes *Crotalus simus*, C. tzabcan, and C. culminatus revealed miRNA-modulated ontogenetic shifts. J Proteome Res 16: 3370–3390.

Durban J, Sasa M, Calvete JJ. 2018. Venom gland transcriptomics and microRNA profiling of juvenile and adult yellow-bellied sea snake, *Hydrophis platurus*, from Playa del Coco (Guanacaste, Costa Rica). Toxicon 153: 96–105.

Enright AJ, John B, Gaul U, Tuschl T, Sander C, Marks DS. 2003. MicroRNA targets in Drosophila. Genome Biol 5: R1.

Feng J, Meyer CA, Wang Q, Liu JS, Shirley Liu X, Zhang Y. 2012. GFOLD: a generalized fold change for ranking differentially expressed genes from RNA-seq data. Bioinformatics 28: 2782–2788.

Fernandes I, Bastien Y, Wai T, Nygard K, Lin R, Cormier O, Lee HS, Eng F, Bertos NR, Pelletier N et al. 2003. Ligand-dependent nuclear receptor corepressor LCoR functions by histone deacetylase-dependent and -independent mechanisms. Mol Cell 11: 139–150.

Flight S, Mirtschin P, Masci PP. 2006. Comparison of active venom components between Eastern brown snakes collected from South Australia and Queensland. Ecotoxicology 15: 133–141.

Friedländer MR, Mackowiak SD, Li N, Chen W, Rajewsky N. 2012. miRDeep2 accurately identifies known and hundreds of novel microRNA genes in seven animal clades. Nucleic Acids Res 40: 37–52.

Friedman RC, Farh KK, Burge CB, Bartel DP. 2009. Most mammalian mRNAs are conserved targets of microRNAs. Genome Res 19: 92–105.

Fry BG, Wüster W. 2004. Assembling an arsenal: origin and evolution of the snake venom proteome inferred from phylogenetic analysis of toxin sequences. Mol Biol Evol 21: 870–883.

Fu L, Niu B, Zhu Z, Wu S, Li W. 2012. CD-HIT: accelerated for clustering the next-generation sequencing data. Bioinformatics 28: 3150–3152.

Fujimi TJ, Yasuoka S, Ogura E, Tsuchiya T, Tamiya T. 2004. Comparative analysis of gene expression mechanisms between group IA and IB phospholipase A2 genes from sea snake Laticauda semifasciata. Gene 332: 179–190.

Gilbert D. 2013. Gene-omes built from mRNA seq not genome DNA. doi:10.7490/f1000research.1112594.1.

Gong N, Armugam A, Jeyaseelan K. 2000. Molecular cloning, characterization and evolution of the gene encoding a new group of short-chain α-neurotoxins in an Australian elapid, *Pseudonaja textilis*. FEBS Lett 473: 303–310.

Gong N, Armugam A, Mirtschin P, Jeyaseelan K. 2001. Cloning and characterization of the pseudonajatoxin b precursor. Biochemical J 358: 647–656.

Grabherr MG, Haas BJ, Yassour M, Levin JZ, Thompson DA, Amit I, Adiconis X, Fan L, Raychowdhury R, Zeng Q et al. 2011. Full-length transcriptome assembly from RNA-Seq data without a reference genome. Nat Biotech 29: 644–652.

Griffiths-Jones S, Grocock RJ, van Dongen S, Bateman A, Enright AJ. 2006. miRBase: microRNA sequences, targets and gene nomenclature. Nucleic Acids Res 34: D140–D144.

Gronostajski RM. 2000. Roles of the NFI/CTF gene family in transcription and development. Gene 249: 31–45.

Grossman SR, Engreitz J, Ray JP, Nguyen TH, Hacohen N, Lander ES. 2018. Positional specificity of different transcription factor classes within enhancers. Proc Natl Acad Sci USA 115: E7222–E7230.

Gutiérrez JM, Calvete JJ, Habib AG, Harrison RA, Williams DJ, Warrell DA. 2017. Snakebite envenoming. Nat Rev Dis Primers 3: 17063.

Han SX, Kwong S, Ge R, Kolatkar PR, Woods AE, Blanchet G, Kini RM. 2016. Regulation of expression of venom toxins: silencing of prothrombin activator trocarin D by AG-rich motifs. FASEB J 30: 2411–2425.

Heinrikson RL, Krueger ET, Keim PS. 1977. Amino acid sequence of phospholipase A2-alpha from the venom of Crotalus adamanteus. A new classification of phospholipases A2 based upon structural determinants. J Biol Chem 252: 4913–4921.

Hite LA, Jia LG, Bjarnason JB, Fox JW. 1994. cDNA sequences for four snake venom metalloproteinases: structure, classification, and their relationship to mammalian reproductive proteins. Arch Biochem Biophys 308: 182–191.

Ho CL, Lee CY. 1981. Presynaptic actions of Mojave toxin isolated from Mojave rattlesnake (*Crotalus scutulatus*) venom. Toxicon 19: 889–892.

Huang da W, Sherman BT, Lempicki RA. 2009a. Bioinformatics enrichment tools: paths toward the comprehensive functional analysis of large gene lists. Nucleic Acids Res 37: 1–13.

Huang da W, Sherman BT, Lempicki RA. 2009b. Systematic and integrative analysis of large gene lists using DAVID bioinformatics resources. Nat Protoc 4: 44–57.

Ibañez-Tallon I, Ferrai C, Longobardi E, Facetti I, Blasi F, Crippa MP. 2002. Binding of Sp1 to the proximal promoter links constitutive expression of the human uPA gene and invasive potential of PC3 cells. Blood 100: 3325–3332.

Jackson K. 2003. The evolution of venom-delivery systems in snakes. Zool J Linn Soc 137: 337–354.

Jackson TNW, Koludarov I, Ali SA, Dobson J, Zdenek CN, Dashevsky D, Op den Brouw B, Masci PP, Nouwens A, Josh P, et al. 2016. Rapid radiations and the race to redundancy: An investigation of the evolution of Australian elapid snake venoms. Toxins 8: 309.

Jeyaseelan K, Armugam A, Donghui M, Tan N-H. 2000. Structure and phylogeny of the venom group I phospholipase A2 gene. Mol Bio Evol 17: 1010–1021.

John B, Enright AJ, Aravin A, Tuschl T, Sander C, Marks DS. 2004. Human microRNA targets. PLoS Biol 2: e363.

Joseph JS, Chung MC, Jeyaseelan K, Kini RM. 1999. Amino acid sequence of trocarin, a prothrombin activator from *Tropidechis carinatus* venom: its structural similarity to coagulation factor Xa. Blood 94: 621–631.

Kardong KV. 1982. The evolution of the venom apparatus in snakes from colubrids to viperids and elapids. Mem Inst Butantan 46: 105–118.

Kazandjian TD, Petras D, Robinson SD, van Thiel J, Greene HW, Arbuckle K, Barlow A, Carter DA, Wouters RM, Whiteley G et al. 2021. Convergent evolution of pain-inducing defensive venom components in spitting cobras. Science 371: 386–390.

Kerkkamp HMI, Casewell NR, Vonk FJ. 2015. Evolution of the Snake Venom Delivery System. In Evolution of Venomous Animals and Their Toxins, doi:10.1007/978-94-007-6727-0_11-1 (ed. P Gopalakrishnakone, A Malhotra), pp. 1–11. Springer Netherlands, Dordrecht.

Kochva E. 1987. The origin of snakes and evolution of the venom apparatus. Toxicon 25: 65–106.

Kochva E, Bdolah A, Wollberg Z. 1993. Sarafotoxins and endothelins: evolution, structure and function. Toxicon 31: 541–568.

Kochva E, Tönsing L, Louw AI, Liebenberg NvdW, Visser L. 1982. Biosynthesis, secretion and *in vivo* isotopic labelling of venom of the Egyptian cobra, *Naja haje annulifera*. Toxicon 20: 615–635.

Kwong S, Woods AE, Mirtschin PJ, Ge R, Kini RM. 2009. The recruitment of blood coagulation factor X into snake venom gland as a toxin: The role of promoter Cis-elements in its expression. Thromb Haemost 103: 469–78.

Lachumanan R, Armugam A, Durairaj P, Gopalakrishnakone P, Tan CH, Jeyaseelan K. 1999. In situ hybridization and immunohistochemical analysis of the expression of cardiotoxin and neurotoxin genes in *Naja naja sputatrix*. J Histochem Cytochem 47: 551–560.

Lai B, Lee J-E, Jang Y, Wang L, Peng W, Ge K. 2017. MLL3/MLL4 are required for CBP/p300 binding on enhancers and super-enhancer formation in brown adipogenesis. Nucleic Acids Res 45: 6388–6403.

Langmead B, Salzberg SL. 2012. Fast gapped-read alignment with Bowtie 2. Nat Methods 9: 357–359.

Li B, Dewey CN. 2011. RSEM: accurate transcript quantification from RNA-Seq data with or without a reference genome. BMC Bioinform 12: 323.

Li W, Godzik A. 2006. Cd-hit: a fast program for clustering and comparing large sets of protein or nucleotide sequences. Bioinformatics 22: 1658–1659.

Luna MS, Hortencio TM, Ferreira ZS, Yamanouye N. 2009. Sympathetic outflow activates the venom gland of the snake *Bothrops jararaca* by regulating the activation of transcription factors and the synthesis of venom gland proteins. J Exp Biol 212: 1535–1543.

Ma D, Armugam A, Jeyaseelan K. 2001. Expression of cardiotoxin-2 gene. Cloning, characterization and deletion analysis of the promoter. Eur J Biochem 268: 1844–1850.

Ma D, Armugam A, Jeyaseelan K. 2002. Alpha-neurotoxin gene expression in *Naja sputatrix*: identification of a silencer element in the promoter region. Arch Biochem Biophys 404: 98–105.

Mackessy SP. 1991. Morphology and ultrastructure of the venom glands of the northern pacific rattlesnake *Crotalus viridis oreganus*. J Morphol 208: 109–128.

Mackessy SP. 2010. The field of reptile toxinology: snakes, lizards and their venoms. In Handbook of Venoms and Toxins of Reptiles, (ed. M SP), pp. 2–23. CRC Press/Taylor & Francis Group, Boca Raton, FL.

Macrander J, Panda J, Janies D, Daly M, Reitzel AM. 2018. Venomix: a simple bioinformatic pipeline for identifying and characterizing toxin gene candidates from transcriptomic data. PeerJ 6: e5361.

Margres MJ, Rautsaw RM, Strickland JL, Mason AJ, Schramer TD, Hofmann EP, Stiers E, Ellsworth SA, Nystrom GS, Hogan MP et al. 2021. The Tiger Rattlesnake genome reveals a complex genotype underlying a simple venom phenotype. Proc Natl Acad Sci USA 118: e2014634118.

McCleary RJ, Sridharan S, Dunstan NL, Mirtschin PJ, Kini RM. 2016. Proteomic comparisons of venoms of long-term captive and recently wild-caught Eastern brown snakes (*Pseudonaja textilis*) indicate venom does not change due to captivity. J Proteomics 144: 51–62.

Min XJ, Butler G, Storms R, Tsang A. 2005. OrfPredictor: predicting protein-coding regions in EST-derived sequences. Nucleic Acids Res 33: W677–W680.

Miwa JM, Ibaňez-Tallon I, Crabtree GW, Sánchez R, Šali A, Role LW, Heintz N. 1999. lynx1, an endogenous toxin-like modulator of nicotinic acetylcholine receptors in the mammalian CNS. Neuron 23: 105–114.

Monroy MA, Schott NM, Cox L, Chen JD, Ruh M, Chrivia JC. 2003. SNF2-related CBP activator protein (SRCAP) functions as a coactivator of steroid receptor-mediated transcription through synergistic interactions with CARM-1 and GRIP-1. Mol Endocrinol 17: 2519–2528.

Mootha VK, Lindgren CM, Eriksson K-F, Subramanian A, Sihag S, Lehar J, Puigserver P, Carlsson E, Ridderstråle M, Laurila E et al. 2003. PGC-1α-responsive genes involved in oxidative phosphorylation are coordinately downregulated in human diabetes. Nat Genet 34: 267–273.

Murtagh J, Martin F, Gronostajski RM. 2003. The nuclear factor I (NFI) gene family in mammary gland development and function. J Mammary Gland Biol and Neoplasia 8: 241–254.

Nakamura H, Murakami T, Hattori S, Sakaki Y, Ohkuri T, Chijiwa T, Ohno M, Oda-Ueda N. 2014. Epithelium specific ETS transcription factor, ESE-3, of Protobothrops flavoviridis snake venom gland transactivates the promoters of venom phospholipase A2 isozyme genes. Toxicon 92: 133–139.

Nakamura T, Mori T, Tada S, Krajewski W, Rozovskaia T, Wassell R, Dubois G, Mazo A, Croce CM, Canaani E. 2002. ALL-1 is a histone methyltransferase that assembles a supercomplex of proteins involved in transcriptional regulation. Mol Cell 10: 1119–1128.

Nirthanan S, Gopalakrishnakone P, Gwee MCE, Khoo HE, Kini RM. 2003. Non-conventional toxins from elapid venoms. Toxicon 41: 397–407.

Nishiyama M, Skoultchi AI, Nakayama KI. 2012. Histone H1 recruitment by CHD8 is essential for suppression of the Wnt-β-catenin signaling pathway. Mol Cell Biol 32: 501–512.

Nobuhisa I, Nakashima K, Deshimaru M, Ogawa T, Shimohigashi Y, Fukumaki Y, Sakaki Y, Hattori S, Kihara H, Ohno M. 1996. Accelerated evolution of *Trimeresurus okinavensis* venom gland phospholipase A2 isozyme-encoding genes. Gene 172: 267–272.

O’Brien J, Hayder H, Zayed Y, Peng C. 2018. Overview of microRNA biogenesis, mechanisms of actions, and circulation. Front Endocrinol 9: 402–402.

O’Connor L, Gilmour J, Bonifer C. 2016. The role of the ubiquitously expressed transcription factor Sp1 in tissue-specific transcriptional regulation and in disease. Yale J Biol Med 89: 513–525.

Ohno M, Menez R, Ogawa T, Danse JM, Shimohigashi Y, Fromen C, Ducancel F, Zinn-Justin S, Le Du MH, Boulain JC et al. 1998. Molecular evolution of snake toxins: is the functional diversity of snake toxins associated with a mechanism of accelerated evolution? Prog Nucleic Acid Res Mol Biol 59: 307–364.

Oron U, Bdolah A. 1973. Regulation of protein synthesis in the venom gland of viperid snakes. J Cell Biol 56: 177–190.

Oron U, Kinamon S, Bdolah A. 1978. Asynchrony in the synthesis of secretory proteins in the venom gland of the snake *Vipera palaestinae*. Biochem J 174: 733–739.

Paine MJ, Desmond HP, Theakston RD, Crampton JM. 1992. Gene expression in *Echis carinatus* (carpet viper) venom glands following milking. Toxicon 30: 379–386.

Pascal E, Tjian R. 1991. Different activation domains of Sp1 govern formation of multimers and mediate transcriptional synergism. Genes Dev 5: 1646–1656.

Perry BW, Gopalan SS, Pasquesi GIM, Schield DR, Westfall AK, Smith CF, Koludarov I, Chippindale PT, Pellegrino MW, Chuong EB et al. 2022. Snake venom gene expression is coordinated by novel regulatory architecture and the integration of multiple co-opted vertebrate pathways. Genome Res doi:10.1101/gr.276251.121.

Perry BW, Schield DR, Westfall AK, Mackessy SP, Castoe TA. 2020. Physiological demands and signaling associated with snake venom production and storage illustrated by transcriptional analyses of venom glands. Sci Rep 10: 18083.

Philipsen S, Suske G. 1999. A tale of three fingers: the family of mammalian Sp/XKLF transcription factors. Nucleic Acids Res 27: 2991–3000.

Post Y, Puschhof J, Beumer J, Kerkkamp HM, de Bakker MAG, Slagboom J, de Barbanson B, Wevers NR, Spijkers XM, Olivier T et al. 2020. Snake venom gland organoids. Cell 180: 233–247.e221.

Puschhof J, Post Y, Beumer J, Kerkkamp HM, Bittenbinder M, Vonk FJ, Casewell NR, Richardson MK, Clevers H. 2021. Derivation of snake venom gland organoids for *in vitro* venom production. Nat Protocols 16: 1494–1510.

Rao VS, Kini RM. 2002. Pseutarin C, a prothrombin activator from *Pseudonaja textilis* venom: its structural and functional similarity to mammalian coagulation factor Xa-Va complex. Thromb Haemost 88: 611–619.

Rao VS, Swarup S, Kini RM. 2003. The nonenzymatic subunit of pseutarin C, a prothrombin activator from eastern brown snake (*Pseudonaja textilis*) venom, shows structural similarity to mammalian coagulation factor V. Blood 102: 1347–1354.

Rao VS, Swarup S, Manjunatha Kini R. 2004. The catalytic subunit of pseutarin C, a group C prothrombin activator from the venom of *Pseudonaja textilis*, is structurally similar to mammalian blood coagulation factor Xa. Thromb Haemost 92: 509–521.

Reeks T, Lavergne V, Sunagar K, Jones A, Undheim E, Dunstan N, Fry B, Alewood PF. 2016. Deep venomics of the *Pseudonaja* genus reveals inter-and intraspecific variation. J Proteomics 133: 20–32.

Reza MA, Swarup S, Kini RM. 2007. Structure of two genes encoding parallel prothrombin activators in *Tropidechis carinatus* snake: gene duplication and recruitment of factor X gene to the venom gland. J Thromb Haemost 5: 117–126.

Robertson G, Schein J, Chiu R, Corbett R, Field M, Jackman SD, Mungall K, Lee S, Okada HM, Qian JQ et al. 2010. De novo assembly and analysis of RNA-seq data. Nat Methods 7: 909.

Rokyta DR, Lemmon AR, Margres MJ, Aronow K. 2012. The venom-gland transcriptome of the eastern diamondback rattlesnake (*Crotalus adamanteus*). BMC Genom 13: 312.

Rotenberg D, Bamberger ES, Kochva E. 1971. Studies on ribonucleic acid synthesis in the venom glands of *Vipera palaestinae* (Ophidia, Reptilia). J Biochem 121: 609–612.

Salaün C, Gould GW, Chamberlain LH. 2005. Lipid raft association of SNARE proteins regulates exocytosis in PC12 cells. J Biol Chem 280: 19449–19453.

Savic D, Partridge EC, Newberry KM, Smith SB, Meadows SK, Roberts BS, Mackiewicz M, Mendenhall EM, Myers RM. 2015. CETCh-seq: CRISPR epitope tagging ChIP-seq of DNA-binding proteins. Genome Res 25: 1581–1589.

Saviola AJ, Pla D, Sanz L, Castoe TA, Calvete JJ, Mackessy SP. 2015. Comparative venomics of the Prairie Rattlesnake (*Crotalus viridis viridis*) from Colorado: Identification of a novel pattern of ontogenetic changes in venom composition and assessment of the immunoreactivity of the commercial antivenom CroFab(R). J Proteomics 121: 28–43.

Schield DR, Card DC, Hales NR, Perry BW, Pasquesi GM, Blackmon H, Adams RH, Corbin AB, Smith CF, Ramesh B et al. 2019. The origins and evolution of chromosomes, dosage compensation, and mechanisms underlying venom regulation in snakes. Genome Res 29: 590–601.

Seilhamer JJ, Pruzanski W, Vadas P, Plant S, Miller JA, Kloss J, Johnson LK. 1989. Cloning and recombinant expression of phospholipase A2 present in rheumatoid arthritic synovial fluid. J Biol Chem 264: 5335–5338.

Shannon P, Markiel A, Ozier O, Baliga NS, Wang JT, Ramage D, Amin N, Schwikowski B, Ideker T. 2003. Cytoscape: a software environment for integrated models of biomolecular interaction networks. Genome Res 13: 2498–2504.

Sherman BT, Hao M, Qiu J, Jiao X, Baseler MW, Lane HC, Imamichi T, Chang W. 2022. DAVID: a web server for functional enrichment analysis and functional annotation of gene lists (2021 update). Nucleic Acids Res 50: W216–221.

Shibata H, Chijiwa T, Oda-Ueda N, Nakamura H, Yamaguchi K, Hattori S, Matsubara K, Matsuda Y, Yamashita A, Isomoto A et al. 2018. The habu genome reveals accelerated evolution of venom protein genes. Sci Rep 8: 11300.

Simpson JT, Wong K, Jackman SD, Schein JE, Jones SJM, Birol İ. 2009. ABySS: A parallel assembler for short read sequence data. Genome Res 19: 1117–1123.

Sims JK, Wade PA. 2011. SnapShot: Chromatin remodeling: CHD. Cell 144: 626–626.e621.

Skejić J, Hodgson WC. 2013. Population divergence in venom bioactivities of elapid snake *Pseudonaja textilis*: role of procoagulant proteins in rapid rodent prey incapacitation. PloS One 8: e63988.

Skejić J, Steer DL, Dunstan N, Hodgson WC. 2015. Label-Free (XIC) quantification of venom procoagulant and neurotoxin expression in related Australian elapid snakes gives insight into venom toxicity evolution. J Proteome Res 14: 4896–4906.

Su MJ, Coulter AR, Sutherland SK, Chang CC. 1983. The presynaptic neuromuscular blocking effect and phospholipase A2 activity of textilotoxin, a potent toxin isolated from the venom of the Australian brown snake, *Pseudonaja textilis*. Toxicon 21: 143–151.

Subramanian A, Tamayo P, Mootha VK, Mukherjee S, Ebert BL, Gillette MA, Paulovich A, Pomeroy SL, Golub TR, Lander ES et al. 2005. Gene set enrichment analysis: A knowledge-based approach for interpreting genome-wide expression profiles. Proc Natl Acad Sci USA 102: 15545–15550.

Suryamohan K, Krishnankutty SP, Guillory J, Jevit M, Schroder MS, Wu M, Kuriakose B, Mathew OK, Perumal RC, Koludarov I et al. 2020. The Indian cobra reference genome and transcriptome enables comprehensive identification of venom toxins. Nat Genet 52: 106–117.

Sutherland SK. 1992. Deaths from snake bite in Australia, 1981–1991. Med J Aust 157: 740–746.

Tamiya T, Fujimi TJ. 2006. Molecular evolution of toxin genes in Elapidae snakes. Mol Divers 10: 529–543.

Tasoulis T, Isbister G. 2017. A review and database of snake venom proteomes. Toxins 9: 290.

The UniProt C. 2021. UniProt: the universal protein knowledgebase in 2021. Nucleic Acids Res 49: D480–D489.

Thompson BA, Tremblay V, Lin G, Bochar DA. 2008. CHD8 is an ATP-dependent chromatin remodeling factor that regulates beta-catenin target genes. Mol Cell Biol 28: 3894–3904.

Tuteja G, Kaestner KH. 2007. SnapShot:Forkhead Transcription Factors I. Cell 130: 1160.e1161–1160.e1162.

Tyler MI, Barnett D, Nicholson P, Spence I, Howden MEH. 1987a. Studies on the subunit structure of textilotoxin, a potent neurotoxin from the venom of the Australian common brown snake (*Pseudonaja textilis*). Biochim et Biophys Acta - Prot Struct Mol Enzymol 915: 210–216.

Tyler MI, Spence I, Barnett D, Howden ME. 1987b. Pseudonajatoxin b: unusual amino acid sequence of a lethal neurotoxin from the venom of the Australian common brown snake, *Pseudonaja textilis*. Eur J Biochem 166: 139–143.

van Thiel J, Alonso LL, Slagboom J, Dunstan N, Wouters RM, Modahl CM, Vonk FJ, Jackson TNW, Kool J. 2023. Highly evolvable: Investigating interspecific and intraspecific venom variation in taipans (*Oxyuranus* spp.) and brown snakes (*Pseudonaja* spp.). Toxins 15: 74.

Viala V, Hildebrand D, Riedner M, Mieco Fucase T, Sciani J, Arni R, Schlüter H, Betzel C, Mirtschin P, Dunstan N et al. 2015. Venomics of the Australian eastern brown snake (*Pseudonaja textilis*): Detection of new venom proteins and splicing variants. Toxicon 107: 252–65.

Viscarra JA, Wang Y, Nguyen HP, Choi YG, Sul HS. 2020. Histone demethylase JMJD1C is phosphorylated by mTOR to activate de novo lipogenesis. Nature Com 11: 796.

Vogel CW, Smith CA, Müller-Eberhard HJ. 1984. Cobra venom factor: structural homology with the third component of human complement. Journal Immun 133: 3235.

Vonk FJ, Admiraal JF, Jackson K, Reshef R, de Bakker MA, Vanderschoot K, van den Berge I, van Atten M, Burgerhout E, Beck A, et al. 2008. Evolutionary origin and development of snake fangs. Nature 454: 630–633.

Vonk FJ, Casewell NR, Henkel CV, Heimberg AM, Jansen HJ, McCleary RJ, Kerkkamp HM, Vos RA, Guerreiro I, Calvete JJ et al. 2013. The king cobra genome reveals dynamic gene evolution and adaptation in the snake venom system. Proc Natl Acad Sci USA 110: 20651–20656.

Wang X, Serrano Martinez P, Terpstra JH, Shaalan A, Proctor GB, Spijkervet FKL, Vissink A, Bootsma H, Kroese FGM, Coppes RP et al. 2021. β-Adrenergic signaling induces Notch-mediated salivary gland progenitor cell control. Stem Cell Rep 16: 2813–2824.

Watanabe S, Radman-Livaja M, Rando OJ, Peterson CL. 2013. A histone acetylation switch regulates H2A.Z deposition by the SWR-C remodeling enzyme. Science 340: 195–199.

White J. 2009. Envenomation, prevention and treatment in Australia. CRC Press, Boca Raton.

Williams V, White J. 1992. Variation in the composition of the venom from a single specimen of *Pseudonaja textilis* (common brown snake) over one year. Toxicon 30: 202–206.

Wong MM, Cox LK, Chrivia JC. 2007. The chromatin remodeling protein, SRCAP, is critical for deposition of the histone variant H2A.Z at promoters. J Biol Chem 282: 26132–26139.

Yamanouye N, Britto LR, Carneiro SM, Markus RP. 1997. Control of venom production and secretion by sympathetic outflow in the snake *Bothrops jararaca*. J Exp Biol 200: 2547–2556.

Yang C, Bolotin E, Jiang T, Sladek FM, Martinez E. 2007. Prevalence of the initiator over the TATA box in human and yeast genes and identification of DNA motifs enriched in human TATA-less core promoters. Gene 389: 52–65.

Yin X, Guo S, Gao J, Luo L, Liao X, Li M, Su H, Huang Z, Xu J, Pei J et al. 2020. Kinetic analysis of effects of temperature and time on the regulation of venom expression in *Bungarus multicinctus*. Sci Rep 10: 14142.

Zancolli G, Reijnders M, Waterhouse RM, Robinson-Rechavi M. 2022. Convergent evolution of venom gland transcriptomes across Metazoa. Proc Natl Acad Sci USA 119: e2111392119.

Zaret KS, Carroll JS. 2011. Pioneer transcription factors: establishing competence for gene expression. Genes Dev 25: 2227–2241.

Zhang J, Kobert K, T. F, Stamatakis A. 2014. PEAR: A fast and accurate Illumina Paired-End reAd mergeR. Bioinformatics 30: 614–620.

Zhang K, Shen X, Wu J, Sakaki K, Saunders T, Rutkowski DT, Back SH, Kaufman RJ. 2006. Endoplasmic reticulum stress activates cleavage of CREBH to induce a systemic inflammatory response. Cell 124: 587–599.

Zhang P, Lee H, Brunzelle JS, Couture JF. 2012. The plasticity of WDR5 peptide-binding cleft enables the binding of the SET1 family of histone methyltransferases. Nucleic Acids Res 40: 4237–4246.

Zhang ZY, Lv Y, Wu W, Yan C, Tang CY, Peng C, Li JT. 2022. The structural and functional divergence of a neglected three-finger toxin subfamily in lethal elapids. Cell Rep 40: 111079.

