## Supplemental Figures for "Distinct regulatory networks control toxin gene expression in elapid and viperid snakes"

### Supplemental Figure 1

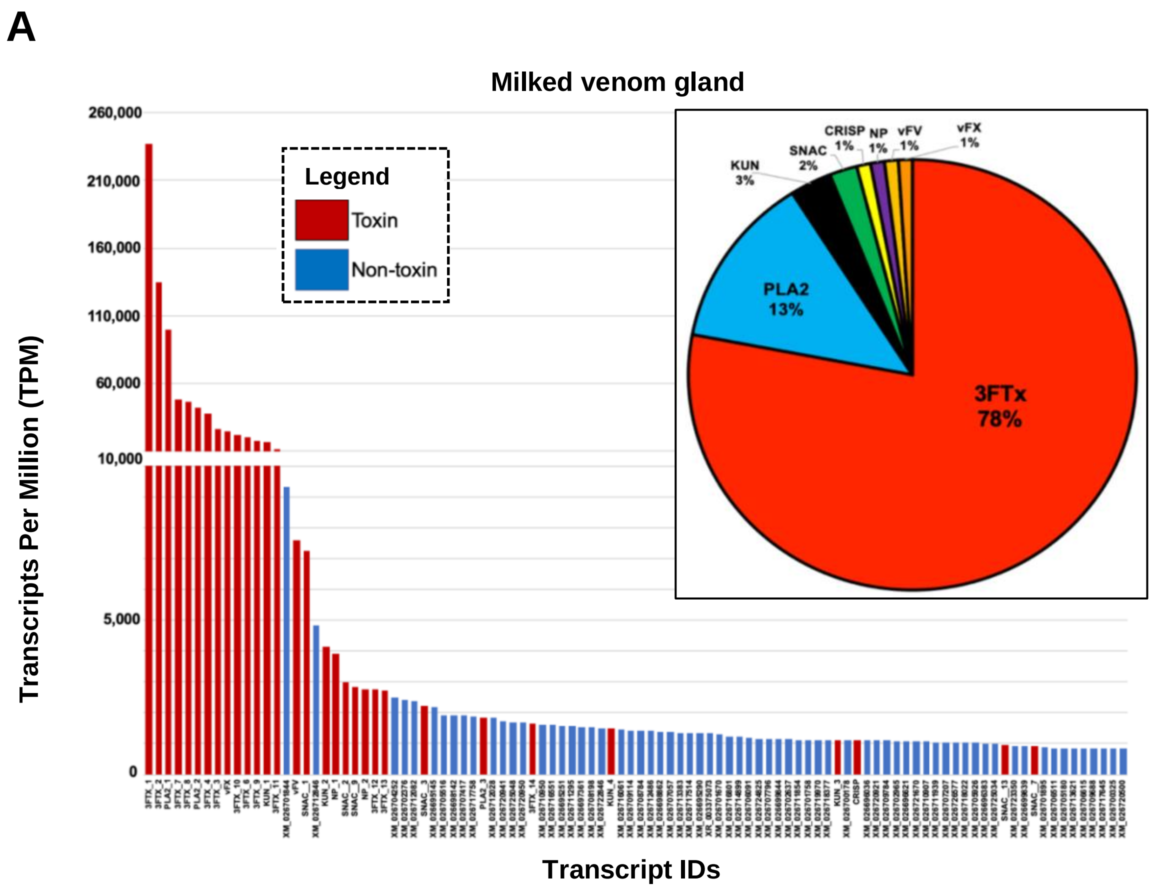


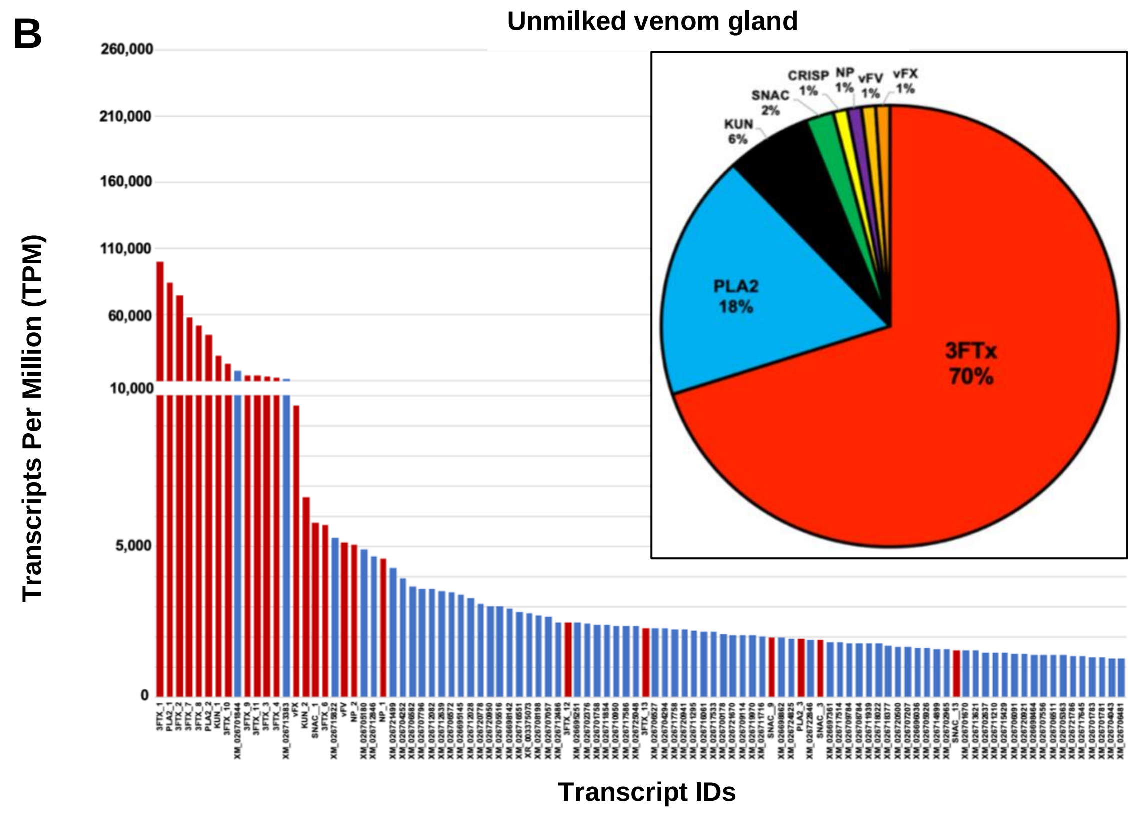

**Supplemental Figure 1.** Toxin and non-toxin expression in a *Pseudonaja textilis* venom gland 96 hours post milking in comparison to an unmilked venom gland. Toxin transcripts (red bars with transcript identities determine from *de novo* assembled venom gland transcriptomes) are expressed in high abundance (TPM; transcripts per million) in comparison to non-toxin transcripts (blue bars with accession numbers from *P. textilis* genome annotations) in the top 100 expressed transcripts in (A) milked and (B) unmilked venom glands. Pie charts (insets in respective panels) show the percentages of total toxin reads belonging to each toxin superfamily, with toxin superfamilies that made up less than 1% excluded due to low abundance. Toxin identifications are as follows: 3FTx = three-finger toxin, CRISP = cysteine-rich secretory protein; KUN = Kunitz serine proteinase inhibitor; NP = natriuretic peptide; PLA2 = phospholipase $A_2$; SNAC = snake C-type lectin; vFA = venom factor V (pseutarin C non-catalytic subunit); vFX = venom factor X (pseutarin C catalytic subunit).

### Supplemental Figure 2

#### A Three-finger toxins

##### Long-chain like

```
3FTx_1                        LICYLDFS-VPHTCAPGEKLCYTRTWNDG----RGTRIERGCAATCPIPKKPEIHVTCCSTDRCNPHPKEKPH
Pseudonaja_LC                 LICYLDFS-VPHTCAPGEKLCYTRTWNDG----RGTRIERGCAATCPIPKKPEIHVTCCSTDRCNPHPKQKPH
```

##### Long-chain

```
Notechis                      LICYMGPK-TPRTCPRGQNLCYTKTWCDAFCSSRGKVVELGCAATCPIA-KSYEDVTCCSTDNCNPFPVRPRPHP-------------
Oxyuranus                     RRCFITPDVRSERCPPGQEVCYTKTWCDGFCGSRGKRVDLGCAATCPTPKKKGIDIICCSKDNCNTFPKWP-----------------
3FTx_3                        RTCFITPDVKSKPCPPGEEVCYTKTWCDGFCGIRGKRVDLGCAATCPTPKKTGIDIICCSTDDCNTFPLRP------RGRLSSIKDHP
3FTx_6*                       RTCFITPDVKSKPCPPGQEVCYTKTWCDGFCGIRGKRVDLGCAATCPTPKKTGIDIICCSTDDCNTFPLRP------RGRLSSIKDHP
Pseudonajatoxinb_homolog*     RTCFITPDVKSKPCPPGQEVCYTKTWCDGFCGIRGKRVDLGCAATCPTPKKTGIDIICCSTDDCNTFPLRP------RGRLSSIKDHP
3FTx_5                        RTCFITPDVKSKPCPPGQEVCYTETWCDGVCGIRGKRVDLGCAATCPTPKKTGIDIICCSTDDCNTFPLRP------RGRLSSIKDHP
3FTx_2                        RTCFITPDVKSKPCPPGQEVCYTETWCDGFCGIRGKRVELGCAATCPTPKKTGIDIICCSTDDCNTFPLRP------RGRLSSIKDHP
Pseudonajatoxinb              RTCFITPDVKSKPCPPGQEVCYTETWCDGFCGIRGKRVELGCAATCPTPKKTGIDIQCCSTDDCNTFPLRP-----------------
Drysdalia                     RKCYKTHPYKSEPCAPGENLCYTKTWCDFRCSQLGKAVELGCAATCPTT-KPYEEVTCCSTDDCNRFPNWERPRPRPRGLLSSIMDHP
Austrelaps                    FSCYKTPDVKSEPCAPGENLCYTKTWCDRFCSIRGKVIELGCAATCPPA-EPRKDITCCSTDNCNPHPAH------------------
3FTx_4                        RTCFKTPYVKSEPCPPGQEVCYTKTWCDRFCSIRGRVIELGCAATCPPA-GPKEDVTCCSTDNCNTHP--------------------
Demansia                      RTCLKTPEVKSEPCPPGQEVCYTKAWCDRMCSFRGKVIELGCAATCPRQ-EPGKEITCCSTDDCNTHP--------------------
```

##### Short-chain

```
3FTx_9                        LICHDSENLDDHVVCKEDETMCYQYTFVPFRDFEVVARGCSP-SCPEEEDAVCCSTDLCNK
Oxyuranus                     LTC--YMNPSGTMVCKEHETMCYRLIVWTFQYHVLYLKGCSS-SCPGGNNCACCSTDLCNN
3FTx_13                       LTC--YNTLGGTVVCKPHETICYEHTFCPFPNRVIFLRGCST-SCPGGNNPVCCSTDLCNL
3FTx_14                       LTC--YKGYHDTVVCKPHETICYEHTFCPFPNRVIFLRGCST-SCPGGNNPVCCSTDLCNL
P.textilis_SC7                LTC--YKRYFDTVVCKPQETICYRYIIPATHGNAITYRGCST-SCPSGIRLVCCSTDLCNK
3FTx_12                       LTC--NKSYYDTVVCKPHETICYRYHVPATHGNVITVRGCST-SCPGGNNPVCCSTDLCNL
P.textilis_SC6                LTC--YKSLSGTVVCKPHETICYRRLIPATHGNAIIDRGCST-SCPGGNRPVCCSTDLCNK
3FTx_11                       LTC--YKGYHDTVVCKPHETICYRYALPATHGNAVTLRGCGT-SCPEGIRPVCCSTDLCNK
P.textilis_SC8                LTC--YKGYHDTVVCKPHETICYEYFIPATHGNVITTRGCST-SCPGGIRPVCCSTDLCNN
3FTx_7**                      LTC--YKGYHDTVVCKPHETICYEYFIPATHGNAILARGCGT-SCPGGIRPVCCRTDLCNK
P.textilis_SC1/5**            LTC--YKGYHDTVVCKPHETICYEYFIPATHGNAILARGCGT-SCPGGIRPVCCRTDLCNK
P.textilis_SC4                LTC--YKGYHDTVVCKPHETICYEYFIPATHGNAILARGCGT-SCPGGIRPVCCRTDLSNK
P.textilis_SC3                LTC--YKGYHDTVVCKPHETICYRYLVPATHGNAIPARGCGT-SCPGGNHPVCCSTDLCNK
3FTx_8***                     LTC--YKGYHDTVVCKPHETICYRYLIPATHGNAIPARGCGT-SCPGGNHPVCCSTDLCNK
P.textilis_SC2***             LTC--YKGYHDTVVCKPHETICYRYLIPATHGNAIPARGCGT-SCPGGNHPVCCSTDLCNK
3FTx_10                       LTC--YKGYHDTVVCKPHETICYRYLIPATHGNAILARGCGT-SCPGGNHPVCCSTDLCNK
```

#### B Cysteine-rich secretory proteins

```
Pseudechis_australis          TADFASESSNKKNYQKEIVDKHNALRRSVKPTARNMLQMKWNSRAAQNAKRWANRCTFAHSPPNKRTVGKLRCGENIFMSSQPFPWSGVV
Pseudechis                    TVDFASESSNKKNYQKEIVDKHNALRRSVKPTARNMLQMKWNSHAAQNAKRWADRCTFAHSPPNTRTVGKLRCGENIFMSSQPFPWSGVV
CRISP_1*                      TVDFASESSNKNDYQKEIVDKHNDLRRSVKPTARNMLQMKWNSRAAQNAKRWANRCTFAHSPPYTRTVGKLRCGENIFMSSQPFAWSGVV
Pseudonaja_textilis*          TVDFASESSNKNDYQKEIVDKHNDLRRSVKPTARNMLQMKWNSRAAQNAKRWANRCTFAHSPPYTRTVGKLRCGENIFMSSQPFAWSGVV
Oxyuranus                     TVDFASESSNKKDYRKEIVDKHNDLRRSVKPTARNMLQMKWNSRAAQNAKRWANRCTFAHSPPYTRTVGKLRCGENIFMSSQPFAWSGVV
Drysdalia                     TVDFASESSNKKDYRKEIVDKHNALRRSVKPTARNMLQMEWNSHAAQNAKRWADRCTFAHSPPHTRTVGQLRCGENIFMSSQPFAWSGVV
Notechis                      TVDFASESSNKKDYQKEIVDKHNALRRSVKPTARNMLRMEWNSHAAQNAKRWADRCTFAHSPPHTRTVGKLRCGENIFMSSQPFAWSGVV
Austrelaps                    TVDFASESSNKKDYRKEIVDKHNALRRSVKPTARNMLRMEWNSRAAQNAKRWADRCTFAHSPPHTRTVGKLRCGENIFMSTQPFAWSGVV
```

```
Pseudechis_australis      QAWYDEIKNFVYGIGAKPPGSVIGHYTQVVWYKSYLIGCASAKCSSSKYLYVCQYCPAGNIRGSIATPYKSGPPCADCPSACVNKLCTNP
Pseudechis                QAWYDEIKNFVYGIGAKPPGSVIGHYTQVVWYKSHLIGCASAKCSSSKYLYVCQYCPAGNIRGSIATPYKSGPPCADCPSACVNRLCTNP
CRISP_1*                  QAWYDEVKKFVYGIGAKPPSSVTGHYTQVVWYKSHLLGCASAKCSSTKYLYVCQYCPAGNIVGSIATPYKSGPPCGDCPSACDNGLCTNP
Pseudonaja_textilis*      QAWYDEVKKFVYGIGAKPPSSVTGHYTQVVWYKSHLLGCASAKCSSTKYLYVCQYCPAGNIVGSIATPYKSGPPCGDCPSACDNGLCTNP
Oxyuranus                 QAWYDEVKKFVYGIGAKPPSSVIGHYTQVVWYKSHLLGCASAKCSSTKYLYVCQYCPAGNIIGSIATPYKSGPPCGDCPSACDNGLCTNP
Drysdalia                 QAWYDEVKKFVYGIGAKPPGSVIGHYTQVVWYKSHLLGCASAKCSSTKYLYVCQYCPAGNIRGSIATPYKSGPTCGDCPSACVNGLCTNP
Notechis                  QAWYDEVKKFVYGIGAKPPGSVIGHYTQVVWYKSHLLGCASAKCSSTKYLYVCQYCPAGNIRGSIATPYKSGPTCGDCPSACVNGLCTNP
Austrelaps                QAWYDEVKKFVYGIGAKPPGSVIGHYTQVVWYKSHLLGCASAKCSSTKYLYVCQYCPAGNIRGSIATPYKSGPACGDCPSACVNGLCTNP

Pseudechis_australis      CKRNNDFSNCKSLAKKSKCQTEWIKKKCPASCFCHNKII
Pseudechis                CNYNNDFSNCKSLAKKSKCQTEWIKKKCPASCFCHNKII
CRISP_1*                  CKHNDDLSNCKTLVKKHKCQTEWIKSKCPATCFCRTEII
Pseudonaja_textilis*      CKHNDDLSNCKTLVKKHKCQTEWIKSKCPATCFCRTEII
Oxyuranus                 CKHNDDLSNCKPLAKKSKCQTEWIKSKCPATCFCRTEII
Drysdalia                 CKYEDAFTNCNELAKETKCKTEWIKSKCPATCFCHTEII
Notechis                  CKYEDDFSNCKALAKNSKCQTEWIKSKCPAACFCHNKII
Austrelaps                CKYEDAFTNCKALAKKTKCKTEWIKSKCPATCFCHNKII
```

#### C Kunitz-type serine protease inhibitors

```
KUN_3                     KDRPKFCELPADIGPCDDFTGAFHYSPREHECIEFIYGGCKGNANNFNTQEECESACAA-
Textilinin-6              KDRPKFCELPADIGPCDDFTGAFHYSPREHECIEFIYGGCKGNANNFNTQEECESTCAA-
Scutellin-3               KDRPKFCELPADIGPCEDFTGAFHYSPREHECIEFIYGGCKGNANNFNTLEECESACAA-
Microlepidin-3            KDRPKFCELPADIGPCEDFTGAFHYSPREHECIEFIYGGCEGNANNFNTLEECESACAA-
KUN_4                     KDRPKFCELLPDTGPCDDFTGAFHYSTRDRECIEFIYGGCGGNANKFNTLEECESTCARK
Textilinin-5              KDRPKFCELLPDTGSCEDFTGAFHYSTRDRECIEFIYGGCGGNANNFITKEECESTCAA-
Textilinin-7              KDRPKFCELLPDTGSCEDFTGAFHYSTRDRECIEFIYGGCGGNANNFKTLEECESTCAA-
Textilinin-2              KDRPELCELPPDTGPCRVRFPSFYYNPDEQKCLEFIYGGCEGNANNFITKEECESTCAA-
KUN_1*                    KDRPDFCELPADTGPCRVRFPSFYYNPDEKKCLEFIYGGCEGNANNFITKEECESTCAA-
Textilinin-1*             KDRPDFCELPADTGPCRVRFPSFYYNPDEKKCLEFIYGGCEGNANNFITKEECESTCAA-
Mulgin-3                  KDRPDFCELPADTGPCRVGFPSFYYNPDEKKCLEFIYGGCQGNANNFITKEECESTCAA-
Textilinin-3              KDRPNFCKLPAETGRCNAKIPRFYYNPRQHQCIEFLYGGCGGNANNFKTIKECESTCAA-
KUN_2                     KDRPEFCELPADTGSCKGNVPRFYYNADHHQCLKFIYGGCGGNANNFKTIEECKSTCAA-
Textilinin-4              KDHPKFCELPADTGSCKGNVPRFYYNADHHQCLKFIYGGCGGNANNFKTIEECKSTCAA-
```

#### D Phospholipase $A_2$

```
textilotoxin_C            ARIPPLPLNLIQFSNMIKCTIPGSQPLLDYANYGCYCGPGNNGTPVDDVDRCCQAHDECYDEASNHG-CY-----PELTLYDYYCDTGV
textilotoxin_A            SDIPPLPLNLVQFSYLIRCANKYKRPGWHYANYGCYCGSGGRGTPVDDVDRCCQAHDKCYEDAEKLG-CY-----PKWTTYYYYCGANG
textilotoxin_B            --------DLVEFGFMIRCANRNSQPAWQYMDYGCYCGKRGSGTPVDDVDRCCQTHNECYDEAAKIPGCK-----PKWTFYFYQCGSGS
textilotoxin_D            -SIPRPSLNIMLFGNMIQCTIPCEQSWLGYLDYGCYCGSGSSGIPVDDVDKCCKTHDECYYKAGQIPGCSVQPNEVFNVDYSYECNEG-
Austrelaps                SNIPLPSLDFEQFGKMIQCTIPCEESCLAYMDYGCYCGPGGSGTPSDELDRCCQTHDNCYAEAGKLPACKAMLSEPYNDTYSYSCIER-
Tropidechis               --IPARPLNLYQFGNMIQCANHGRRPTWHYMDYGCYCGKGGSGTPVDELDRCCQIHDDCYGEAEKLPACNYMMSGPYYNTYSYECNEG-
Notechis                  --------NLYQFGNMIQCANHGRRPTRHYMDYGCYCGKGGSGTPVDELDRCCQTHDDCYGEAEKLPACNYMMSGPYYNTYSYECNEG-
Oxyuranus                 ARIPPLPLSLLNFANLIECANHGTRSALAYADYGCYCGKGGRGTPLDDLDRCCHVHDDCYGEAEKLPACNYLMSSPYFNSYSYKCNEG-
PLA2_1*                   -RIPPLPLSLDDFSNLITCANRGSRSWLDYAHYGCFCGSGGSGTPVDDLDRCCQVHDNCFGDAEKLPACNYLFSGPYWNPYSYKCNEG-
Pseudonaja_1*             -RIPPLPLSLDDFSNLITCANRGSRSWLDYAHYGCFCGSGGSGTPVDDLDRCCQVHDNCFGDAEKLPACNYLFSGPYWNPYSYKCNEG-
PLA2_3                    -RIPPLPLSLVEFRILIKCANHNSRNVLDYADYGCYCGKGGSGTPVDELDRCCQAHDYCYDDAEKLPACNYLFSGPYWNPYSYKCNEG-
PLA2_2                    -RIPPLPLSLVEFRILIKCANHNSRNVLDYADYGCYCGKGGSGTPVDELDRCCQAHDYCYDDAEKLPACNYRFSGPYWNPYSYKCNEG-
Pseudonaja_2              -RIPPLPLSLVEFRILIKCANHNSRNVLDYADYGCYCGKGGSGTPVDELDRCCQAHDYCYDDAEKLPACNYRFSGPYWNPYSYKCNEG-

textilotoxin_C            -PYC-KARTQCQVFVCGCDLAVAKCLAGATYNDENKNINTGER--CQ
textilotoxin_A            -PYC-KTRTKCQRFVCNCDVVAADCFASYPYNRRYWFYSNKKR--CR
textilotoxin_B            QFTCRKSKDVCRNVVCDCDFKAALCLTGARYNSANYNIDIKTH--CR
textilotoxin_D            QLTCNESNNECEMAVCNCDRAAAICFARFPYNKNYWSINTEIH--CR
Austrelaps                QLTCNDDNDECKAFICNCDRAAVICFSGAPYNDSNYDIGTIEH--CK
Tropidechis               ELTCKDNNDECKAFICNCDRTAAICFARTPYNDANWNIDTKTR--C-
Notechis                  ELTCKDNNDECKAFICNCDRTAAICFARAPYNDANWNIDTKTR--CQ
Oxyuranus                 KVTCTDDNDECKAFICNCDRTAAICFAGATYNDENFMISKKRNDICQ
PLA2_1*                   EITCTDDNDECAAFICNCDRTAAICFAGATYNDENFMVTIKKKNICQ
Pseudonaja_1*             EITCTDDNDECAAFICNCDRTAAICFAGATYNDENFMVTIKKKNICQ
PLA2_3                    EITCTDDNDECAAFICNCDRTAAICFAGATYNDENFMVTIKKKNICQ
PLA2_2                    EVTCTDDNDECKAFICNCDRTAAICFAGAPYNDENFMITTKKKNICQ
Pseudonaja_2              EVTCTDDNDECKAFICNCDRTAAICFAGAPYNDENFMITIKKKNICQ
```

#### E Coagulation factor V

```
FV                        AQLREYHIAAQLEDWDYNPQPEELSRLSESDLTFKKIVYREYELDFKQEKPRDELSGLLGPTLRGEVGDILIIYFKNFATQPVSIHPQSAVY
vFV                       AQLREYHIAAQLEDWDYNPQPEELSRLSESDLTFKKIVYREYELDFKQEKPRDALSGLLGPTLRGEVGDSLIIYFKNFATQPVSIHPQSAVY
Pseutarin C               AQLREYHIAAQLEDWDYNPQPEELSRLSESDLTFKKIVYREYELDFKQEEPRDALSGLLGPTLRGEVGDSLIIYFKNFATQPVSIHPQSAVY
Omicarin C                AQLREYRIAAQLEDWDYNPQPEELSRLSESELTFKKIVYREYELDFKQEKPRDELSGLLGPTLRGEVGDILIIYFKNFATQPVSIHPQSAVY
Oscutarin C               AQLREYRLAAQLEDWDYNPQPEELSRLSESDLTFKKIVYREYELDFKQEKPRDELSGLLGPTLRGEVGDSLIIYFKNFATQPVSIHPQSAVY

FV                        NKWSEGSSYSDGTSDVERLDDAVPPGQSFKYVWNITAEIGPKKADPPCLTYAYYSHVNMVRDFNSGLIGALLICKEGSLNANGSQKFFNREY
vFV                       NKWSEGSSYSDGTSDVERLDDAVPPGQSFKYVWNITAEIGPKKADPPCLTYAYYSHVNMVRDFNSGLIGALLICKEGSLNANGSQKFFNREY
Pseutarin C               NKWSEGSSYSDGTSDVERLDDAVPPGQSFKYVWNITAEIGPKKADPPCLTYAYYSHVNMVRDFNSGLIGALLICKEGSLNANGSQKFFNREY
Omicarin C                NKWSEGSSYSDGTSDVERLDDAVPPGQSFKYVWNITAEIGPKKADPPCLTYAYYSHVNMVRDFNSGLIGALLICKEGSLNANGAQKFFNREY
Oscutarin C               NKWSEGSSYSDGTSDVERLDDAVPPGQSFKYVWNITAEIGPKKADPPCLTYAYYSHVNMVRDFNSGLIGALLICKEGSLNADGAQKFFNREY
```

```
FV            VLMFSVFDESKNWYRKPSLQYTINGFANGTLPDVQACAYDHISWHLIGMSSSPEIFSVHFNGQTLEQNHYKVSTINLVGGASVTANMSVSRT
vFV           VLMFSVFDESKNWYRKPSLQYTINGFANGTLPDVQACAYDHISWHLIGMSSSPEIFSVHFNGQTLEQNHYKVSTINLVGGASVTADMSVSRT
Pseutarin C   VLMFSVFDESKNWYRKPSLQYTINGFANGTLPDVQACAYDHISWHLIGMSSSPEIFSVHFNGQTLEQNHYKVSTINLVGGASVTADMSVSRT
Omicarin C    VLMFSVFDESKNWYRKPSLQYTINGFANGTLPDVQACAYDHISWHLIGMSSSPEIFSVHFNGQTLEQNHYKVSTINLVGGASVTANMSVSRT
Oscutarin C   VLMFSVFDESKNWYRKPSLQYTINGFANGTLPDVQACAYDHISWHLIGMSSSPEIFSVHFNGQTLEQNHYKVSTINLVGGASVTANMSVSRT

FV            GKWLISSLVAKHLQAGMYGYLNIKDCGNPDTLTRKLSFRELRRIMNWEYFIAAEEITWDYAPEIPSSVDRRYKAQYLDNFSNFIGKKYKKAV
vFV           GKWLISSLVAKHLQAGMYGYLNIKDCGNPDTLTRKLSFRELMKIKNWEYFIAAEEITWDYAPEIPSSVDRRYKAQYLDNFSNFIGKKYKKAV
Pseutarin C   GKWLISSLVAKHLQAGMYGYLNIKDCGNPDTLTRKLSFRELMKIKNWEYFIAAEEITWDYAPEIPSSVDRRYKAQYLDNFSNFIGKKYKKAV
Omicarin C    GKWLISSLVAKHLQAGMYGYLNIKDCGHPNTLTRKLSFRELRRIMNWEYFIAAEEITWDYAPEIPSSVDRRYKAQYLDNFSNFIGKKYKKAV
Oscutarin C   GKWLISSLVAKHLQAGMYGYLNIKDCGNPDTLTRKLSFRERRRIMKWEYFIAAEEITWDYAPEIPSSVDRRYKAQYLDNFSNFIGKKYKKAV

FV            FRQYKDSNFTKPTYAIWPKERGILGPVIRAKVRDTISIVFKNLASRPYSIYVHGVSVSKDAEGAIYPSDPKENITHGKAVEPGQVYTYKWTV
vFV           FRQYEDGNFTKPTYAIWPKERGILGPVIKAKVRDTVTIVFKNLASRPYSIYVHGVSVSKDAEGAIYPSDPKENITHGKAVEPGQVYTYKWTV
Pseutarin C   FRQYEDGNFTKPTYAIWPKERGILGPVIKAKVRDTVTIVFKNLASRPYSIYVHGVSVSKDAEGAIYPSDPKENITHGKAVEPGQVYTYKWTV
Omicarin C    FRQYEDGNFTKPTYAIWPKERGILGPVIKAKVRDTVTIVFKNLASRPYSIYVHGVSVSKDAEGAIYPSDPKENITHGKAVEPGQVYTYKWTV
Oscutarin C   FRQYEDSNFTKPTYAIWPKERGILGPVIKAKVRDTVTIVFKNLASRPYSIYVHGVSVSKDAEGAVYPSDPKENITHGKAVEPGQVYTYKWTV

FV            LDTDEPTVKDSECITKLYHSAVDMTRDIASGLIGPLLVCKHKALSVKGVQNKADVEQHAVFAVFDENKSWYLEDNIKKYCSNPSTVKKDDPK
vFV           LDTDEPTVKDSECITKLYHSAVDMTRDIASGLIGPLLVCKHKALSVKGVQNKADVEQHAVFAVFDENKSWYLEDNIKKYCSNPSAVKKDDPK
Pseutarin C   LDTDEPTVKDSECITKLYHSAVDMTRDIASGLIGPLLVCKHKALSVKGVQNKADVEQHAVFAVFDENKSWYLEDNIKKYCSNPSAVKKDDPK
Omicarin C    LDTDEPTVKDSECITKLYHSAVDMTRDIASGLIGPLLVCKLKALSVKGVQNKADVEQHAVFAVFDENKSWYLEDNIKKYCSNPSSVKKDDPK
Oscutarin C   LDTDEPTVKDSECITKLYHSAVDMTRDIASGLIGPPLVCKRKALSIRGVQNKADVEQHAVFAVFDENKSWYLEDNIKKYCSNPSSVKKDDPK

FV            FYKSNVMYTLNGYASDRTEVLGFHQSEVVEWHLTSVGTVDEIVPVHLSGHTFLSKGKHQDILNLFPMSGESATVTMDNLGTWLLSSWGSCEM
vFV           FYKSNVMYTLNGYASDRTEVLRFHQSEVVQWHLTSVGTVDEIVPVHLSGHTFLSKGKHQDILNLFPMSGESATVTMDNLGTWLLSSWGSCEM
Pseutarin C   FYKSNVMYTLNGYASDRTEVLRFHQSEVVQWHLTSVGTVDEIVPVHLSGHTFLSKGKHQDILNLFPMSGESATVTMDNLGTWLLSSWGSCEM
Omicarin C    FYKSNVMYTLNGYASDRTEVLGFHQSEVVQWHLTSVGTVDEIVPVHLSGHTFLSKGKHQDILNLFPMSGESATVTMDNLGTWLLSSWGSCEM
Oscutarin C   FYKSNVMYTLNGYASDRTEVWGFHQSEVVEWHLTSVGTVDEIVPVHLSGHTFLSKGKHQDILNLFPMSGESATVTMDNLGTWLLSSWGSCEM

FV            SNGMRLRFLDANYDDEDEGNEEEEEDDGDIFADIFIPPEVVKKKEKDPVNFVSDPESDKIAKELGLLDDEDNQE-ESHNVQTEDDEEQLMIA
vFV           SNGMRLRFLDANYDDEDEGNEEEEEDDGDIFADIFIPSEVVKKKEEVPVNFVPDPESDALAKELGLIDDEGNPIIQPRREQTEDDEEQLMKA
Pseutarin C   SNGMRLRFLDANYDDEDEGNEEEEEDDGDIFADIFIPSEVVKKKEEVPVNFVPDPESDALAKELGLIDDEGNPIIQPRREQTEDDEEQLMKA
Omicarin C    SNGMRLRFLDANYDDEDEGNEEEEEDDGDIFADIFSPPEVVKKKEEVPVNFVPDPESDALAKELGLLDDEDNPE-QSRSEQTEDDEEQLMIA
Oscutarin C   SNGMRLRFLDANYDDEDEGNEEEEEDDGDIFADIFNPPEVVIKKEEVPVNFVPDPESDALAKELGLFDDEDNPK-QSRSEQTEDDEEQLMIA

FV            TMLGFRSFKGSVAEEELNLTALALEEDAHASDPRIDSNSARNPDDIAGRYLRTINRGNKRRYYIAAEEVLWDYSPIGKSQVRSRAAKTTFKK
vFV           SMLGLRSFKGSVAEEELKHTALALEEDAHASDPRIDSNSARNPDDIAGRYLRTINRGNKRRYYIAAEEVLWDYSPIGKSQVRSRAAKTTFKK
Pseutarin C   SMLGLRSFKGSVAEEELKHTALALEEDAHASDPRIDSNSARNPDDIAGRYLRTINRGNKRRYYIAAEEVLWDYSPIGKSQVRSRAAKTTFKK
Omicarin C    SVLGLRSFKGSVAEEELKHTALALEEDAHASDPRIDSNSARNSDDIAGRYLRTINRRNKRRYYIAAEEVLWDYSPIGKSQVRSLPAKTTFKK
Oscutarin C   SMLGLRSFKGSVAEEELKHTALALEEDAHASDPRIDSNSAHNSDDIAGRYLRTINRRNKRRYYIAAEEVLWDYSPIGKSQVRSLPAKTTFKK

FV            AIFRSYLDDTFQTPSTGGEYEKHLGILGPIIRAEVDDVIEVQFRNLASRPYSLHAHGLLYEKSSEGRSYDDKSPELFKKDDAIMPNGTYTYV
vFV           AIFRSYLDDTFQTPSTGGEYEKHLGILGPIIRAEVDDVIEIQFRNLASRPYSLHAHGLLYEKSSEGRSYDDKSPELFKKDDAIMPNGTYTYV
Pseutarin C   AIFRSYLDDTFQTPSTGGEYEKHLGILGPIIRAEVDDVIEIQFRNLASRPYSLHAHGLLYEKSSEGRSYDDKSPELFKKDDAIMPNGTYTYV
Omicarin C    AIFRSYLDDTFQTPSTGGEYEKHLGILGPIIRAEVDDVIEVQFRNLASRPYSLHAHGLLYEKSSEGRSYDDNSPELFKKDDAIMPNGTYTYV
Oscutarin C   AIFRSYLDDTFQTPSTGGEYEKHLGILGPIIRAEVDDVIEVQFRNLASRPYSLHAHGLLYEKSSEGRSYDDNSPELFKKDDAIMPNGTYTYV

FV            WQVPPRSGPTDNTEKCKSWAYYSGVNPEKDIHSGLIGPILICQKGMIDKYNRTIDIREFVLFFMVFDEEKSWYFPKSDKSTRAEKLIGVQS-
vFV           WQVPPRSGPTDNTEKCKSWAYYSGVNPEKDIHSGLIGPILICQKGMIDKYNRTIDIREFVLFFMVFDEEKSWYFPKSDKSTCEEKLIGVQS-
Pseutarin C   WQVPPRSGPTDNTEKCKSWAYYSGVNPEKDIHSGLIGPILICQKGMIDKYNRTIDIREFVLFFMVFDEEKSWYFPKSDKSTCEEKLIGVQS-
Omicarin C    WQVPPRSGPTDNTEKCKSWAYYSGVNPEKDIHSGLIGPILICQKGMIDKYNRTIDIREFVLFFMVFDEEKSWYFPKSDKSTCEEKLIGVQSS
Oscutarin C   WQVPPRSGPTDNTEKCKSWAYYSGVNPEKDIHSGLIGPILICQKGMIDKYNRTIDIREFVLFFMVFDEEKSWYFPKSDKSTCEEKLIGVQS-

FV            RHTFPAINGIPYQLQGLTMYKDENVHWHLLNMGGPKDIHVVNFHGQTFTEEGREDNQLGVLPLLPGTFASIKMKPSKIGTWLLETEVGENQE
vFV           LHTFPAINGIPYQLQGLTMYKDENVHWHLLNMGGPKDIHVVNFHGQTFTEEGREDNQLGVLPLLPGTFASIKMKPSKIGTWLLETEVGENQE
Pseutarin C   LHTFPAINGIPYQLQGLTMYKDENVHWHLLNMGGPKDIHVVNFHGQTFTEEGREDNQLGVLPLLPGTFASIKMKPSKIGTWLLETEVGENQE
Omicarin C    HHTFPAINGIPYQLQGLMMYKDENVHWHLLNMGGPKDIHVVNFHGQTFTEEGREDNQLGVLPLLPGTFASIKMKPSKIGTWLLETEVGENQE
Oscutarin C   RHTFPAINGIPYQLQGLMMYKDENVHWHLLNMGGPKDVHVVNFHGQTFTEEGREDNQLGVLPLLPGTFASIKMKPSKIGTWLLETEVGENQE

FV            RGMQALFTVIDKDCKLPMGLASGIIQDSQISASGHVGYWEPKLARLNNTGKYNAWSIIKKEHEHPWIQIDLQRQVVITGIQTQGAMQLLKHL
vFV           RGMQALFTVIDKDCKLPMGLASGIIQDSQISASGHVGYWEPKLARLNNTAIFNAWSIIKKEHEHPWIQIDLQRQVVITGIQTQGTVQLLQHS
Pseutarin C   RGMQALFTVIDKDCKLPMGLASGIIQDSQISASGHVGYWEPKLARLNNTGKYNAWSIIKKEHEHPWIQIDLQRQVVITGIQTQGTVQLLQHS
Omicarin C    RGMQALFTVIDKDCKLPMGLASGIIQDSQISASGHVEYWEPKLARLNNTGMFNAWSIIKKEHEHPWIQIDLQRQVVITGIQTQGTVQLLKHS
Oscutarin C   RGMQALFTVIDKDCKLPMGLASGIIQDSQISASGHVGYWEPKLARLNNTGMFNAWSIIKKEHEHPWIQIDLQRQVVITGIQTQGTVQLLKHS

FV            YTVEYFFTYSKDGQNWITFKGRHSETQMHFEGNSDGTTVKENHIDPPIIARYIRLHPTKFHNRPTFRIELLGCEVEGCSVPLGMESGAIKNS
vFV           YTVEYFVTYSEDGQNWITFKGRHSETQMHFEGNSDGTTVKENHIDPPIIARYIRLHPTKFYNRPTFRIELLGCEVEGCSVPLGMESGAIKNS
Pseutarin C   YTVEYFVTYSEDGQNWITFKGRHSETQMHFEGNSDGTTVKENHIDPPIIARYIRLHPTKFYNRPTFRIELLGCEVEGCSVPLGMESGAIKNS
Omicarin C    YTVEYFVTYSKDGQNWITFKGRHSETQMHFEGNSDGTTVKENHIDPPIIARYIRLHPTKFYNTPTFRIELLGCEVEGCSVPLGMESGAIKNS
Oscutarin C   YTVEYFVTYSKDGQNWITFKGRHSKTQMHFEGNSDGTTVKENHIDPPIIARYIRLHPTKFYNTPTFRIELLGCEVEGCSVPLGMESGAIKDS

FV            EITASSYKKTWWSSWEPSLARLNLKGRTNAWQPKVNNKDQWLQIDLQHLTKITSIITQGATSMTTSMYVKTFSIHYTDDNSTWKPYLDVRTS
vFV           EITASSYKKTWWSSWEPFLARLNLEGGTNAWQPEVNNKDQWLQIDLQHLTKITSIITQGATSMTTSMYVKTFSIHYTDDNSTWKPYLDVRTS
Pseutarin C   EITASSYKKTWWSSWEPSLARLNLEGGTNAWQPEVNNKDQWLQIDLQHLTKITSIITQGATSMTTSMYVKTFSIHYTDDNSTWKPYLDVRTS
Omicarin C    EITASSYKKTWWSSWEPFLARLNLEGGTNAWQPEVNNKDQWLQIDLQHLTKITSIITQGATSMTTAMYVKTFSIHYTDDNSTWKPYLDVRTS
Oscutarin C   EITASSYKKTWWSSWEPFLARLNLKGRTNAWQPKVNNKDQWLQIDLQHLTKITSIITQGATSMTTSMYVKTFSIHYTDDNSTWKPYLDVRTS

FV            MEKVFTGNINSDGHVKHFFKPPILSRFIRIIPKTWNQYIALRIELFGCEVF
vFV           MEKVFTGNINSDGHVKHFFKPPILSRFIRIIPKTWNQYIALRIELFGCEVF
Pseutarin C   MEKVFTGNINSDGHVKHFFKPPILSRFIRIIPKTWNQYIALRIELFGCEVF
Omicarin C    MEKVFTGNINSDGHVKHFFKPPILSRFIRIIPKTWNQYIALRIELFGCEVF
Oscutarin C   MEKVFTGNINSDGHVKHFFNPPILSRFIRIIPKTWNQYIALRIELFGCEVF
```

#### F Coagulation factor X

```
Omicarin C    NVFLKSKVANRFLQRTKRANSLFEEFRSGNIERECIEERCSKEEAREVFEDDEKTETFWNVYVDGDQCSSNPCHYRGTCKDGIGSYTCTCLF
Oscutarin C   NVFLKSKVANRFLQRTKRANSLYEEFRSGNIERECIEERCSKEEAREVFEDDEKTETFWNVYVDGDQCSSNPCHYRGTCKDGIGSYTCTCLS
FX isoform 2  NVFLKSKVANRFLQRTKRANSLVEEFKSGNIERECIEERCSKEEAREAFEDDEKTETFWNVYVDGDQCSSNPCHYRGICKDGIGSYTCTCLS
```

```
vFX                       NVFLKSKVANRFLQRTKRANSLVEEFKSGNIERECIEERCSKEEAREVFEDDEKTETFWNVYVDGDQCSSNPCHYRGICKDGIGSYTCTCLS
Pseutarin C               NVFLKSKVANRFLQRTKRANSLVEEFKSGNIERECIEERCSKEEAREVFEDDEKTETFWNVYVDGDQCSSNPCHYRGICKDGIGSYTCTCLS
Tr FX                     NVFLKSKVANRFLQRTKRANSLFEEFKAGNIERECIEERCSKEEAREAFEDNEKTETFWNVYVDGDQCSSNPCHYGGTCKDGIGSYTCTCLA
FX isoform 1              NVFLKSKVANRFLQRTKRANSLFEEFKSGNIERECIEERCSKEEAREAFEDDEKTETFWNVYVDGDQCSSNPCHYGGTCKDGIGSYTCTCLA
Porpharin D               NVFLKSKEANRFLQRTKRSNSLFEEFRPGNIERECIEEKCSKEEAREIFKDNEKTEAFWNVYVDGDQCSSNPCHYGGTCKDGIGSYTCTCLP
Notecarin D               NVFLKSKVANRFLQRTKRSNSLFEEIRPGNIERECIEEKCSKEEAREVFEDNEKTETFWNVYVDGDQCSSNPCHYRGTCKDGIGSYTCTCLP
Trocarin D                NVFLKSKVANRFLQRTKRSNSLFEEIRPGNIERECIEEKCSKEEAREVFEDNEKTETFWNVYVDGDQCSSNPCHYRGTCKDGIGSYTCTCLP

Omicarin C                GYEGKNCERVLYKSCRVDNGNCWHFCKPVQNDIQCSCAEGYLLGEDGHSCVAGGNFSCGRNIKTRNKREASLPDF-----------------
Oscutarin C               GYEGKNCERVLYKSCRVDNGNCWHFCKPVQNDIQCSCAEGYLLGEDGHSCVAGGNFSCGRNIKTRNKREASLPDF-----------------
FX isoform 2              GYEGKNCERVLYKSCRVDNGNCWHFCKHVQNDIQCSCAEGYLLGEDGHSCVAGGNFSCGRNIKTRNKREANLPDF-----------------
vFX                       GYEGKNCERVLYKSCRVDNGNCWHFCKSVQNDIQCSCAEGYLLGEDGHSCVAGGNFSCGRNIKTRNKREASLPDF-----------------
Pseutarin C               GYEGKNCERVLYKSCRVDNGNCWHFCKSVQNDIQCSCAEGYLLGEDGHSCVAGGNFSCGRNIKTRNKREASLPDF-----------------
Tr FX                     GYEGKNCQYVLYQSCRVDNGNCWHFCKPVQNEIQCSCAESYLLGDDGYSCVAGGDFSCGRNIKARNKREASLPDFQTDFSDDYDAIDENNFV
FX isoform 1              GYEGKNCEYVLYKSCRVDNGDCWHFCKPVQNGIQCSCAESYLLGEDGHSCVAGGDFSCGRNIKTRNKREANLPDFQTDFSDDYDEIDENNFV
Porpharin D               NYEGKNCEHLLFKSCRFFNGNCWHFCKPVQNDTQCSCAESYRLGDDGHSCVAEGDFSCGRNIKARNKREASLPDF-----------------
Notecarin D               NYEGKNCEKVLYQSCRVDNGNCWHFCKRVQSETQCSCAESYRLGVDGHSCVAEGDFSCGRNIKARNKREASLPDF-----------------
Trocarin D                NYEGKNCEKVLYQSCRVDNGNCWHFCKRVQSETQCSCAESYRLGVDGHSCVAEGDFSCGRNIKARNKREASLPDF-----------------

Omicarin C                ------------VQSQNATLLKKSDNPSPDIRIVNGMDCKLGECPWQAVLVDEKEGVFCGGTILSPIYVLTAAHCINQTEKISVVVGEIDKS
Oscutarin C               ------------VQSQNAILLKKSDNPSPDIRIVNGMDCKLGECPWQAVLVDEKEDAFCGGTILSPIYVLTAAHCINQTKMISVVVGEINIS
FX isoform 2              ------------VQSQNATLLKKSDNPSPDIRIVNGMDCKLGECPWQAALVDEKEGVFCGGTILSPIYVLTAAHCINETETISVVVGEIDKS
vFX                       ------------VQSQNATLLKKSDNPSPDIRIVNGMDCKLGECPWQATLVDEKEGVFCGGTILSPIYVLTAAHCFNETETISVVVGEIDKS
Pseutarin C               ------------VQSHNATLLKKSDNPSPDIRIVNGMDCKLGECPWQAALVDDKKGVFCGGTILSPIYVLTAAHCINETETISVVVGEIDRS
Tr FX                     ETPTNFSGLVPTVQSQNATLLKKSDNPSPDIRVVNGTDCKLGECPWQALLINDQGDGFCGGTILSPIYVLTAAHCINQTKYIRVVVGEIDIS
FX isoform 1              ETPTNFSGLVLTVQSQNATLLKKSDNPSPDIRVVNGTDCKLGECPWQALLLNDEGDGFCGGTILSPIYVLTAAHCINQTKYITVVVGEIDIS
Porpharin D               ------------VQSQNATLLKKSDNPSPDIRIINGMDCKLGECPWQAVLLDKEGDVFCGGTILSPIYVLTAAHCITQSKHISVVVGEIDIS
Notecarin D               ------------VQSQKATLLKKSDNPSPDIRIVNGMDCKLGECPWQAVLINEKGEVFCGGTILSPIHVLTAAHCINQTKSVSVIVGEIDIS
Trocarin D                ------------VQSQKATLLKKSDNPSPDIRIVNGMDCKLGECPWQAVLINEKGEVFCGGTILSPIHVLTAAHCINQTKSVSVIVGEIDIS

Omicarin C                RVETGHLLSVDKIYVHKKFVPPKKGYKFYEKFDLVSYDYDIAIIQMKTPIQFSENVVPACLPTADFANQVLMKQDFGIISGFGRIFEKGPKS
Oscutarin C               RKNPGRLLSVDKIYVHQKFVPPKKGYEFYEKFDLVSYDYDIAILQMKTPIQFSENVVPACLPTADFANQVLMKQDFGIVSGFGRIFEKGPQS
FX isoform 2              RIETGPLLSVDKIYVHKKFVPPQKAY----KFDLAAYDYDIAIIQMKTPIQFSENVVPACLPTADFANQVLMKQDFGIVSGFGRIFEKGPKS
vFX                       RVETGPLLSVDKIYVHKKFVPPKRGYKFYEKFDLVSYDYDIAIIQLKTPIQFSENVVPACLPTADFANQVLMKQDFGIVSGFGHIFERGPKS
Pseutarin C               RAETGPLLSVDKVYVHKKFVPPKKSQEFYEKFDLVSYDYDIAIIQMKTPIQFSENVVPACLPTADFANQVLMKQDFGIVSGFGGIFERGPNS
Tr FX                     RKKTGRLLSVDKIYVHQKFVP-------------STYDYDIALIQMKTPIQFSENVVPACLPTADFANQVLMKQDFGIVSGFGRTRERGQTS
FX isoform 1              SKKTGRLHSVDKIYVHQKFVP-------------ATYDYDIAIIQLKTPIQFSENVVPACLPTADFANQVLMKQNFGIVSGFGRTRERGKTS
Porpharin D               RKETRHLLSVDKAYVHTKFV-------------LATYDYDIAIIQLKTPIQFSENVVPACLPTADFANQVLMKQDFGIISGFGHTRSGGQTS
Notecarin                 RKETRRLLSVDKIYVHTKFVPPNYYY-VHQNFDRVAYDYDIAIIRMKTPIQFSENVVPACLPTADFANEVLMKQDSGIVSGFGRIRFKEPTS
Trocarin D                RKETRRLLSVDKIYVHTKFVPPNYYY-VHQNFDRVAYDYDIAIIRMKTPIQFSENVVPACLPTADFANEVLMKQDSGIVSGFGRIQFKQPTS

Omicarin C                NTLKVLKVPYVDRHTCMVSSESPITPTMFCAGYDTLPRDACQGDSGGPHITAYRDTHFITGIVSWGEGCAKKGKYGIYTKVS
Oscutarin C               KTLKVLKVPYVDRHTCMLSSESPITPTMFCAGYDTLPRDACQGDSGGPHITAYRDTHFITGIVSWGEGCAQTGKYGVYTKVS
FX isoform 2              KTLKVLKVPYVDRHTCMVSSETPITPNMFCAGYDTLPRDACQGDSGGPHTTVYRDTHFITGIVSSGEGCARNGKYGIYTKLS
vFX                       KTLKVLKVPYVDRHTCMLSSNFPITQNMFCAGYDTLPQDACQGDSGGPHITAYRDTHFITGIVSWGEGCAQTGKYGVYTKVS
Pseutarin C               KTLKVLKVPYVDRHTCMLSSNFPITPTMFCAGYDTLPQDACQGDSGGPHITAYRDTHFITGIVSWGEGCARKGRYGIYTKLS
Tr FX                     NTLKVVTLPYVDRHTCMLSSNFPITQNMFCAGYNTLPQDACQGDSGGPHITAYRDTHFITGIISWGEGCAQTGKYGAYTKVS
FX isoform 1              NTLKVVTLPYVDRHTCMLSSNFPITQNMFCAGYDTLPQDACQGDSGGPHITAYRDTHFITGIVSWGEGCAQTGKYGVYTKVS
Porpharin D               NTLKVVTIPYVDRHTCMLSSDFRITPNMFCAGYDTLPRDACQGDSGGPHITAYRDTHFITGIISWGEGCAKKGKYGVYTKVS
Notecarin D               NTLKVITVPYVDRHTCMLSSDFRITQNMFCAGYDTLPQDACQGDSGGPHITAYRDTHFITGIISWGEGCARKGKYGVYTKVS
Trocarin D                NTLKVITVPYVDRHTCMLSSDFRITQNMFCAGYDTLPQDACQGDSGGPHITAYRDTHFITGIISWGEGCARKGKYGVYTKVS

Omicarin C                KFILWIKRIMRQKLPSTESSTGRL
Oscutarin C               KFILWIKRIMRQKLPSTESSTGRL
FX isoform 2              KFIPWIKRIMRQKLPSTESSTGRL
vFX                       KFIPWIKRIMRQKLPSTESSTGRL
Pseutarin C               KFIPWIKRIMRQKLPSTESSTGRL
Tr FX                     RFILWIKRIMRLKLPSTESSTGRL
FX isoform 1              KFILWIKRIIRQKQPSTESSTGRL
Porpharin D               NFIPWIKAVMRKHQPSTESSTGRL
Notecarin                 RFIPWIKKIMSLK-----------
Trocarin D                KFIPWIKKIMSLK-----------
```

**Supplemental Figure 2.** Multiple sequence alignments of all major venom protein sequences deduced from *Pseudonaja textilis* venom gland transcripts. A) Alignment of protein sequences from *P. textilis* three-finger toxin transcripts (3FTx_1 - 3FTx_14; this study), Pseudonaja_LC (A8HDK6; long neurotoxin 1), Pseudonajatoxin b (P13495), Pseudonajatoxin b homolog (Q9W7J5), short-chain neurotoxin 1/ 5 (P.textilis_SC1/5; Q9W7K2), short-chain neurotoxin 2 (P.textilis_SC2; Q9W7K1), short-chain neurotoxin 3 (P.textilis_SC3; Q9W7K0), short-chain neurotoxin 4 (P.textilis_SC4; Q9W7J9), short-chain neurotoxin 6 (P.textilis_SC6; Q9W7J7), short-chain neurotoxin 7 (P.textilis_SC7; Q9W7J6), and short-chain neurotoxin 8 (P.textilis_SC8; A8HDK1). Three-finger toxin sequences (long-chain) from Australian species *Austrelaps superbus* (A8S6A8), *Demansia vestigiata* (A6MFK4), *Drysdalia coronoides* (F8J2D7), *Notechis scutatus* (P01384), and *Oxyuranus microlepidotus* (A7X4Q3). Three-finger toxin sequences (short-chain) from Australian species *D. coronoides* (F8J2G3) and *O. microlepidotus* (A7X4S7).

B) Alignment of protein sequences from cysteine-rich secretory proteins from *P. textilis* (CRISP_1; this study), *P. textilis* pseudechetoxin-like (Q3SB05), and Australian species *A. superbus* (A8S6B6), *D. coronoides* (F8J2D4), *N. scutatus* (Q3SB04), *O. microlepidotus* (Q3SB06), *P. australis* pseudechetoxin (Q8AVA4), and *P. porphyriacus* (Q8AVA3). C) Alignment of protein sequences from Kunitz-type serine protease inhibitors from *P. textilis* venom transcripts (KUN_1-4; this study), textilinin-1 (Q90WA1), textilinin-2 (Q90WA0), textilinin-3 (Q90W99), textilinin-4 (Q90W98), textilinin-5 (Q90W97), textilinin-6 (Q90W96), textilinin-7 (B5L5Q1), and Australian species *O. microlepidotus* (Microlepidin-3; B5KL27), *O. scutellatus* (Scutellin-3; B5KL29) and *Pseudechis australis* (Mulgin-3; Q6ITB9). D) Alignment of protein sequences from *P. textilis* phospholipase $A_2$ ($PLA_2$) transcripts (PLA2_1-3; this study), textilotoxin subunit A (P23026), textilotoxin subunit B (P23027), textilotoxin subunit C (P30811), textilotoxin subunit D (P23028), *P. textilis* acidic phospholipase $A_2$ 1 (Pseudonaja_1; Q9W7J4), *P. textilis* acidic phospholipase $A_2$ 2 (Pseudonaja_2; Q9W7J3), and $PLA_2$s from Australian species *A. superbus* (Q9PUG7), *N. scutatus* (Q9PSN5), *Oxyuranus scutellatus* (Q4VRI5), and *Tropidechis carinatus* (Q45Z25). E) Alignment of protein sequences from pseutarin C venom factor V (vFV; this study), characterized pseutarin C venom factor V (Q7SZN0), *P. textilis* coagulation factor V (Q593B6), and non-catalytic subunits of prothrombin activators in *O. microlepidotus* (omicarin C; Q58L90) and *O. scutellatus* (oscutarin C; Q58L91). F) Alignment of protein sequences from pseutarin C venom factor X (vFX; this study), characterized pseutarin C venom factor X (Q56VR3), *P. textilis* coagulation factor X isoform 1 (FX isoform 1; Q1L659), *P. textilis* coagulation factor X isoform 2 (FX isoform 2; Q1L658), *T. carinatus* coagulation factor X (Tr FX; Q4QXT9) and catalytic subunits of prothrombin activators in *N. scutatus* (Notecarin D1; P82807), *O. microlepidotus* (omicarin C; Q58L95), *O. scutellatus* (oscutarin C; Q58L96), *Pseudechis porphyriacus* (porpharin D; Q58L93), and *T. carinatus* (trocarin D; P81428).

#### Supplemental Figure 3

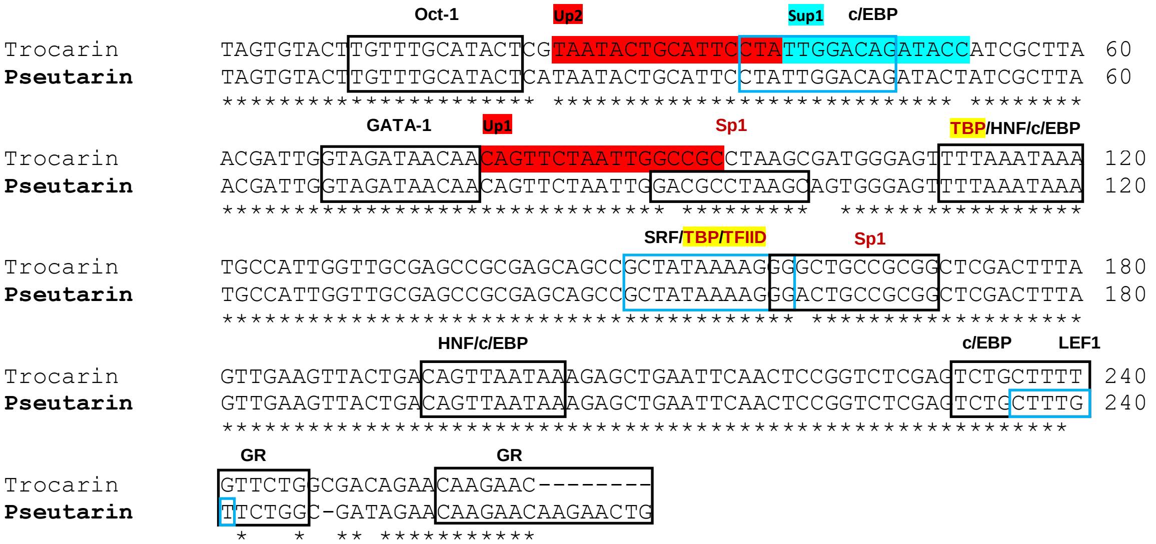


**Supplemental Figure 3.** Alignments of the two *VERSE* core promoter sequences from pseutarin C (*Pseudonaja textilis*) and trocarin D (*Tropidechis carinatus*) with predicted *cis*-regulatory elements. Three regulatory regions have been identified in this *VERSE* core promoter, two that upregulate venom factor X expression (Up1 and Up2, highlighted in red) and one that suppresses expression (Sup1, highlighted in teal) (Kwong et al. 2009). *Cis*-regulatory element (CRE) predictions were completed with the online server AliBaba2.1 using the TRANSFAC 4.0 database. *Trans*-regulatory factors that bind to CREs that were upregulated more than 10-fold in the venom gland of *P. textilis* after venom milking are colored in red.

#### Supplemental Figure 4

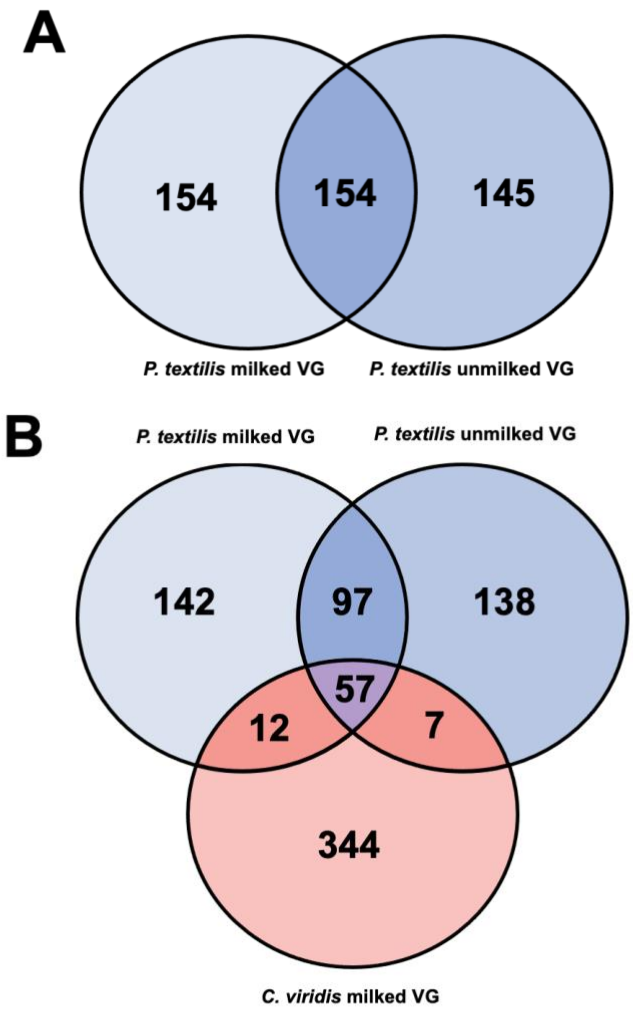


**Supplemental Figure 4.** Conservation of microRNAs between milked and unmilked *Pseudonaja textilis* venom glands and a milked *Crotalus viridis* venom gland. (A) There were approximately equal numbers of common and unique mature miRNA sequences in the milked and unmilked *P. textilis* venom glands. (B) Only 76 miRNAs were common to both *P. textilis* venom glands and the *Crotalus viridis* venom gland. VG = venom gland.

### Supplemental Figure 5

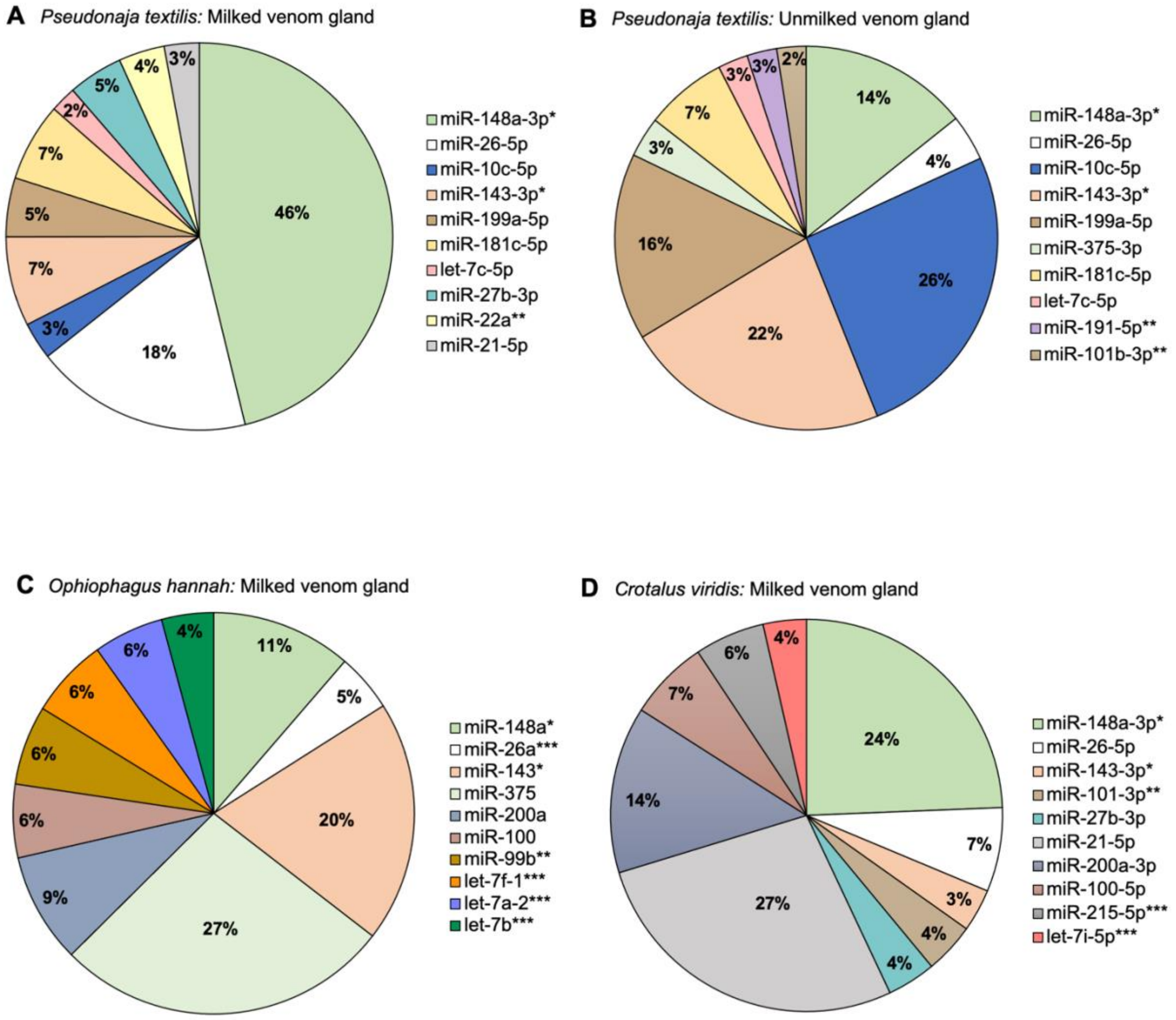


**Supplemental Figure 5.** Abundances of the top ten expressed miRNAs in snake venom glands. Shown are the expression percentages for each of the top ten miRNAs in the (A) *Pseudonaja textilis* milked venom gland, (B) *P. textilis* unmilked venom gland, (C) *Ophiophagus hannah* milked venom gland, and (D) *Crotalus viridis* milked venom gland. * = miRNAs that are in the top ten most abundant miRNAs for all species, ** = miRNAs shared between species, but not in the top ten for all, and *** = miRNAs that are species-specific.

#### Supplemental Figure 6

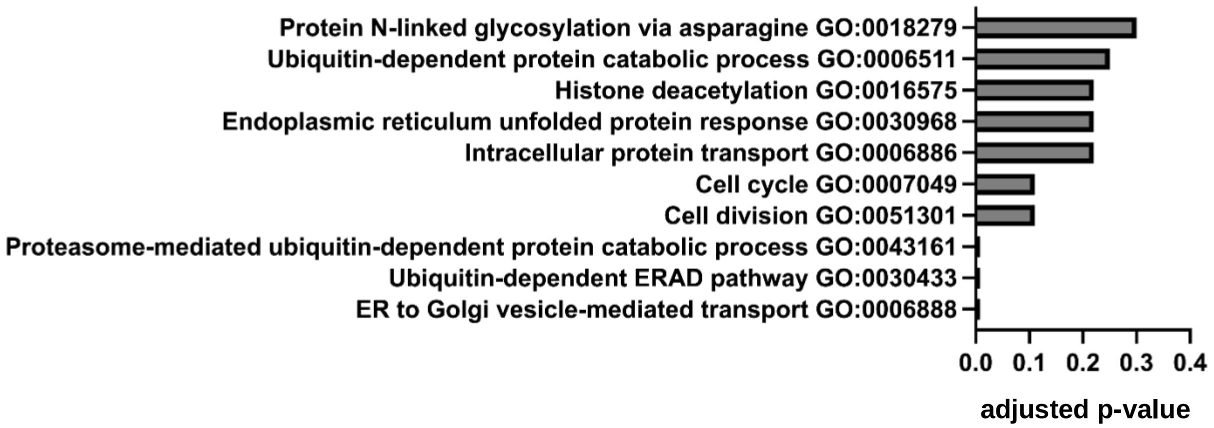


**Supplemental Figure 6.** Biological processes associated with the transcripts targeted by *Pte-miR-1*. Analysis was completed using DAVID Bioinformatics Resources (Huang da et al. 2009b; Sherman et al. 2022) and Benjamini-Hochberg adjusted p-values used for identifying levels of significance for each biological process.
