## Supplemental Tables for "Distinct regulatory networks control toxin gene expression in elapid and viperid snakes"

**Supplemental Table 1.** ***De novo*** **assembled venom gland transcriptomes for** ***Pseudonaja textilis.***

| Gland | Total reads | Assembly quality reads (phred > 30) | Genome alignment % | ABySS contig # & average length | GG-Trinity contig # & average length | Extender contig # & average length | Nonredundant contig # & average length* | Expressed contigs (≥1 TPM) | Expressed contigs average length |
|---|---|---|---|---|---|---|---|---|---|
| Milked | 75,175,149 | 45,740,915 | 58.87% | 412,981<br>469 bp | 207,695<br>764 bp | 2,657<br>675 bp | 74,133<br>941 bp | 28,569 | 1,479 bp |
| Unmilked | 62,074,532 | 37,583,615 | 62.51% | | 152,263<br>726 bp | 3,251<br>688 bp | | | |

*Coding and minimum of 150 bp

**Supplemental Table 2. Gene expression data for** ***Pseudonaja textilis*** **from milked and unmilked venom glands. [EXCEL]**

**Supplemental Table 3. Venom protein isoform diversity in** ***Pseudonaja textilis.***

| Toxin | Targeted toxin work<br><br>cDNA libraries or proteomes | Reeks et al., 2015<br><br>Venom proteome (QLD) | Reeks et al. 2015*<br><br>Venom gland transcriptome (QLD) | Skejic et al., 2013<br><br>Venom proteome (QLD) | Skejic et al., 2013<br><br>Venom proteome (SA) | Birrell et al., 2006<br><br>Venom proteome (SA) | Viala et al., 2015<br><br>Venom proteome (SA) | Viala et al., 2015<br><br>Venom gland transcriptome (SA) | McCleary et al., 2016<br><br>Venom proteome (SA) | This paper<br><br>Venom gland transcriptome (SA) |
|---|---|---|---|---|---|---|---|---|---|---|
| **3FTxs** | 10-11 (Gong et al. 2000; Gong | 8 | 11 | 1 | 8 | n.s. | 5 | 24 | 5-13 | 15 |

| | | | | | | | | | | |
|---|---|---|---|---|---|---|---|---|---|---|
| | et al. 2001; Skejić et al. 2015) | | | | | | | | | |
| **5’-NUC** | | n.d. | n.s. | n.d. | n.d. | n.d. | n.d. | n.d. | 1 | 1 |
| **AChE** | | n.d. | n.s. | n.d. | n.d. | n.d. | n.d. | n.d. | n.d. | 1 |
| **CRISP** | 1 (Pierre et al. 2005) | 1 | n.s. | n.d.. | 1 | 1 | 4 | 1 | 1 | 1 |
| **CVF-like** | | n.d. | n.s. | n.d. | n.d. | n.d. | n.d. | 1 | n.d. | 1 |
| **Cystatin** | | n.d. | n.s. | n.d. | n.d. | n.d. | n.d. | n.d. | n.s. | 3 |
| **Factor Va** | 1 (Rao et al. 2003; Skejić et al. 2015) | 1 | n.s. | 1 | 1 | n.s. | 5 | 2 | 1-3 | 1 |
| **Factor X** | 2 (Skejić et al. 2015) | 1 | n.s. | 2 | 4 | n.s. | 1 | 2 | 1-2 | 1 |
| **HYAL** | | n.d. | n.s. | n.d.. | n.d. | n.d. | n.d. | 2 | n.d. | 1 |
| **KUN** | 6 (Filippovich et al. 2002) | 6 | 7 | 2 | 4 | n.s. | 3 | 1 | 1-5 | 4 |
| **LAAO** | | n.d. | n.d. | n.d. | n.d. | n.d. | n.d. | n.d. | n.d | 1 |
| **NGF** | 1 (Earl et al. 2006) | n.d | n.s. | n.d.. | n.d. | n.d. | n.d. | n.d. | n.d. | 1 |
| **NP** | | 1 | 1 | n.d.. | n.d. | n.d. | n.d. | 2 | n.d. | 2 |
| **$PLA_2$** | 3-8 (Armugam et al. 2004; Pierre et al. 2005; Skejić et al. 2015) | 7 | n.s. | 5 | 5 | n.s. | 3 | 7 | 1-6 | 3 |
| **Snaclec** | | n.d. | n.s. | n.d.. | 1 | n.d. | 5 | 4 | 0-2 | 13 |

| | | | | | | | | | | |
|---|---|---|---|---|---|---|---|---|---|---|
| **SVMP** | | n.d. | n.s. | n.d.. | n.d. | n.d. | 1 | 1 | 1 | 2 |
| **Veficolin** | | n.d | n.d. | n.d. | n.d. | n.d. | n.d. | 1 | n.d | Partials |
| **Vespryn** | | n.d. | n.s. | n.d.. | n.d. | n.d. | n.d. | n.d. | n.d. | 1 |
| **Waprin** | | n.d. | n.s. | n.d. | n.d. | n.d. | n.d. | n.s. | n.d | 1 |
| TOTAL | | **25** | | **11** | **24** | | **27** | **48** | **12-25** | **53** |

*based identities on reads, complete transcripts not assembled; n.d. = not detected; n.s. = not specified, detected but isoform number not specified; 3FTx = three-finger toxin; 5'-NUC = 5'-nucleotidases; AChE = acetylcholinesterase; CRISP = cysteine-rich secretory protein; CVF = cobra venom factor; HYAL = hyaluronidase; KUN = Kunitz serine proteinase inhibitor; LAAO = L-amino acid oxidase; NGF = nerve growth factor; NP = natriuretic peptide; $PLA_2$ = phospholipase $A_2$; Snaclec = snake venom C-type lectin; SVMP = snake venom metalloproteinase.

**Supplemental Table 4. *De novo* assembled venom gland transcriptome for *Crotalus viridis*.**

| **Total reads** | **Assembly quality reads (phred > 30)** | **Genome alignment %** | **GG-Trinity contig # & average length** | ***De novo* Trinity contig # & average length** | **Extender contig # & average length** | **Nonredundant contig # & average length*** | **Expressed contigs ($\geq$1 TPM)** | **Expressed contigs average length** |
|---|---|---|---|---|---|---|---|---|
| 44,703,539 | 19,097,986 | 57.62% | 105,222<br>626 bp | 99,687<br>746 bp | 508<br>1,521 bp | 151,032<br>769 bp | 72,658 | 752 bp |

*Minimum of 150 bp

**Supplemental Table 5.** ***Crotalus viridis*** **toxin transcripts.**

| *Crotalus viridis* genome annotated transcript | ID used in this study | Transcripts per Million (TPM) at 96 hours post venom milking (hpvm) | Percentage of total toxin transcripts |
|---|---|---|---|
| crovir-transcript-17319 | BPP&NP | 1,874 | 0.2% |
| crovir-transcript-PLA2_A1 | PLA2_A1 | 33,873 | 5% |
| crovir-transcript-PLA2_B1 | PLA2_B1 | 16,788 | 2.2% |
| crovir-transcript-PLA2_C1 | PLA2_C1 | 10 | 0% |
| crovir-transcript-PLA2_gIIE | PLA2_gIIE | 125 | 0% |
| crovir-transcript-PLA2_K | PLA2_K | 202 | 0% |
| crovir-transcript-SVMP_1 | SVMP_1 | 6,980 | 1% |
| crovir-transcript-SVMP_2 | SVMP_2 | 465 | 0% |
| crovir-transcript-SVMP_3 | SVMP_3 | 182 | 0% |
| crovir-transcript-SVMP_4 | SVMP_4 | 66 | 0% |
| crovir-transcript-SVMP_5 | SVMP_5 | 506 | 0% |
| crovir-transcript-SVMP_6 | SVMP_6 | 9,765 | 1.3% |
| crovir-transcript-SVMP_7 | SVMP_7 | 8,102 | 1% |
| crovir-transcript-SVMP_8 | SVMP_8 | 5,419 | 1% |
| crovir-transcript-SVMP_9 | SVMP_9 | 5,290 | 1% |
| crovir-transcript-SVMP_10 | SVMP_10 | 6,594 | 1% |
| crovir-transcript-SVMP_11 | SVMP_11 | 0 | 0% |
| crovir-transcript-SVSP_1 | SVSP_1 | 9,393 | 1% |
| crovir-transcript-SVSP_2 | SVSP_2 | 208 | 0% |
| crovir-transcript-SVSP_3 | SVSP_3 | 1,470 | 0.1% |
| crovir-transcript-SVSP_4 | SVSP_4 | 2,024 | 0.2% |
| crovir-transcript-SVSP_5 | SVSP_5 | 4,2782 | 5.7% |
| crovir-transcript-SVSP_6 | SVSP_6 | 63 | 0% |
| crovir-transcript-SVSP_7 | SVSP_7 | 26,920 | 3.6% |
| crovir-transcript-SVSP_8 | SVSP_8 | 2,974 | 0.4% |

| | | | |
|---|---|---|---|
| crovir-transcript-SVSP_9 | SVSP_9 | 1,491 | 0.1% |
| Myotoxin_1 *de novo* assembly | Myotoxin_1 | 520,544 | 69.6% |
| Myotoxin_2 *de novo* assembly | Myotoxin_2 | 42,903 | 5.7% |

**Supplemental Table 6. *Crotalus tigris* toxin transcripts.**

| *Crotalus tigris* genome annotated transcript | Locus | ID used in this study | Transcripts per Million (TPM) at 96 hours post venom milking (hpvm) | Percentage of total toxin transcripts |
|---|---|---|---|---|
| XM_039327382.1 | n.s. | BPP&NP | 29,905 | 11.7% |
| XM_039329581.1 | LOC120301862 | CRISP_1 | 3,675 | 1.4% |
| XM_039361013.1 | LOC120315945 | LAAO_1 | 42 | 0% |
| XM_039367474.1 | LOC120319095 | PLA2_acidic | 84,191 | 33.1% |
| XM_039367475.1 | LOC120319096 | PLA2_basic | 57,284 | 22.5% |
| XM_039318646.1 | LOC120296723 | SNAC_1 | 1,223 | 0.4% |
| XM_039354202.1 | LOC120312896 | SVMP_1 | 8,457 | 3.3% |
| XM_039354199.1 | LOC120312894 | SVMP_2 | 1,670 | 0.6% |
| XM_039325703.1 | LOC120300115 | SVSP_1 | 1 | 0% |
| XM_039360544.1 | LOC120315719 | SVSP_2 | 49 | 0% |
| XM_039325705.1 | LOC120300115 | SVSP_3 | 46 | 0% |
| XM_039325727.1 | LOC120300123 | SVSP_4 | 32,407 | 12.7% |
| XM_039325704.1 | LOC120300115 | SVSP_5 | 9,808 | 3.8% |
| XM_039325717.1 | LOC120300120 | SVSP_6 | 4,603 | 1.8% |
| XM_039325725.1 | LOC120300122 | SVSP_7 | 2,555 | 1% |
| XM_039325712.1 | LOC120300119 | SVSP_8 | 1,633 | 0.6% |
| XM_039360553.1 | LOC120315719 | SVSP_9 | 887 | 0.3% |

| | | | | |
|---|---|---|---|---|
| XM_039360536.1 | LOC120315719 | SVSP_10 | 1,798 | 0.7% |
| XM_039325550.1 | LOC120300052 | SVSP_11 | 0 | 0% |
| XM_039325700.1 | LOC120300114 | SVSP_12 | 185 | 0% |
| XM_039325689.1 | LOC120300113 | SVSP_13 | 210 | 0% |
| XM_039325720.1 | LOC120300120 | SVSP_14 | 39 | 0% |
| XM_039325743.1 | LOC120300128 | SVSP_15 | 12 | 0% |
| XM_039325696.1 | LOC120300114 | SVSP_16 | 10 | 0% |
| XM_039325721.1 | LOC120300120 | SVSP_17 | 15 | 0% |
| XM_039325690.1 | LOC120300113 | SVSP_18 | 487 | 0.2% |
| XM_039325719.1 | LOC120300120 | SVSP_19 | 11 | 0% |
| XM_039325693.1 | LOC120300114 | SVSP_20 | 0 | 0% |
| XM_039325694.1 | LOC120300114 | SVSP_21 | 0 | 0% |
| XM_039325695.1 | LOC120300114 | SVSP_22 | 0 | 0% |
| XM_039325697.1 | LOC120300114 | SVSP_23 | 0 | 0% |
| XM_039325716.1 | L OC120300120 | SVSP_24 | 2 | 0% |
| XM_039325710.1 | LOC120300118 | SVSP_25 | 0 | 0% |
| XM_039325714.1 | LOC120300119 | SVSP_26 | 0 | 0% |
| XM_039325715.1 | LOC120300120 | SVSP_27 | 0 | 0% |
| XM_039325734.1 | LOC120300124 | SVSP_28 | 1 | 0% |
| XM_039325735.1 | LOC120300124 | SVSP_29 | 0 | 0% |
| XM_039325742.1 | LOC120300128 | SVSP_30 | 1 | 0% |
| XM_039325745.1 | LOC120300129 | SVSP_31 | 0 | 0% |
| XM_039325699.1 | LOC120300114 | SVSP_32 | 0 | 0% |
| XM_039325691.1 | LOC120300113 | SVSP_33 | 4 | 0% |
| XM_039325744.1 | LOC120300128 | SVSP_34 | 0 | 0% |
| XM_039325713.1 | LOC120300119 | SVSP_35 | 0 | 0% |
| XM_039325718.1 | LOC120300120 | SVSP_36 | 6 | 0% |
| XM_039325698.1 | LOC120300114 | SVSP_37 | 14 | 0% |

| | | | | |
|---|---|---|---|---|
| XM_039325701.1 | LOC120300114 | SVSP_38 | 9 | 0% |
| XM_039362118.1 | LOC120316470 | VEGF_1 | 13,081 | 5.1% |

n.s. = not specified

**Supplemental Table 7. Gene expression data for viperids from milked and unmilked venom glands. [EXCEL]**

**Supplemental Table 8. Gene Set Enrichment Analysis for milked and unmilked elapid and viperid venom glands. [EXCEL]**

**Supplemental Table 9. Epigenetic modifiers, transcription factors and co-factors upregulated over 40-fold after venom milking of *Pseudonaja textilis* venom glands.**

| Gene | Species | Expression | Activity | Function | UniProtKB entry name | Fold-change* | hpvm |
|---|---|---|---|---|---|---|---|
| **Epigenetic Modifiers** | | | | | | | |
| *SRCAP* | *P. textilis* | Upregulated variant X2 [Downregulated variant X1] | Exchanges nucleosome histones to render DNA accessible | Activation | Q6ZRS2 | 86-fold [0.004-fold] | 96 |
| *KMT2A* | *P. textilis* | Upregulated variant X7 | Trimethylates Lys-4 of H3, and acetylates Lys-16 of H4 | Activation | Q03164 | 80-fold | 96 |
| *KMT2D* | *P. textilis* | Upregulated variant X4 [Downregulated variant X6] | Monomethylates Lys-4 of H3 | Activation (super-enhancer) | O14686 | 64-fold [0.006-fold] | 96 |
| *CHD8* | *P. textilis* | Upregulated variant X4 | Interacts with H3 di- and trimethylated at Lys-4, recruiting H1 | Repression | Q9HCK8 | 52-fold | 96 |

| | | | | | | | |
|---|---|---|---|---|---|---|---|
| *KMT2C* | *P. textilis* | Upregulated variant X9 | Methylates Lys-4 of H3 | Activation (super-enhancer) | Q8NEZ4 | 45-fold | 96 |
| *JMJD1C* | *P. textilis* | Upregulated variant X4 | Demethylates Lys-9 of H3 | Activation | Q15652 | 43-fold | 96 |
| *PCGF5* | *P. textilis* | Downregulated variant X3 | Component of a Polycomb group repressor complex | Repression | Q86SE9 | 0.032-fold | 96 |
| *SETDB1* | *P. textilis* | Downregulated variant X1 | Trimethylates Lys-9 of H3 | Repression | Q15047 | 0.021-fold | 96 |
| *PRDM2* | *P. textilis* | Downregulated variant X4 | Methylates Lys-9 of H3 | Activation | Q13029 | 0.010-fold | 96 |
| **Transcription Factors and Co-factors** | | | | | | | |
| *SP1* | *P. textilis* | Upregulated variant X2 [Downregulated variant X3] | Transcription factor that binds GC-rich motifs | Activation or Repression | P08047 | 79-fold [0.025-fold] | 96 |
| *FOXN2* | *P. textilis* | Upregulated variant X2 | Transcription factor that binds purine-rich regions | Activation | P32314 | 41-fold | 96 |
| *LCOR* | *P. textilis* | Upregulated variant X1 | Transcriptional corepressor | Repression | Q96JN0 | 40-fold | 96 |
| *LRRFIP1* | *P. textilis* | Downregulated variant X16 | Transcriptional repressor that binds a GC-rich consensus sequence | Repression | Q13045 | 0.039-fold | 96 |
| *ZNF665* | *P. textilis* | Downregulated variant X2 | Unknown | Unknown | Q9H7R5 | 0.034-fold | 96 |
| *EBF1* | *P. textilis* | Downregulated variant X1 | Transcription factor that binds variations of the palindromic sequence 5’-ATTCCCNNGGGAATT-3’ | Activation | Q9UH73 | 0.030-fold | 96 |
| *GTF3C1* | *P. textilis* | Downregulated variant X1 | Required for the transcription of rRNA | Activation | Q12789 | 0.025-fold | 96 |

| | | | | | | | |
|---|---|---|---|---|---|---|---|
| *ATF2* | *P. textilis* | Downregulated variant X2 | Transcription factor that binds to cAMP response element or AP-1 consensus sequences | Activation | P15336 | 0.011-fold | 96 |
| *MLF2* | *P. textilis* | Downregulated variant X2 | Regulates transcription | Unknown | Q15773 | 0.010-fold | 96 |

*Determined from GFOLD values; hpvm = hours post venom milking

**Supplemental Table 10. Epigenetic modifiers, transcription factors and co-factors upregulated over 40-fold, downregulated less than 0.04-fold after venom milking from viperid venom glands.**

| Gene | Species | Condition | Protein Activity | Function | UniProtKB entry name | Fold-change* | hpvm |
|---|---|---|---|---|---|---|---|
| **Epigenetic Modifiers** | | | | | | | |
| *JADE2* | *C. tigris* | Downregulated variant X3 | Scaffold for HBO1 complexes with acetyltransferase activity | Activation | Q9NQC1 | 0.032-fold | 96 |
| *BICRAL* | *C. tigris* | Downregulated variant X1 | Component of the SWI/SNF chromatin remodeling subcomplex | Activation | Q6AI39 | 0.032-fold | 96 |
| *ATF7* | *C. tigris* | Downregulated variant X2 | Recruits histone methyltransferases | Activation or Repression | P17544 | 0.029-fold | 24 |
| *CREBBP* | *C. tigris* | Downregulated variant X1 | Acetylates histones, transcriptional coactivator | Activation | Q92793 | 0.026-fold<br>0.020-fold | 24<br>96 |
| *CBX7* | *C. tigris* | Downregulated variant X1 | Component of a Polycomb group repressor complex | Repression | O95931 | 0.024-fold | 24 |
| *WIZ* | *C. tigris* | Downregulated variant X4 | Recruits histone methyltransferases | Repression | O95785 | 0.019-fold | 24 |

| | | | | | | | |
|---|---|---|---|---|---|---|---|
| *KDM5C* | *C. tigris* | Downregulated variant X2 | Demethylates Lys-4 of H3 | Repression | P41229 | 0.018-fold | 24 |
| *EZH1* | *C. tigris* | Downregulated variant X7 | Component of a Polycomb group repressor complex, methylates Lys-27 of H3 | Repression | Q92800 | 0.017-fold | 24 |
| *CBX5* | *C. tigris* | Downregulated variant X2 | Repressor that binds H3 tails methylated at Lys-9 | Repression | P45973 | 0.016-fold | 24 |
| *ASH1L* | *C. tigris* | Downregulated variant X2 | Trimethylates Lys-36 of H3 | Activation or Repression | Q9NR48 | 0.016-fold | 24 |
| *HDAC7* | *C. tigris* | Downregulated variant X7 | Deacetylates core histones H2A, H2B, H3 and H4 | Repression | Q8WUI4 | 0.013-fold | 24 |
| *MLLT10* | *C. tigris* | Downregulated variant X1 | Dimethylation of Lys-79 of H3 | Activation | P55197 | 0.011-fold | 24 |
| *BPTF* | *C. tigris* | Downregulated variant X1 | Component of the NURF-ISWI chromatin-remodeling complex | Activation or Repression | Q12830 | 0.010-fold | 96 |
| **Transcription Factors and Co-factors** | | | | | | | |
| *NFIB* | *C. tigris* | Upregulated variant X12 [Downregulated variant X3] | Transcription factor | Activation | O00712 | 104-fold [0.026-fold] 133-fold | 24<br><br>96 |
| *NFKBIE* | *C. viridis* | Downregulated | Inhibits NFκB | Repression | O00221 | 0.034-fold | 96 |
| *NCOA1* | *C. tigris* | Downregulated variant X3 | Coactivator that stimulates transcription in a hormone-dependent fashion | Activation | Q15788 | 0.039-fold | 24 |
| *ZBTB14* | *C. tigris* | Downregulated variant X1 | Transcriptional activator that binds at the consensus sequence 5'-CCTGCACAGTTCACGGA-3' | Activation or Repression | Q9ULJ3 | 0.039-fold | 24 |

| | | | | | | | |
|---|---|---|---|---|---|---|---|
| *TRERF1* | *C. tigris* | Downregulated variant X15 | Regulates transcription | Activation | Q96PN7 | 0.038-fold | 96 |
| *CUX1* | *C. tigris* | Downregulated variant X3 | Prevents binding of positively-activing CCAAT factors to promoters | Repression | P39880 | 0.037-fold | 24 |
| *ZNF777* | *C. tigris* | Downregulated variant X3 | Unknown | Unknown | Q9ULD5 | 0.034-fold | 24 |
| *NFKB2* | *C. tigris* | Downregulated variant X1 and X4 | Regulates transcription | Activation or Repression | Q00653 | X1: 0.034-fold;<br>X4: 0.030-fold | 24 |
| | | | | | | X1: 0.025-fold;<br>X4: 0.022-fold | 96 |
| *TCF4* | *C. tigris* | Downregulated variant X11 | Transcriptional activator by binding to the E box | Activation | P15884 | 0.033-fold<br>0.025-fold | 24<br>96 |
| *NFAT5* | *C. tigris* | Downregulated variant X8 | Transcription factor | Activation | O94916 | 0.031-fold | 24 |
| *ZNF507* | *C. tigris* | Downregulated variant X3 | Unknown | Unknown | Q8TCN5 | 0.031-fold | 96 |
| *ZBF146* | *C. tigris* | Downregulated variant X2 | Regulates transcription | Activation | P28347 | 0.030-fold | 96 |
| *TEAD1* | *C. tigris* | Downregulated variant X8 | Transcription factor | Activation | P28347 | 0.030-fold | 96 |
| *ZKSCAN7* | *C. tigris* | Downregulated variant X7 | Unknown | Unknown | Q9P0L1 | 0.029-fold | 96 |
| *ZNF25* | *C. tigris* | Downregulated variant X1 | Regulates transcription | Unknown | P17030 | 0.029-fold<br>0.012-fold | 24<br>96 |
| *ZNF148* | *C. tigris* | Downregulated variant X2 | Represses transcription by binding to the G-rich box in the enhancer region of genes | Repression | Q9UQR1 | 0.027-fold | 24 |
| *GLI3* | *C. tigris* | Downregulated variant X9 | Regulates transcription | Activation or Repression | P10071 | 0.024-fold | 24 |

| | | | | | | | |
|---|---|---|---|---|---|---|---|
| *ETV5* | *C. tigris* | Downregulated variant X1 | Regulates transcription | Unknown | P41161 | 0.023-fold | 24 |
| *NCOA5* | *C. tigris* | Downregulated variant X3 | Regulates transcription | Activation or Repression | Q9HCD5 | 0.022-fold | 24 |
| *SOX6* | *C. tigris* | Downregulated variant X15 | Regulates transcription, can bind to enhancer motifs | Activation or Repression | P35712 | 0.021-fold | 24 |
| *NFIC* | *C. tigris* | Downregulated variant X2 | Transcription factor | Activation | P08651 | 0.020-fold | 24 |
| *ZSCAN20* | *C. tigris* | Downregulated variant X2 | Unknown | Unknown | P17040 | 0.020-fold | 96 |
| *ZNF292* | *C. tigris* | Downregulated variant X2 | Transcription factor | Unknown | O60281 | 0.019-fold<br>0.014-fold | 24<br>96 |
| *MED24* | *C. tigris* | Downregulated variant X1 | Component of the Mediator complex for transcription | Activation | O75448 | 0.019-fold | 24 |
| *ARID4B* | *C. tigris* | Downregulated variant X1 | Recruits regulatory complexes | Activation or Repression | Q4LE39 | 0.019-fold | 24 |
| *MED12L* | *C. tigris* | Downregulated variant X2 | Component of the Mediator complex for transcription | Activation | Q86YW9 | 0.018-fold | 24 |
| *FOXO3* | *C. tigris* | Downregulated variant X2 | Transcription factor that binds 5’-[AG]TAAA[TC]A-3’ | Activation | O43524 | 0.018-fold | 24 |
| *MED23* | *C. tigris* | Downregulated variant X2 | Component of the Mediator complex for transcription | Activation | Q9ULK4 | 0.017-fold | 24 |
| *NR1D1* | *C. tigris* | Downregulated variant X2 | Regulates transcription of genes involved in circadian rhythm and metabolic pathways | Repression | P20393 | 0.016-fold | 24 |
| *HIVEP2* | *C. tigris* | Downregulated variant X3 | Bind to enhancer elements to regulate transcription | Activation | P31629 | 0.015-fold<br>0.011-fold | 24<br>96 |
| *DBP* | *C. tigris* | Downregulated variant X2 | Transcription factor that binds 5’-RTTAYGTAAY-3’ | Activation | Q10586 | 0.013-fold | 24 |

| | | | | | | | |
|---|---|---|---|---|---|---|---|
| *ZNF384* | *C. tigris* | Downregulated variant X1 | Transcription factor that binds the consensus sequence [GC]AAAAA | Activation | Q8TF68 | 0.012-fold | 96 |
| *GTF2IRD2* | *C. tigris* | Downregulated variant X20 | Regulates transcription | Activation | Q86UP8 | 0.009-fold | 24 |
| *FOXP4* | *C. tigris* | Downregulated variant X9 | Regulates transcription | Repression | Q8IVH2 | 0.009-fold<br>0.007-fold | 24<br>96 |
| *ZNF652* | *C. tigris* | Downregulated variant X2 | Regulates transcription | Repression | Q9Y2D9 | 0.009-fold | 24 |
| *NCOA2* | *C. tigris* | Downregulated variant X6 | Transcriptional coactivator | Activation | Q15596 | 0.008-fold | 24 |
| *PCGF2* | *C. tigris* | Downregulated variant X1 | Transcriptional repressor that binds 5’-GACTNGACT-3’ | Repression | P35227 | 0.005-fold | 24 |

*Determined from GFOLD values; hpvm = hours post venom milking

**Supplemental Table 11. miRNAs from the *Pseudonaja textilis* milked and unmilked venom glands [EXCEL]**

**Supplemental Table 12. miRNAs from the *Crotalus viridis* milked venom glands [EXCEL]**

**Supplemental Table 13. Toxins transcripts that are targeted by the top ten miRNAs in each venom gland.**

| miRDeep2 program ID | Genome coordinate | Snake miRNA ortholog | Toxin target |
|---|---|---|---|
| ***Pseudonaja textilis* milked venom gland** | | | |
| NW_020769346.1_3091 | NW_020769346.1:11487596..11487656:+ | oha-miR-148a-3p | SNAC_5 |
| NW_020769345.1_3030 | NW_020769345.1:9005083..9005144:+ | oha-miR-26-5p | |
| NW_020769314.1_968 | NW_020769314.1:12875940..12875994:- | oha-miR-143-3p | |
| NW_020769360.1_3679 | NW_020769321.1:22243475..22243537:+ | oha-miR-181c-5p | PLA2_1, 2, 3 |
| NW_020777308.1_6410 | NW_020777308.1:336..396:+ | oha-miR-199a-5p | SNAC_8, vFX |
| NW_020769331.1_2223 | NW_020769331.1:10873124..10873186:+ | oha-miR-27b-3p | |
| NW_020769464.1_5375 | NW_020769464.1:684555..684617:+ | oha-miR-22a | |
| NW_020769315.1_1059 | NW_020769315.1:16026068..16026129:- | oha-miR-10c-5p | SNAC_2, 3, 7, 9, and 12 |
| NW_020769354.1_3443 | NW_020769354.1:3064746..3064807:- | oha-miR-21-5p | |
| NW_020769311.1_516 | NW_020769311.1:29915245..29915323:+ | oha-let-7c-5p | |
| ***Pseudonaja textilis* unmilked venom gland** | | | |
| NW_020769315.1_6937 | NW_020769315.1:16026068..16026129:- | oha-miR-10c-5p | SNAC_2, 3, 7, 9, and 12 |
| NW_020769314.1_6160 | NW_020769314.1:12875939..12875994:- | oha-miR-143-3p | |
| NW_020777307.1_38858 | NW_020777308.1:336..396:+ | oha-miR-199a-5p | SNAC_8, vFX |
| NW_020769346.1_19401 | NW_020769346.1:11487596..11487656:+ | oha-miR-148a-3p | SNAC_5 |
| NW_020769360.1_23039 | NW_020769321.1:22243475..22243537:+ | oha-miR-181c-5p | PLA2_1, 2, and 3 |
| NW_020769345.1_19102 | NW_020769345.1:9005083..9005144:+ | oha-miR-26-5p | |

| | | | |
|---|---|---|---|
| NW_020769345.1_19115 | NW_020769345.1:9521124..9521181:+ | oha-miR-375-3p | |
| NW_020769311.1_3516 | NW_020769311.1:29915245..29915323:+ | oha-let-7c-5p | |
| NW_020769408.1_30640 | NW_020769408.1:2603663..2603725:+ | oha-miR-191-5p | |
| NW_020769331.1_14662 | NW_020769331.1:1863827..1863885:- | oha-miR-101b-3p | 3FTx_7, 8, 10, 11, 12, and 13, vFV |
| ***Crotalus viridis*** **milked venom gland** | | | |
| CM012306.1_1866 | CM012306.1:271893842..271893901:+ | pbv-miR-21-5p | |
| CM012323.1_17538 | CM012323.1:35887469..35887529:- | pbv-miR-148a-3p | |
| CM012319.1_15599 | CM012319.1:559445..559506:+ | pbv-miR-200a-3p | SVMP_9 |
| CM012307.1_4766 | CM012307.1:128522563..128522624:+ | pbv-miR-26-5p | SVMP_9 |
| CM012306.1_736 | CM012306.1:103174546..103174607:+ | pbv-miR-26-5p | SVMP_9 |
| CM012323.1_17395 | CM012323.1:2374795..2374856:- | pbv-miR-26-5p | SVMP_9 |
| CM012309.1_9993 | CM012309.1:67134688..67134748:- | pbv-miR-100-5p | |
| CM012306.1_1850 | CM012306.1:270639415..270639475:+ | pbv-miR-215-5p | SVMP_4 |
| CM012307.1_6610 | CM012307.1:174840972..174841030:- | pbv-miR-101-3p | |
| CM012308.1_7771 | CM012308.1:122669715..122669774:+ | pbv-miR-101-3p | |

**Supplemental Table 14. miRNA target predictions for *Pseudonaja textilis* milked and unmilked venom glands [EXCEL]**

**Supplemental Table 15. miRNA target predictions for *Crotalus viridis* venom glands [EXCEL]**

**Supplemental Table 16. Transcription factors regulating venom genes.**

| Species | Gene(s) | Transcription factors | Activity | Reference |
|---|---|---|---|---|
| **Elapidae** | | | | |
| *Bungarus multicinctus* | 3FTxs | CACCC-box<br>EFII<br>GR<br>LF-A1<br>NF-I<br>Pit-1<br>Sp1<br>TATA-box | Promoter binding sites | (Chang et al. 2002a; Chang et al. 2002b) |
| *Boiga dendrophilia* | 3FTx | NFI<br>TATA-box | Promoter binding sites | (Pawlak and Kini 2008) |
| *Naja atra* | 3FTx | CACCC-box<br>EFII<br>NFI<br>Sp1<br>TATA-box | Promoter binding sites | (Chang et al. 2004) |
| *Naja sputatrix* | 3FTx | **AP-1**<br>AP-2<br>**CACCC-box**<br>**C/EBP**<br>CP-1<br>c-Myc<br>E2BP<br>E47<br>EFII<br>GATA-1<br>**GR** | Promoter binding sites (Bold have been experimentally shown to regulate expression) | (Lachumanan et al. 1998; Afifiyan et al. 1999; Ma et al. 2001; Ma et al. 2002) |

| | | | | |
|---|---|---|---|---|
| | | Ikaros<br>NF$\kappa$B<br>NF-E<br>NFI<br>NF-IL6<br>PEA3<br>Pit-1<br>PuF<br>**Sp1**<br>**TATA-box**<br>TCF-1<br>UBP-1 | | |
| *Pseudonaja textilis* | 3FTx (long-chain like) | AP-2<br>GATA-2<br>GC-box (Sp1)<br>TATA-box | Expression | (Gong et al. 2001) |
| *Pseudonaja textilis* | 3FTx (short-chain) | AP-1<br>CCAAT-box (C/EBP)<br>C/EBP<br>GATA-2<br>TATA-box | Expression | (Gong et al. 2000) |
| *Tropidechis carinatus*<br>*Pseudonaja textilis* | FX | GATA-4<br>HNF-4<br>CCAAT-box (C/EBP)<br>Sp1/Sp3<br>TATA-box<br>Y-box | Expression | (Reza et al. 2007) |
| *Tropidechis carinatus* | FX | HMGB2<br>Sp3<br>YY1 | Silencing | (Han et al. 2016) |
| *Bungarus multicinctus* | KUN | BR-C<br>CD28RC | | (Wu and Chang 2000) |

| | | | | |
|---|---|---|---|---|
| | | CF2-11<br>Croc<br>Elf-1<br>Elk-1<br>GATA-1<br>GC1<br>HFH<br>MEF-2<br>Myc-CF1<br>NF-1<br>NF-1L-2A<br>NFAT-1<br>NFH3-1<br>NF$\kappa$B<br>Pit-1a<br>Pu-box<br>Sox-5<br>SRF<br>TATA-box | | |
| *Bungarus multicinctus* | $PLA_2$ Group I | AP-1<br>AP-2<br>Bcd<br>C/EBP<br>CS<br>G6-factor<br>GAGA-factor<br>LyF-1<br>NF-IL6<br>CF1<br>$\gamma$-IRE<br>cMyb<br>MyoD<br>Sn | Promoter binding sites | (Wu and Chang 2000; Chu and Chang 2002) |

| | | | | |
|---|---|---|---|---|
| | | Sox-5<br>Sp1<br>STE12<br>TATA-box<br>TG repeats | | |
| *Laticauda semifasciata* | $PLA_2$<br>Group I | CCAAT-box (C/EBP)<br>E-box<br>GC-box (Sp1)<br>TATA-box | Promoter binding sites | (Fujimi et al. 2004) |
| *Naja sputatrix* | $PLA_2$<br>Group I | AP-1<br>**AP-2**<br>C/EBP<br>CF1<br>CS<br>cMyb<br>**γ-IRE**<br>MyoD<br>NF-IL6<br>**Sp1**<br>TATA-box<br>**TG repeats** | Promoter binding sites<br>(Bold have been experimentally shown to regulate expression) | (Jeyaseelan et al. 2000) |
| *Protobothrops flavoviridis* | $PLA_2$<br>Group I | ESE-3 | Expression | (Nakamura et al. 2014) |
| *Pseudonaja textilis* | $PLA_2$<br>Group IB | C/EBP<br>GATA-3<br>GC-box (Sp1)<br>TATA-box | Expression | (Armugam et al. 2004) |
| **Viperidae** | | | | |
| *Crotalus viridis* | $PLA_2$<br>SVMP<br>SVSP | GRHL1<br>NFI | Promoter binding sites and | (Schield et al. 2019) |

| | | | | |
|---|---|---|---|---|
| | | | upregulation after venom milking | |
| *Crotalus tigris* | $PLA_2$ Group II | NFIC | Accessible promoter binding sites | (Margres et al. 2021) |
| *Crotalus tigris* | SVMP | FOXA1<br>FOXA2 | Accessible promoter binding sites | (Margres et al. 2021) |
| *Crotalus tigris* | SVSP | GRHL2<br>NFI | Accessible promoter binding sites | (Margres et al. 2021) |
| *Echis coloratus* | SVMP | BARBIE<br>ETS<br>ETS1<br>ETS2<br>KAISO<br>NKX2.5<br>NRF2<br>TEF1<br>ZTA | Promoter binding sites | (Hargreaves et al. 2014) |
| *Echis coloratus* | SVSP | CAAT<br>CRX<br>DTYPEPA<br>FOXD3<br>HFH4<br>HNF3B<br>HNF4<br>LPOLYA<br>MZF1<br>NFY<br>PBX<br>SPZ1 | Promoter binding sites | (Hargreaves et al. 2014) |

| | | | | |
|---|---|---|---|---|
| | | STAT5<br>TCF1 | | |
| *Echis coloratus* | $PLA_2$<br>Group II | E12<br>E2A<br>ETF<br>LFA1<br>MYOD<br>Myogenin<br>PAX6<br>TAL1<br>Tbx | Promoter binding sites | (Hargreaves et al. 2014) |
| *Protobothrops flavoviridis* | $PLA_2$<br>Group II | AP-1<br>AP-2<br>HNF3<br>NF*κ*B<br>Sp1<br>Tbx3 | Promoter binding sites | (Hargreaves et al. 2014) |
| *Crotalus durissus terrificus* | Crotamine | NF-1<br>NF*κ*B<br>Sp1<br>TATA-box | Promoter binding sites | (Rádis-Baptista et al. 2003) |

*Pseudonaja textilis* venom genes are highlighted in grey.

**Supplemental Table 17. Comparison of Elapidae and Viperidae characteristics.**

| | *Pseudonaja textilis* / Elapidae | *Crotalus* spp. / Viperidae | References |
|---|---|---|---|
| **Fangs and skull** | • Proteroglyphous<br>• Fixed fang at the front of the maxilla<br>• Less mobile maxillary bone<br>• Short fangs | • Solenoglyphous<br>• Moveable fang on the shortened maxilla<br>• Highly kinetic maxillary bone<br>• Long fangs | (Kardong 1982; Kochva 1987) |

| | | | |
|---|---|---|---|
| **Venom gland and accessory gland** | • Narrow venom gland lumen<br>• Smaller volume of venom<br>• Venom secretion mainly stored inside cells<br>• Accessory gland is the distal part of the venom gland and only one duct is present | • Wide venom gland lumen<br>• Larger volumes of venom<br>• Fewer secretion granules stored in cells<br>• Accessory gland is separate from the venom gland and connected to a secondary duct | (Kochva 1987; Kerkkamp et al. 2015) |
| **Muscle that compresses venom gland, 'compressor glandulae'** | Adductor externus superficialis | Adductor externus profundus | (Jackson 2003) |
| **Venom gland secretory cells after milking** | Minimal change to secretory cells | Secretory cells become elongated, increasing from cuboidal to columnar with proliferation of the rough ER | (Oron and Bdolah 1973; Kochva et al. 1982; Mackessy 1991; Lachumanan et al. 1999) |
| **Biological processes upregulated after milking** | Chromatin and histone remodeling, regulation of transcription and phospholipid biosynthesis (including phosphatidylinositol-3-phosphate signaling and inositol phosphate metabolic process) | Transcription, protein translation and transport, and the UPR<br>(negative regulation of translation) | (Perry et al. 2020; Perry et al. 2022) for viperid<br><br>This paper |
| **Biological processes downregulated after milking** | Proteins involved in striated muscle contraction and sarcomere assembly | Complement activation, immune response, cellular component organization and metabolic processes, negative mechanisms of regulating nucleobase-containing macromolecules, including transcriptional repressors | This paper |
| **Upregulation of chromatin remodeling factors after milking** | Snf2 related CREB activator protein (*SRCAP*, 86-fold), jumonji domain containing 1C (*JMJD1C*, 43-fold), lysine methyltransferases 2A (*KMT2A*, 80-fold), KMT2C (*KMT2C*, 45-fold), KMT2D | Chromatin modifiers were not seen as highly upregulated | This paper |

| | | | |
|---|---|---|---|
| | (*KMT2D*, 64-fold), and chromodomain-helicase-DNA-binding protein 8-like (*CHD8*, 52-fold) | | |
| **Upregulation of transcription factors after milking** | *SP1*, 79-fold<br>*FOXN2* and *LCOR* 41- and 40-fold<br>*NFIA* 17-fold, *NFIB* 5-fold, and *NFIX* 17-fold at 96 hpvm | *SP1*, 4- to 5-fold<br>*CREB3L3* 11-fold at 96 hpvm. *NFIA* 31-fold at 96 hpvm, *NFIB* 133-fold, and *NFIX* 24-fold 96 hpvm in *C. tigris; NFIA* and *NFIB* 5-fold and 6-fold, at 96 hpvm in *C. viridis* | This paper |
| ***Cis*-Regulatory Elements present in toxin promoters that are upregulated over 10-fold** | For three-finger toxins (3FTx): Sp1/CACCC-box, NFI, and interferon regulatory factor (IRF)<br>For phospholipase $A_2$ ($PLA_2$) toxins: Sp1/CACCC-box and NFI | For $PLA_2$ toxins:<br>NFI, retinoic acid receptor (RAR), upstream stimulatory factor 1 (USF1), and thyroid hormone (3,5,3'-triiodothyronine) receptor (T3R) | This paper |
| **Toxins** | Dominated by 3FTxs and group I $PLA_2$s | Primarily snake venom metalloproteinases (SVMPs), serine proteinases (SVSPs), and group II $PLA_2$s | (Mackessy 2010; Tasoulis and Isbister 2017) |
| **Toxin gene locations** | Macrochromosomes | Microchromosomes | (Shibata et al. 2018; Suryamohan et al. 2020) |
| **Fold-change in toxin expression after milking** | 2-fold or less | Increases over 40-fold, one SVMP exhibited a 127-fold change at 96 hpvm in *C. viridis* | This paper |
| **Toxin transcripts targeted by microRNAs** | Snaclecs, group I $PLA_2$s, 3FTxs and pseutarin C | SVMPs | This paper |
| **Biological processes targeted by microRNAs in milked venom glands** | Intracellular transport, catabolic processes, metabolic processes, organelle organization, and ER to Golgi mediated transport | Regulation of mRNA processing, catabolic processes, negative regulation of cytoskeleton organization, steroid hormone receptor signal, stress granule assembly, transport, and ER stress | This paper |
| **MicroRNAs unique** | *miR-375* | *miR-215-5p* | This paper |

Abbreviations: hpvm = hours post venom milking

Afifiyan F, Armugam A, Tan CH, Gopalakrishnakone P, Jeyaseelan K. 1999. Postsynaptic alpha-neurotoxin gene of the spitting cobra, *Naja naja sputatrix*: structure, organization, and phylogenetic analysis. *Genome Res* **9**: 259-266.

Armugam A, Gong N, Li X, Siew PY, Chai SC, Nair R, Jeyaseelan K. 2004. Group IB phospholipase A2 from *Pseudonaja textilis*. *Arch Biochem Biophys* **421**: 10-20.

Chang LS, Chung C, Lin J, Hong E. 2002a. Organization and phylogenetic analysis of kappa-bungarotoxin genes from *Bungarus multicinctus* (Taiwan banded krait). *Genetica* **115**: 213-221.

Chang LS, Chung C, Wu BN, Yang CC. 2002b. Characterization and gene organization of Taiwan banded krait (*Bungarus multicinctus)* gamma-bungarotoxin. *J Protein Chem* **21**: 223-229.

Chang LS, Lin SK, Chung C. 2004. Molecular cloning and evolution of the genes encoding the precursors of taiwan cobra cardiotoxin and cardiotoxin-like basic protein. *Biochemical genetics* **42**: 429-440.

Chu Y-P, Chang L-S. 2002. The organization of the genes encoding the A chains of β-bungarotoxins: evidence for the skipping of exon. *Toxicon* **40**: 1437-1443.

Earl ST, Birrell GW, Wallis TP, St Pierre LD, Masci PP, de Jersey J, Gorman JJ, Lavin MF. 2006. Post-translational modification accounts for the presence of varied forms of nerve growth factor in Australian elapid snake venoms. *Proteomics* **6**: 6554-6565.

Filippovich I, Sorokina N, Masci PP, de Jersey J, Whitaker AN, Winzor DJ, Gaffney PJ, Lavin MF. 2002. A family of textilinin genes, two of which encode proteins with antihaemorrhagic properties. *Br J Haematol* **119**: 376-384.

Fujimi TJ, Yasuoka S, Ogura E, Tsuchiya T, Tamiya T. 2004. Comparative analysis of gene expression mechanisms between group IA and IB phospholipase A2 genes from sea snake *Laticauda semifasciata*. *Gene* **332**: 179-190.

Gong N, Armugam A, Jeyaseelan K. 2000. Molecular cloning, characterization and evolution of the gene encoding a new group of short-chain α-neurotoxins in an Australian elapid, *Pseudonaja textilis*. *FEBS Letters* **473**: 303-310.

Gong N, Armugam A, Mirtschin P, Jeyaseelan K. 2001. Cloning and characterization of the pseudonajatoxin b precursor. *Biochem J* **358**: 647-656.

Han SX, Kwong S, Ge R, Kolatkar PR, Woods AE, Blanchet G, Kini RM. 2016. Regulation of expression of venom toxins: silencing of prothrombin activator trocarin D by AG-rich motifs. *FASEB J* **30**: 2411-2425.

Hargreaves AD, Swain MT, Hegarty MJ, Logan DW, Mulley JF. 2014. Genomic and transcriptomic insights into the regulation of snake venom production. *bioRxiv* doi:10.1101/008474: 008474.

Jackson K. 2003. The evolution of venom-delivery systems in snakes. *Zool J Linn Soc* **137**: 337-354.

Jeyaseelan K, Armugam A, Donghui M, Tan N-H. 2000. Structure and phylogeny of the venom group I phospholipase A2 gene. *Mol Biol Evol* **17**: 1010-1021.

Kardong KV. 1982. The evolution of the venom apparatus in snakes from colubrids to viperids and elapids. *Mem Inst Butantan* **46**: 105-118.

Kerkkamp HMI, Casewell NR, Vonk FJ. 2015. Evolution of the Snake Venom Delivery System. In *Evolution of Venomous Animals and Their Toxins*, doi:10.1007/978-94-007-6727-0_11-1 (ed. P Gopalakrishnakone, A Malhotra), pp. 1-11. Springer Netherlands, Dordrecht.

Kochva E. 1987. The origin of snakes and evolution of the venom apparatus. *Toxicon* **25**: 65-106.

Kochva E, Tönsing L, Louw AI, Liebenberg NvdW, Visser L. 1982. Biosynthesis, secretion and *in vivo* isotopic labelling of venom of the Egyptian cobra, *Naja haje annulifera*. *Toxicon* **20**: 615-635.

Lachumanan R, Armugam A, Durairaj P, Gopalakrishnakone P, Tan CH, Jeyaseelan K. 1999. In situ hybridization and immunohistochemical analysis of the expression of cardiotoxin and neurotoxin genes in *Naja naja sputatrix*. *J Histochem Cytochem* **47**: 551-560.

Lachumanan R, Armugam A, Tan CH, Jeyaseelan K. 1998. Structure and organization of the cardiotoxin genes in *Naja naja sputatrix*. *FEBS Lett* **433**: 119-124.

Ma D, Armugam A, Jeyaseelan K. 2001. Expression of cardiotoxin-2 gene. Cloning, characterization and deletion analysis of the promoter. *Eur J Biochem* **268**: 1844-1850.

Ma D, Armugam A, Jeyaseelan K. 2002. Alpha-neurotoxin gene expression in *Naja sputatrix*: identification of a silencer element in the promoter region. *Arch Biochem Biophys* **404**: 98-105.

Mackessy SP. 1991. Morphology and ultrastructure of the venom glands of the northern pacific rattlesnake *Crotalus viridis oreganus*. *J Morphol* **208**: 109-128.

Mackessy SP. 2010. The field of reptile toxinology: snakes, lizards and their venoms. In *Handbook of Venoms and Toxins of Reptiles*, (ed. M SP), pp. 2-23. CRC Press/Taylor & Francis Group, Boca Raton, FL.

Margres MJ, Rautsaw RM, Strickland JL, Mason AJ, Schramer TD, Hofmann EP, Stiers E, Ellsworth SA, Nystrom GS, Hogan MP et al. 2021. The Tiger Rattlesnake genome reveals a complex genotype underlying a simple venom phenotype. *Proc Natl Acad Sci USA* **118**: e2014634118.

Nakamura H, Murakami T, Hattori S, Sakaki Y, Ohkuri T, Chijiwa T, Ohno M, Oda-Ueda N. 2014. Epithelium specific ETS transcription factor, ESE-3, of *Protobothrops flavoviridis* snake venom gland transactivates the promoters of venom phospholipase A2 isozyme genes. *Toxicon* **92**: 133-139.

Oron U, Bdolah A. 1973. Regulation of protein synthesis in the venom gland of viperid snakes. *J Cell Biol* **56**: 177-190.

Pawlak J, Kini RM. 2008. Unique gene organization of colubrid three-finger toxins: Complete cDNA and gene sequences of denmotoxin, a bird-specific toxin from colubrid snake *Boiga dendrophila* (Mangrove Catsnake). *Biochimie* **90**: 868-877.

Perry BW, Gopalan SS, Pasquesi GIM, Schield DR, Westfall AK, Smith CF, Koludarov I, Chippindale PT, Pellegrino MW, Chuong EB et al. 2022. Snake venom gene expression is coordinated by novel regulatory architecture and the integration of multiple co-opted vertebrate pathways. *GenomeRes* doi:10.1101/gr.276251.121.

Perry BW, Schield DR, Westfall AK, Mackessy SP, Castoe TA. 2020. Physiological demands and signaling associated with snake venom production and storage illustrated by transcriptional analyses of venom glands. *Sci Rep* **10**: 18083.

Pierre LS, Woods R, Earl S, Masci PP, Lavin MF. 2005. Identification and analysis of venom gland-specific genes from the coastal taipan (*Oxyuranus scutellatus*) and related species. *Cell Mol Life Sci* **62**: 2679-2693.

Rádis-Baptista G, Kubo T, Oguiura N, Svartman M, Almeida TMB, Batistic RF, Oliveira EB, Vianna-Morgante ÂM, Yamane T. 2003. Structure and chromosomal localization of the gene for crotamine, a toxin from the South American rattlesnake, *Crotalus durissus terrificus*. *Toxicon* **42**: 747-752.

Rao VS, Swarup S, Kini RM. 2003. The nonenzymatic subunit of pseutarin C, a prothrombin activator from eastern brown snake (*Pseudonaja textilis*) venom, shows structural similarity to mammalian coagulation factor V. *Blood* **102**: 1347-1354.

Reza MA, Swarup S, Kini RM. 2007. Structure of two genes encoding parallel prothrombin activators in *Tropidechis carinatus* snake: gene duplication and recruitment of factor X gene to the venom gland. *Thromb Haemost : JTH* **5**: 117-126.

Schield DR, Card DC, Hales NR, Perry BW, Pasquesi GM, Blackmon H, Adams RH, Corbin AB, Smith CF, Ramesh B et al. 2019. The origins and evolution of chromosomes, dosage compensation, and mechanisms underlying venom regulation in snakes. *Genome Res* **29**: 590-601.

Shibata H, Chijiwa T, Oda-Ueda N, Nakamura H, Yamaguchi K, Hattori S, Matsubara K, Matsuda Y, Yamashita A, Isomoto A et al. 2018. The habu genome reveals accelerated evolution of venom protein genes. *Sci Rep* **8**: 11300.

Skejić J, Steer DL, Dunstan N, Hodgson WC. 2015. Label-Free (XIC) quantification of venom procoagulant and neurotoxin expression in related Australian elapid snakes gives insight into venom toxicity evolution. *J Proteome Res* **14**: 4896-4906.

Suryamohan K, Krishnankutty SP, Guillory J, Jevit M, Schröder MS, Wu M, Kuriakose B, Mathew OK, Perumal RC, Koludarov I et al. 2020. The Indian cobra reference genome and transcriptome enables comprehensive identification of venom toxins. *Nat Genetics* **52**: 106-117.

Tasoulis T, Isbister G. 2017. A review and database of snake venom proteomes. *Toxins* **9**: 290.

Wu PF, Chang LS. 2000. Genetic organization of A chain and B chain of beta-bungarotoxin from Taiwan banded krait (*Bungarus multicinctus*). A chain genes and B chain genes do not share a common origin. *Eur J Biochem* **267**: 4668-4675.
